# Arc Regulates Transcription of Genes for Plasticity, Excitability and Alzheimer’s Disease

**DOI:** 10.1101/833988

**Authors:** How-Wing Leung, Gabriel Wei Quan Foo, Antonius M.J. VanDongen

## Abstract

The immediate-early gene Arc is a master regulator of synaptic function and a critical determinant of memory consolidation. Arc protein is localized to excitatory synapses, where it controls AMPA receptor endocytosis, and to the nucleus, where it associates with Tip60, a subunit of a chromatin modifying complex. Here we show that Arc interacts with dynamic chromatin loops and associates with histone markers for active enhancers and transcription in cultured hippocampal neurons. When Arc induction by pharmacological network activation was prevented using a short hairpin RNA, the expression profile was altered for over 1900 genes. Many gene families were affected by the absence of Arc, most notably those associated with synaptic function, neuronal plasticity, intrinsic excitability (channels, receptors, transporters), and signaling pathways (transcription factors/regulators). Interestingly, about 100 genes whose activity-dependent expression level depends on Arc are associated with the pathophysiology of Alzheimer’s disease, suggesting a critical role for Arc in the development of neurodegenerative disorders. When endogenous Arc expression was induced in a non-neuronal cell line (HEK293T), the transcription of many neuronal genes was increased, suggesting Arc can control expression in the absence of activated signaling pathways. Taken together, these data establish Arc as a master regulator of neuronal activity-dependent gene expression and a significant factor underlying the pathophysiology Alzheimer’s disease.

## INTRODUCTION

The neuronal immediate-early gene **Arc**^1,2^ plays a critical role in memory consolidation^3–6^. Arc expression is rapidly and transiently induced by novel behavioural and sensory experiences^7–11^, while its mRNA is enriched in dendrites and targeted to recently activated synapses, where it is locally translated^12,13^. Arc protein resides in excitatory synapses, where it controls AMPA receptor endocytosis^14^, allowing it to act as a master regulator of synaptic function and plasticity^15,16^ that implements homeostatic synaptic scaling at the neuronal network level^17–19^. While the synaptic role of Arc has been well documented, the observed failure to convert early-to late-LTP in Arc knockout mice cannot be explained by an AMPA receptor endocytosis deficit^4^. This suggests that Arc may have additional functions. Interestingly, Arc protein can also be localized in the nucleus, where it binds to a beta-spectrin IV isoform and associates with PML bodies^20–22^, sites of epigenetic regulation of gene transcription^23^. Nuclear Arc has been reported to regulate transcription of the GluA1 AMPA receptor^24^. Recently, another nuclear function for Arc has been demonstrated: Arc interacts with the histone-acetyltransferase Tip60^25^, a subunit of a chromatin modifying complex^26–28^. Arc expression level correlates with the acetylation status of one of Tip60’s substrate: lysine 12 of histone 4 (H4K12)^25^, a memory-associated histone mark which declines with age^29^. These newly discovered nuclear functions may point to an epigenetic role for Arc in memory consolidation. We have therefore investigated Arc’s interaction with chromatin and its association with histone marks in cultured hippocampal and cortical neurons. Fluorescent microscopy experiments demonstrated a highly dynamic interaction between chromatin and Arc, as well as a tight association between Arc and histone marks for active enhancers and active transcription. RNA-Sequencing (RNA-Seq) experiments in which activity-dependent Arc expression was prevented using a short hairpin RNA showed that Arc regulates the transcription of over nineteen hundred genes controlling memory, cognition, synaptic function, neuronal plasticity, intrinsic excitability and intracellular signaling. Interestingly, Arc also controls the expression of susceptibility genes for Alzheimer’s disease, as well as many genes implicated in the pathophysiology of this disorder. A Gene Ontology (GO) analysis identified downstream signaling pathways and diseases associated with the observed changes in mRNA levels, while an Ingenuity Pathway Analysis (IPA) revealed upstream regulators predicted by the change in gene expression profile caused by Arc knockdown. Finally, we induced expression of Arc in human embryonic kidney 293T (HEK-293T) cells, using CRISPR-Cas9, which resulted in the increased transcription of many neuronal genes. Taken together, our data demonstrate that Arc controls neuronal activity-dependent expression of many genes underlying higher brain functions and may be involved in the development of Alzheimer’s disease (AD) and other neurodegenerative disorders.

## RESULTS

Arc is a neuronal activity-dependent immediate-early gene^1,2^, whose expression is induced by exposure to a novel environment or a new sensory experience^7,8,11^. Knockdown of Arc expression abrogates long-term memory without affecting short-term memory, indicating a critical role for Arc in memory consolidation^3–6^. Arc protein localizes to dendritic spines, where it regulates AMPA receptor endocytosis^14^, and to the nucleus^20,24,30,31^, where its function is less understood. In this study we have used cultured hippocampal and cortical neurons to study the role of Arc in the nucleus. Arc expression can be induced by increasing network activity in neuronal cultures, using a combination of 4-aminopyridine (4AP), bicuculline and forskolin (**4BF**), a form of pharmacological long-term potentiation (LTP)^21,22,32,33^. **Figure 1** shows that this form of network activation strongly induces the expression of Arc in a subset of neurons. In this *in vitro* paradigm, Arc localizes predominantly to the nucleus four hours after network activity-dependent induction of its expression.

**Figure 1.**
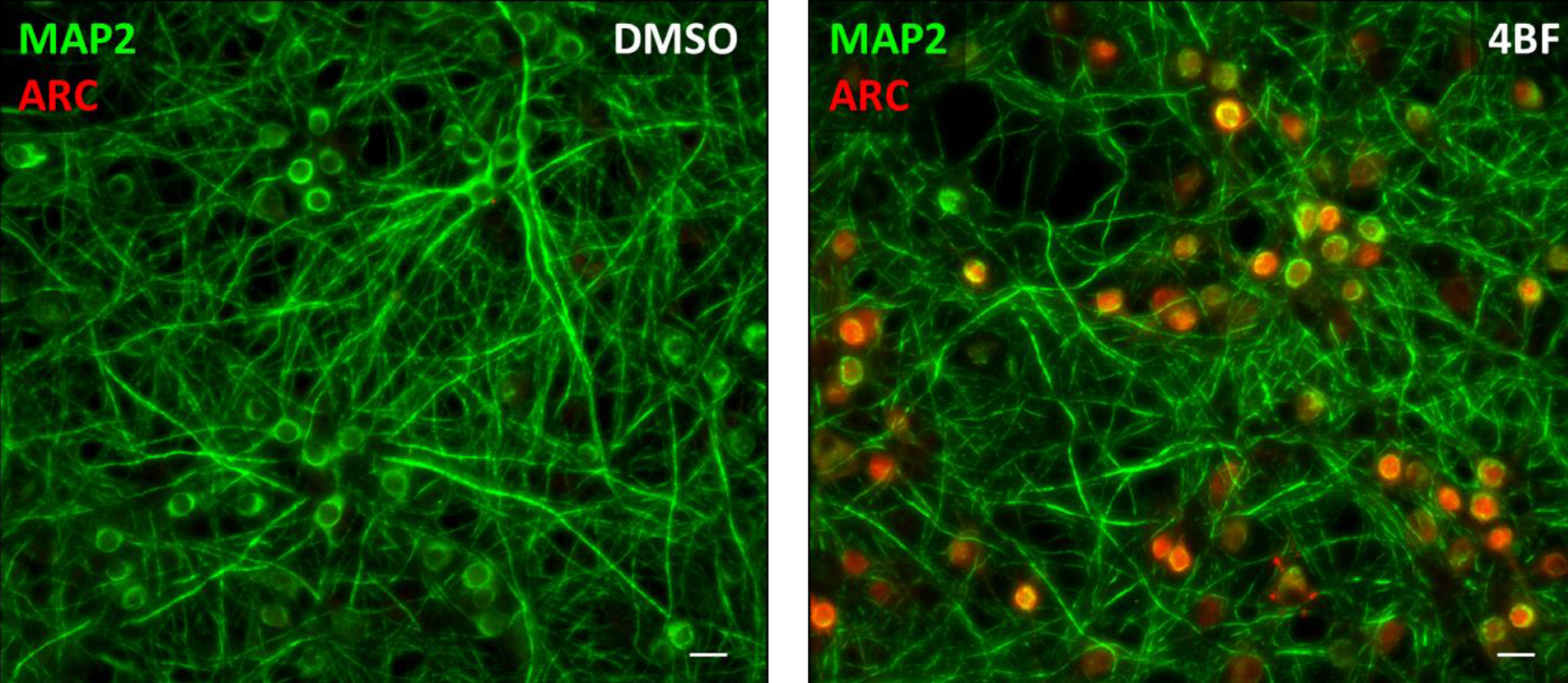
Induction of Arc expression in hippocampal neurons by pharmacological network activation. Hippocampal neurons (DIV19-21) were treated with **4BF**, which pharmacologically stimulates network activity and induces long-term potentiation of excitatory synapses. After 4 hours of enhanced network activity, neurons were fixed and stained for Arc (red) and the neuronal marker Map2 (green). Under vehicle (DMSO) treatment, very little Arc staining could be detected (left panel, Vehicle), whereas the increase in network activity induced strong nuclear Arc expression in approximately half of the neurons: 49 ± 8 % (n=3) (right panel, **4BF**).

Memory consolidation requires *de novo* gene expression^34,35^, which is induced by activation of signaling cascades that originate in the synaptic connections potentiated during learning^36–40^. This synapse-to-nucleus signaling results in post-translational modifications of chromatin, including acetylation, methylation, phosphorylation, and sumoylation of histones and methylation of DNA^41,42^. Chromatin modification alters its nanostructure, which controls accessibility of gene promoters to the transcription machinery^43,44^. These synaptic activity-induced epigenetic processes can alter gene expression and have been shown to be critical for learning and memory^45–52^. We therefore characterized the structure and dynamics of chromatin in cultured hippocampal neurons, evaluated how pharmacological LTP (**4BF** treatment) affected chromatin structure, and compared chromatin properties of neurons expressing Arc protein with control neurons that do not.

### Chromatin reorganization in Arc-positive neurons

Induction of Arc protein expression by pharmacological network activation (**Fig. 1**) is relatively slow and reaches a maximum level between 4 and 8 hours and Arc is only expressed in a subset of neurons. As shown in **Figure 2**, chromatin organization is different between neurons that are positive and negative for Arc. Chromatin was visualized by labeling DNA with the fluorescent dye 4’,6-diamidino-2-phenylindole (DAPI). Whereas chromatin in Arc-negative neurons is relatively homogenous, the nuclei of Arc-positive neurons contain many bright puncta, representing chromocenters with densely packed chromatin, in which genes are likely silenced (**Fig. 2A and 2B**). The puncta are interpersed with domains of highly open chromatin, which is more supportive of efficient gene transcription. The number of puncta increased from 11.1±0.8 puncta in Arc-negative nuclei to 15.9±0.8 puncta in Arc-positive nuclei (**Fig. 2C**). However, the mean area of the puncta was not significantly different between Arc-positive and Arc-negative neurons (**Fig. 2D**).

**Figure 2.**
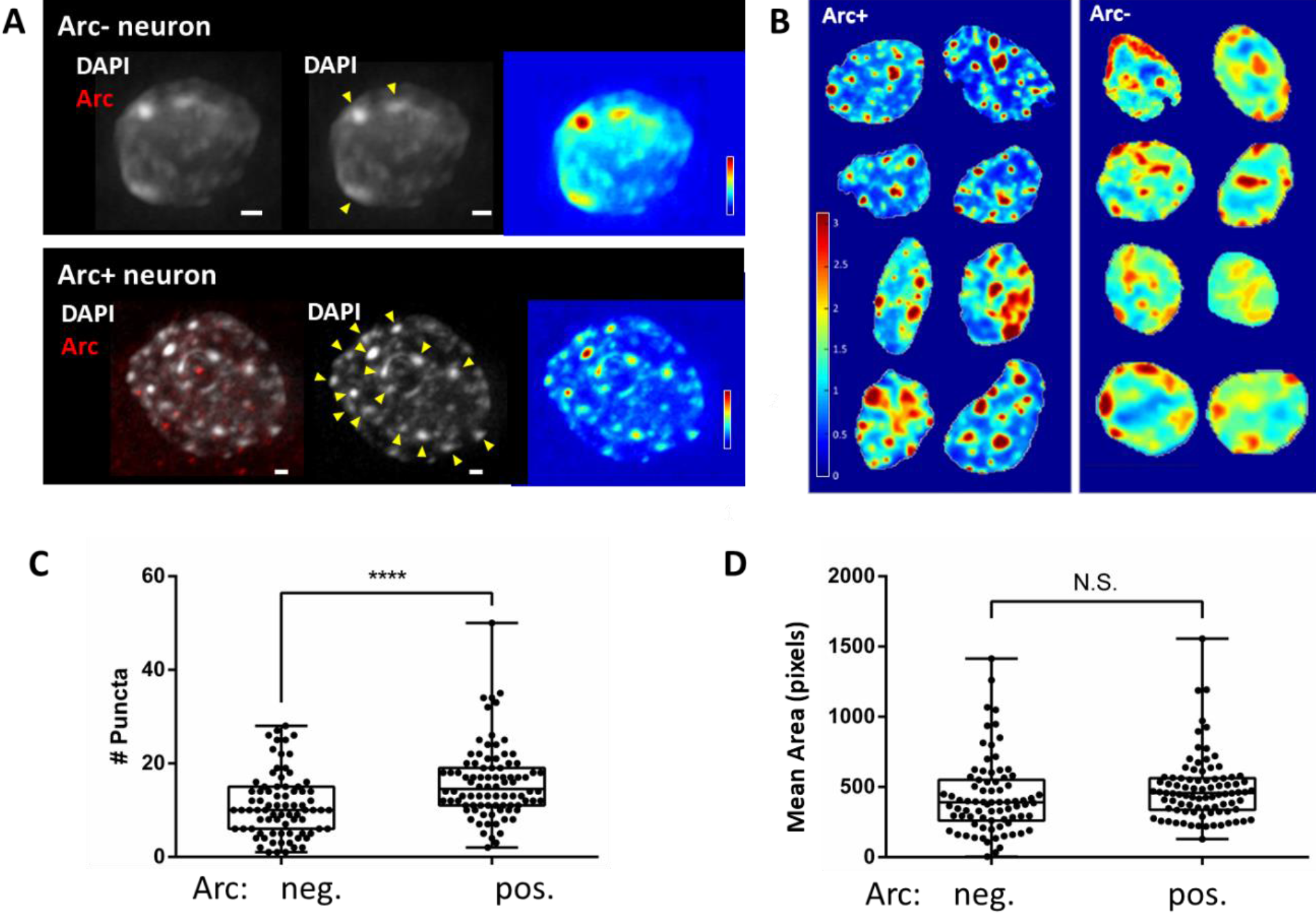
Chromatin reorganization in Arc-positive neurons. Arc expression was induced in a subset of cultured hippocampal neurons by a 4-hour treatment with **4BF**. Cells were fixed and stained for Arc (C7 antibody, Santa Cruz). DNA was labeled using DAPI. Z-stacks of DAPI images were obtained for neuronal nuclei that were positive and negative for Arc expression. Out-of-focus fluorescence was removed using 3D deconvolution (AutoQuant). **(A)** Max-projection images of a representative nucleus from an Arc-negative (top) and Arc-positive neuron (bottom). The white bar indicates a scale bar of 1 μm. DNA, labeled by DAPI, is shown in white while Arc expression is shown in red. Yellow arrowheads indicate DNA puncta. Heat maps of the relative DAPI intensity of the nucleus is shown in the rightmost panels. Relative DAPI intensity is shown using the color scale shown at the bottom. **(B)** DAPI heat maps for nuclei of 8 Arc-positive and 8 Arc-negative neurons. Relative DAPI intensity is shown by the color scale on the left, which was the same for both panels. Whereas chromatin of Arc-negative neurons (right panel) was relative homogenous (turquoise, green yellow), Arc-positive neurons (left panel) was characterized by several areas of high DAPI intensity (red), indication condensed heterochromatin (chromocenters) separating domains with decondensed euchromatin (blue). **(C) and (D)** Puncta were quantified based on their size and intensity. Arc expression, measured as mean Arc intensity, was used to correlate with the properties of the puncta, generating the boxplots. Boxplots of number of puncta **(C)** and area of puncta **(D)** for Arc-negative and Arc-positive neurons. Each • represents the **(C)** number or **(D)** mean area of puncta in a nucleus. A total of 167 nuclei were analyzed from three sets of independent experiments. **(C)** Nuclei of Arc-positive neurons have significantly higher number of puncta with 11.1 ± 0.8 puncta in Arc-negative nuclei and 15.9 ± 0.8 puncta in Arc-positive nuclei. **** indicates P-value < 0.0001, unpaired t test. **(D)** No significant change in area of puncta was observed: Arc-positive had an area of 488 ± 25 pixels, while the area of Arc-negative neurons was 431 ± 30 pixels (p = 0.15, unpaired t test). N.S. indicates not significant.

### Arc associates with dynamic chromatin

The interaction between Arc and chromatin was studied in more detail using time-lapse fluorescence microscopy of hippocampal neurons expressing Arc and histone 2B (H2B) tagged with YFP and mCherry, respectively (**Fig. 3**). Arc was induced in 18 days *in vitro* (DIV) hippocampal neurons by a 4-hour treatment with **4BF**. The time-lapse movies of Arc-eYFP and H2B-mCherry revealed a highly dynamic chromatin that constantly reorganizes on a time scale of seconds (**<u>Movie 1</u>**). Arc is concentrated in small puncta to which the chromatin can be seen to reach out with finger-like structures, which likely represent the dynamic chromatin loops described by others^53–55^.

**Figure 3.**
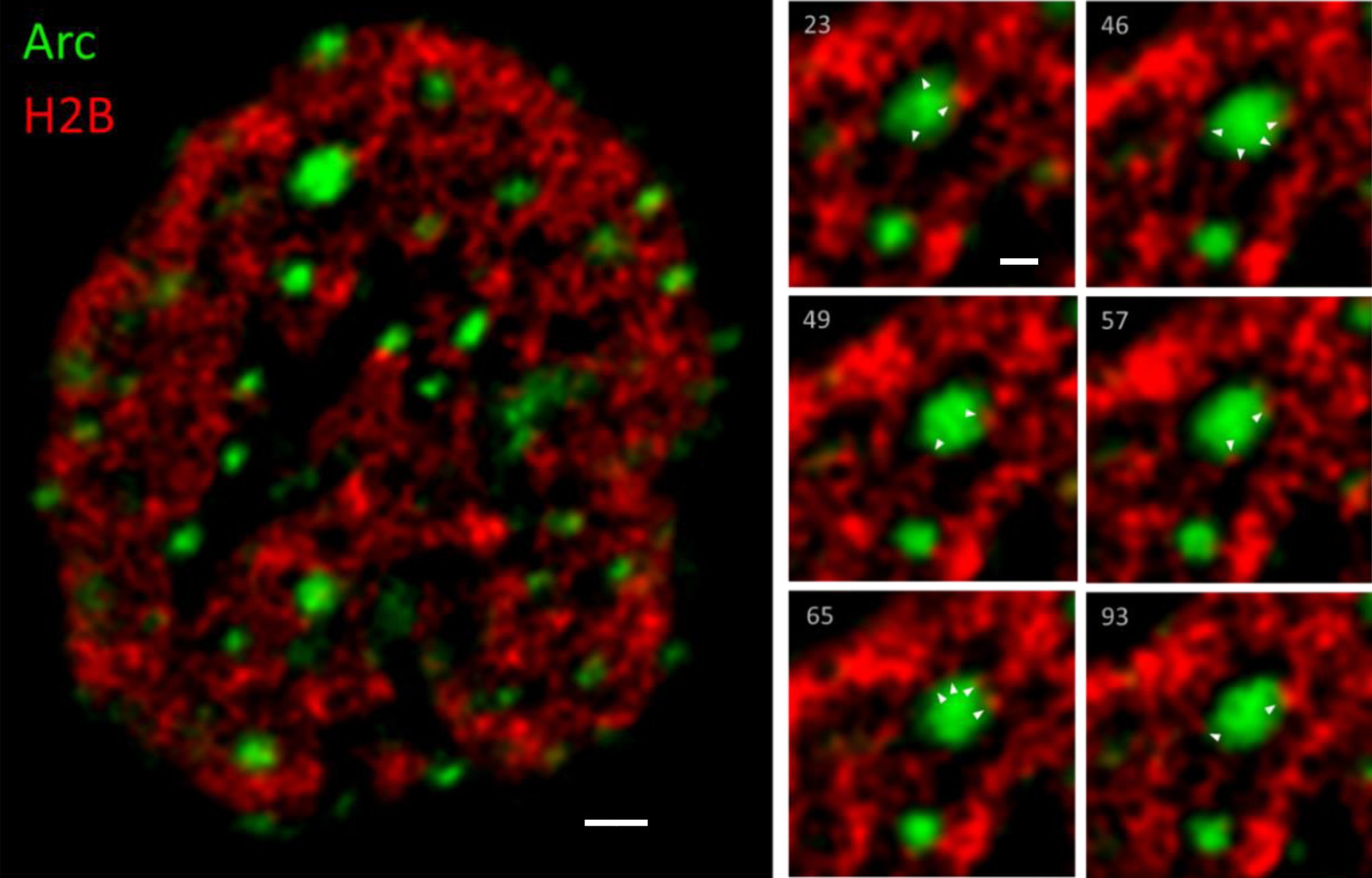
Arc associates with dynamic chromatin. Time-lapse movies of Arc-eYFP and H2B-mCherry expressed in hippocampal neurons (18 DIV) were obtained using a spinning disc confocal microscope (100x, 1.49 NA Apo TIRF objective). Z-stacks (5 images) were acquired for both YFP and mCherry channels. 3D blind deconvolution (AutoQuant) was used to remove out-of-focus fluorescence. The movie is 5 minutes long, 3.2 seconds between frames, which was the time required to acquire Z-stacks from both channels. The image on the left shows a single frame of the movie in the centre of the Z-stack of a neuronal nucleus (scale bar= 1 μm). Arc (green) is seen to form puncta, while H2B (red) labels the lattice-like chromatin structure. The panels on the right show six frames of a zoomed-in section illustrating small chromatin structures transiently interact with the two Arc puncta (scale bar = 500 nm). White arrowheads indicate points of contact between Arc and chromatin. The highly dynamic interaction of chromatin with Arc puncta is most clearly seen in the **<u>Movie</u>**.

### Arc associates with a marker of active enhancers

Because Arc was shown to associate with the Tip60 substrate H4K12Ac^25^, we have examined interactions of Arc with other histone modifications, by comparing Arc-positive and -negative neurons following pharmacological network activation. The ‘histone code’^56^ is complex and still incompletely understood. We have therefore focused on histone modifications whose function is best studied. In our survey we have found several histone modifications for which there was a difference in nuclear organization between Arc positive and negative neurons, including H3K9Ac, H3K4me3, and H3K14Ac (data not shown). **Figure 4** illustrates the close association between Arc and H3K27Ac, which marks active enhancers^57,58^. Arc and H3K27Ac form two separate lattice-like structures that are closely inter-connected and, in some locations, appear to overlap (yellow areas in **Figure 4**).

**Figure 4.**
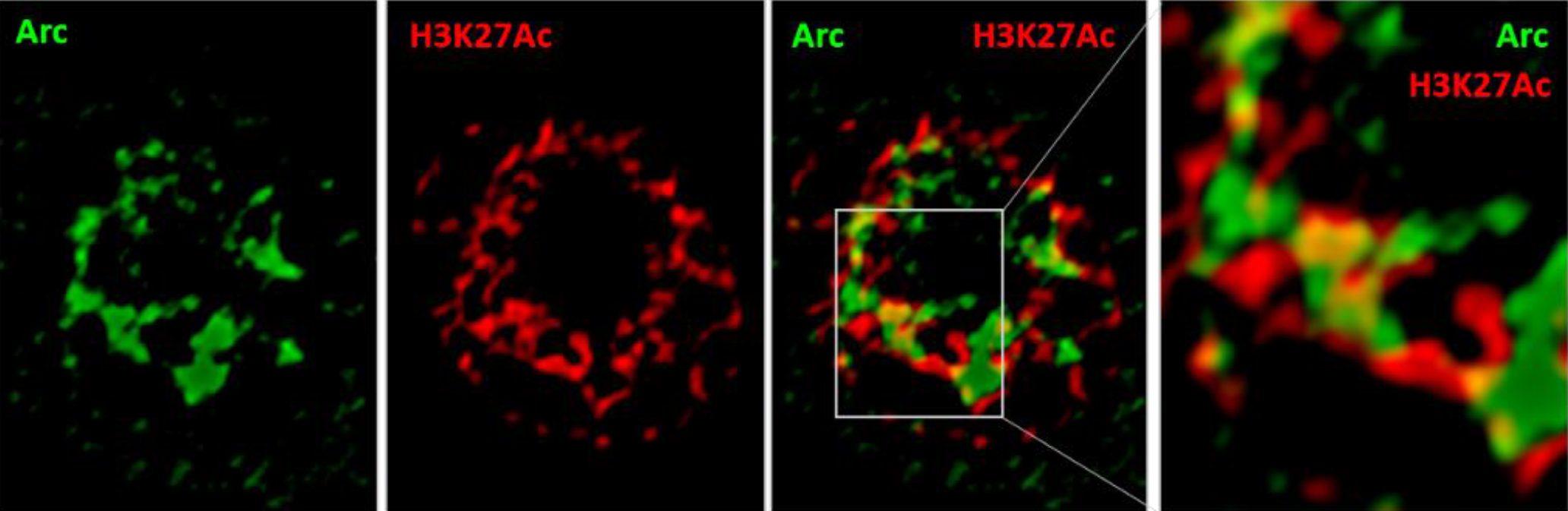
Arc associates with H3K27Ac. Hippocampal neurons were treated with **4BF** for 4 hours, fixed with methanol and stained for Arc and H3K27Ac, which marks sites containing active enhancers. Z-stacks of images were acquired of neuronal nuclei using a spinning disc confocal microscope (60X, 1.49 NA objective). Resolution was increased using 3D blind deconvolution (Autoquant). The enlarged section shows the close interaction between Arc and H3K27Ac.

### Arc associates with a marker for active transcription

Another histone mark that showed a strong interaction with Arc was H3K9Ac-S10P, which requires the concurrent acetylation of lysine 9 of histone H3 (H3K9Ac) and phosphorylation of the neighboring serine 10 (S10P). This dual marker indicates genomic regions undergoing active transcription^22,59,60^. **Figure 5** illustrates the close interaction between Arc and this histone mark, using Stochastic Optical Reconstruction Microscopy (STORM), a form of super-resolution microscopy with a resolution of ~30 nm^61^. Both Arc and H3K9Ac-S10P are enriched at the nuclear periphery, where reorganization of chromatin between active and inactive transcriptional states takes place^62,63^. With the increased resolution of STORM, Arc can be seen to localize to distinct puncta. H3K9Ac-S10P forms an elaborate meshwork, as expected for chromatin, but also is enriched in puncta-like domains. Arrowheads in **Figure 5A** indicate the close apposition between these two sets of puncta. Close inspection of the interface between the two types of puncta revealed invasions of H3K9Ac-S10P into the Arc puncta (arrows in **Fig. 5B**), resembling the finger-like chromatin loop structures seen in live cell imaging (**Fig. 3**,**<u>Movie</u>**).

**Figure 5.**
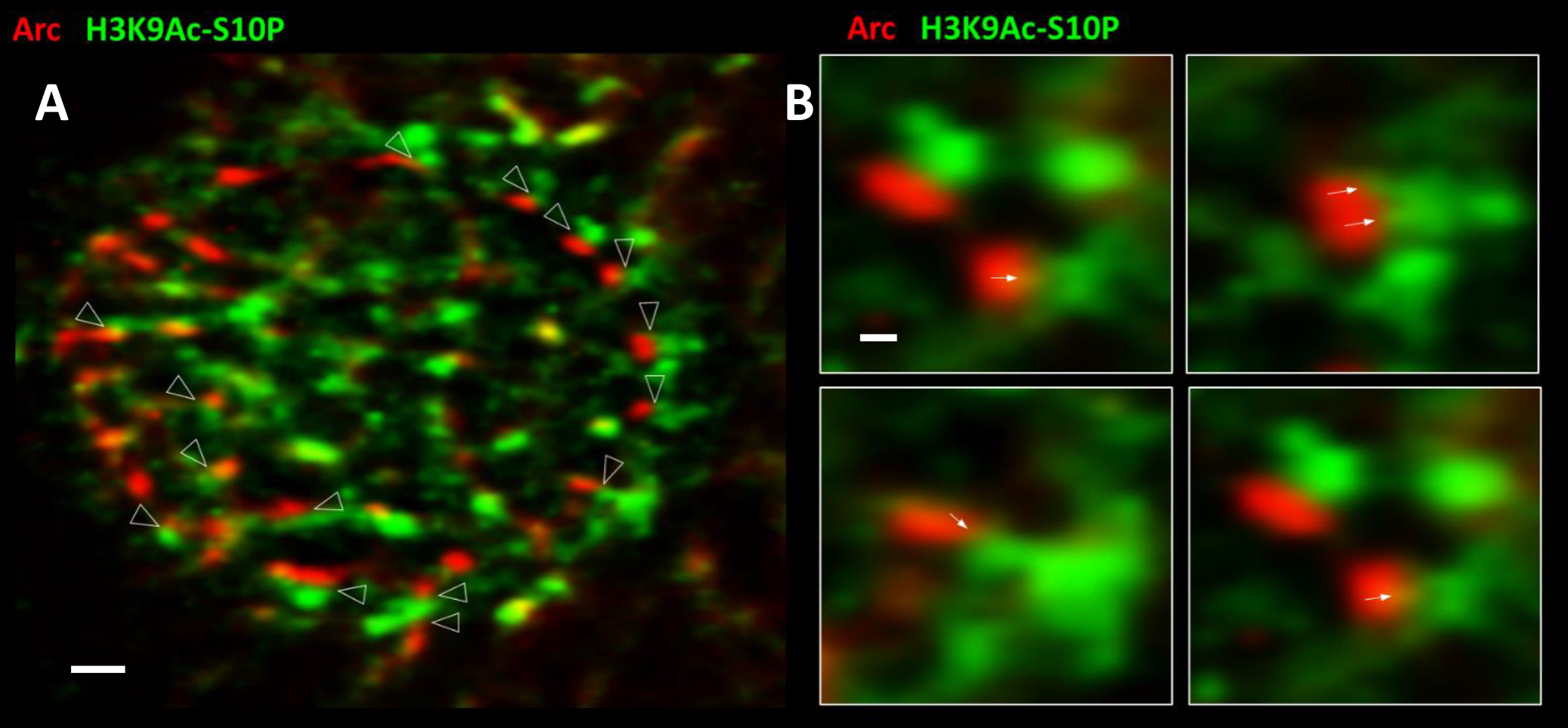
Arc associates with H3K9Ac-S10P. Image of a neuronal nucleus obtained using STORM. Cultured hippocampal neurons were treated with **4BF** for 4 hours, fixed and stained for Arc and H3K9Ac-S10P, which marks sites undergoing active transcription. **(A)** Arrowheads point to close appositions between Arc and the histone mark (scale bar = 1μm). **(B)** Enlarged sections showing the association in greater detail. Arrows point to what appear to be invasions of H3K9Ac-S10P into Arc puncta (scale bar 200 nm).

### Arc regulates activity-dependent gene transcription

Because Arc was found to associate with histone marks involved in transcription activation, we wanted to investigate whether network activity-induced Arc expression alters the gene expression profile of the neurons. Four short hairpin RNAs (shRNAs) targeting the coding region of Arc were tested for their ability to suppress Arc induction following four hours of 4BF treatment. We selected the most effective shRNA to generate an adeno-associated AAV9 virus. Because AAV9 infection itself may alter the gene expression profile, we also generated a negative control consisting of AAV9 virus encoding a scrambled version of the Arc shRNA. We performed an RNA-Seq analysis of cortical neurons expressing either the Arc shRNA or its scrambled control. When **4BF**-mediated Arc expression is prevented using the Arc shRNA (**Fig. 6A**), mRNA levels for more than 1900 genes were altered (**Fig. 6B**). Many gene families were affected, including those associated with plasticity genes (*Jun, Fosb, Bdnf, Dlg4, Egr4, Npas4* and *Nr4a1*), synaptic proteins (syntaxin *Stx12* and synaptotagmin *Syt3*), neurotransmitter receptors (NMDA, AMPA, GABA, glycine, serotonin and metabotropic glutamate receptors) (**Fig. 6B**). Arc also regulated the expression of genes controlling intrinsic excitability: 62 genes encoding ion channels (20 K^+^, 4 Na^+^, and 9 Ca^2+^ channel subunits, 7 transient receptor potential (Trp) channels, 14 ligand-gated ion channels, 7 regulatory subunits and 1 non-selective cation channel), and 139 genes encoding transporters/pumps (for glutamate, GABA, serotonin, ADP, ATP, phosphate, glucose, inositol, alanine, cysteine, glutamine, glycine, proline, Na^+^, Ca^2+^, Cl^-^, H^+^ and Zn^2+^). These results suggest that Arc regulates activity-dependent gene expression relevant for synaptic function, neuronal plasticity and intrinsic excitability.

**Figure 6.**
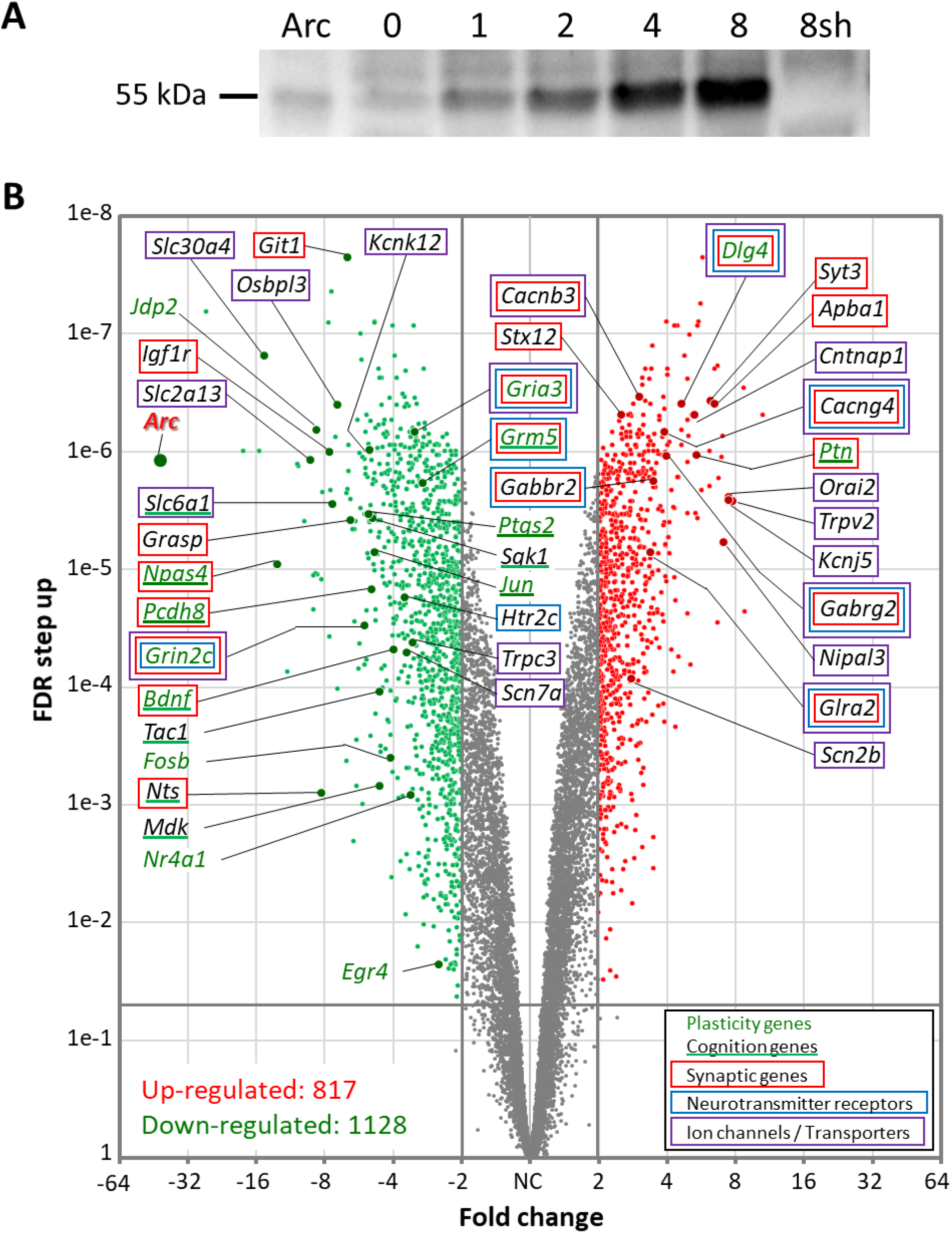
Arc regulates gene transcription. **(A)** Western Blot showing the time course of Arc protein expression in cultured hippocampal neurons following **4BF** treatment (time in hours indicated on top). Lane 7 (8sh) shows that Arc fails to express at 8 hours of **4BF** when the cultures are transduced with an AAV9 virus encoding a short-hairpin RNA (shRNA) targeting the coding region of Arc. Lane 1 has purified Arc protein. **(B)** Volcano plot of RNA-Seq results comparing mRNA isolated from neurons after 8 hours of **4BF** that were transduced with AAV9 virus encoding either the Arc shRNA or a scrambled version of this shRNA, done in triplicate. Preventing activity-dependent Arc expression resulted in the upregulation of 817 genes (red), and down-regulation of 1128 genes (green). Genes that are below the cut-off (FDR > 0.05 or absolute fold change < 2) are marked in gray. Some of the highly regulated genes involved in learning and memory are indicated in the plot. Genes are color coded as stated in the legend. Both axes are log scaled.

**Table 1** shows the 30 top-ranking genes sorted by absolute fold change (FC) caused by the shRNA-mediated knockdown of Arc expression. Gene names are shown together with a description of their function, their Fold Change, False Discovery Rate (FDR), and references to relevant papers. Many of the top-regulated genes are involved in synapse modulation, neurotransmission, neurogenesis and neurological disorders. Interestingly, 9 out of the top 30 genes have been implicated in the pathophysiology of AD (*Fgf1, Slc30a4, Npas4, Cxcl1, Jdp2, Nts, Mmp10, Orai2* and *Tomm34*) while an additional 5 genes are linked to amyloid beta (Aβ) metabolism (*Mmp13, Mmp12, Slc2a13, Igf1r* and *Apba1*).

**Table 1.**
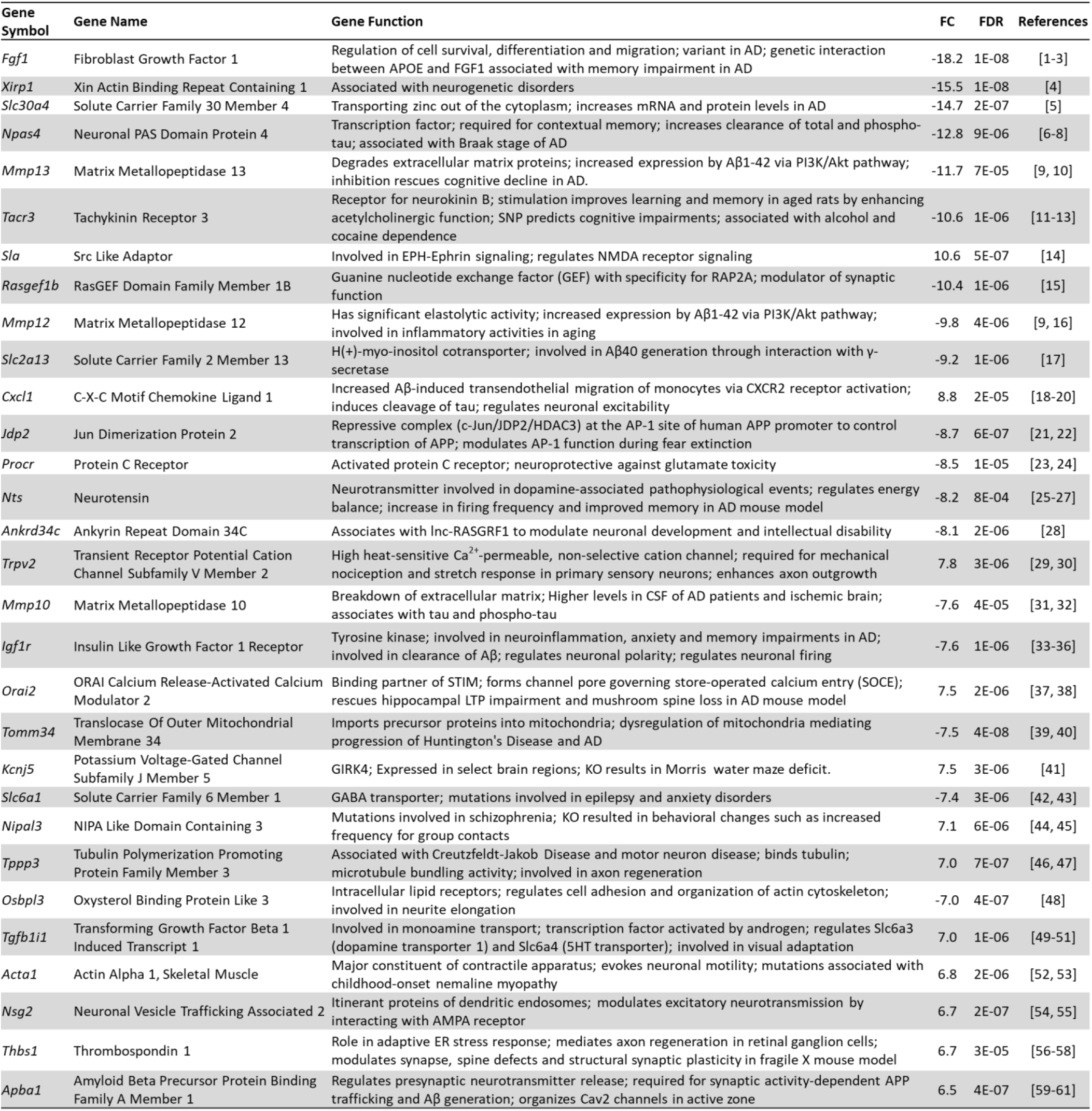
Top ranking genes with neuronal functions. APOE: apolipoprotein E; p-tau: phosphorylated tau; Aβ: Amyloid beta; PI3K/Akt: phosphatidylinositol 3-kinase/protein kinase B; CSCR2: C-X-C motif chemokine receptor 2; HDAC3: histone deacetylase 3; AP-1: activator protein 1; APP: amyloid precursor protein; CSF: cerebrospinal fluid; STIM: stromal interaction molecule; LTP: long term potentiation; GABA: γ-aminobutyric acid; KO: knockout; 5HT: 5-hydroxytryptamine; ER: endoplasmic reticulum; Cav2: neuronal voltage-gated calcium channels. **References:** [1-3]^232–234^, [4]^235^, [5]^236^, [6-8]^237–239^, [9, 10]^240,241^, [11-13]^242–244^, [14]^245^, [15]^246^, [9, 16]^240,247^, [17]^248^, [18-20]^249–251^, [21, 22]^252,253^, [23, 24]^254,255^, [25-27]^256–258^, [28]^259^, [29, 30]^260,261^, [31, 32]^262,263^, [33-36]^264–267^, [37, 38]^268,269^, [39, 40]^270,271^, [41]^272^, [42, 43]^273,274^, [44, 45]^275,276^, [46, 47]^277,278^, [48]^279^, [49-51]^280–282^, [52, 53]^283,284^, [54, 55]^285,286^, [56-58]^287–289^ and [59-61]^290–292^.

### GO analysis of differentially expressed genes

A Gene Ontology (GO) analysis of the RNA-Seq data identified several biological processes and molecular functions that were affected when Arc expression was prevented during network activation (**Fig. 7**). Arc knockdown altered many genes involved in the regulation of nervous system development and neuronal differentiation (**Fig. 7A**). In addition, many of the genes were enriched in biological processes involved in cognition, regulation of cell projection organization and axonogenesis (**Fig. 7A**), processes which could modulate structural plasticity involved in neural development, learning and memory^64,65^. While the top ten regulated genes enriched for the regulation of plasma membrane bounded cell projection organization were both up- and down-regulated (**Fig. 7Cii**), genes enriched for cognition and the regulation of axonogenesis were mostly down-regulated due to the absence of Arc (**Fig. 7Ci** and **iii**). Many of the altered genes were also enriched in molecular functions such as ion channel regulator activity, glutamate receptor binding and ligand-gated ion channel activity (**Fig. 7B**), including *Sgk1* (**Fig. 7Di**), *Dlg4*, which encodes PSD-95 (**Fig. 7Dii**), and *Grin2c*, which encodes the NMDA receptor NR2C subunit(**Fig. 7Diii**). These molecular functions are well-established to underlie synaptic plasticity processes crucial for formation of memory^66,67^.

**Figure 7.**
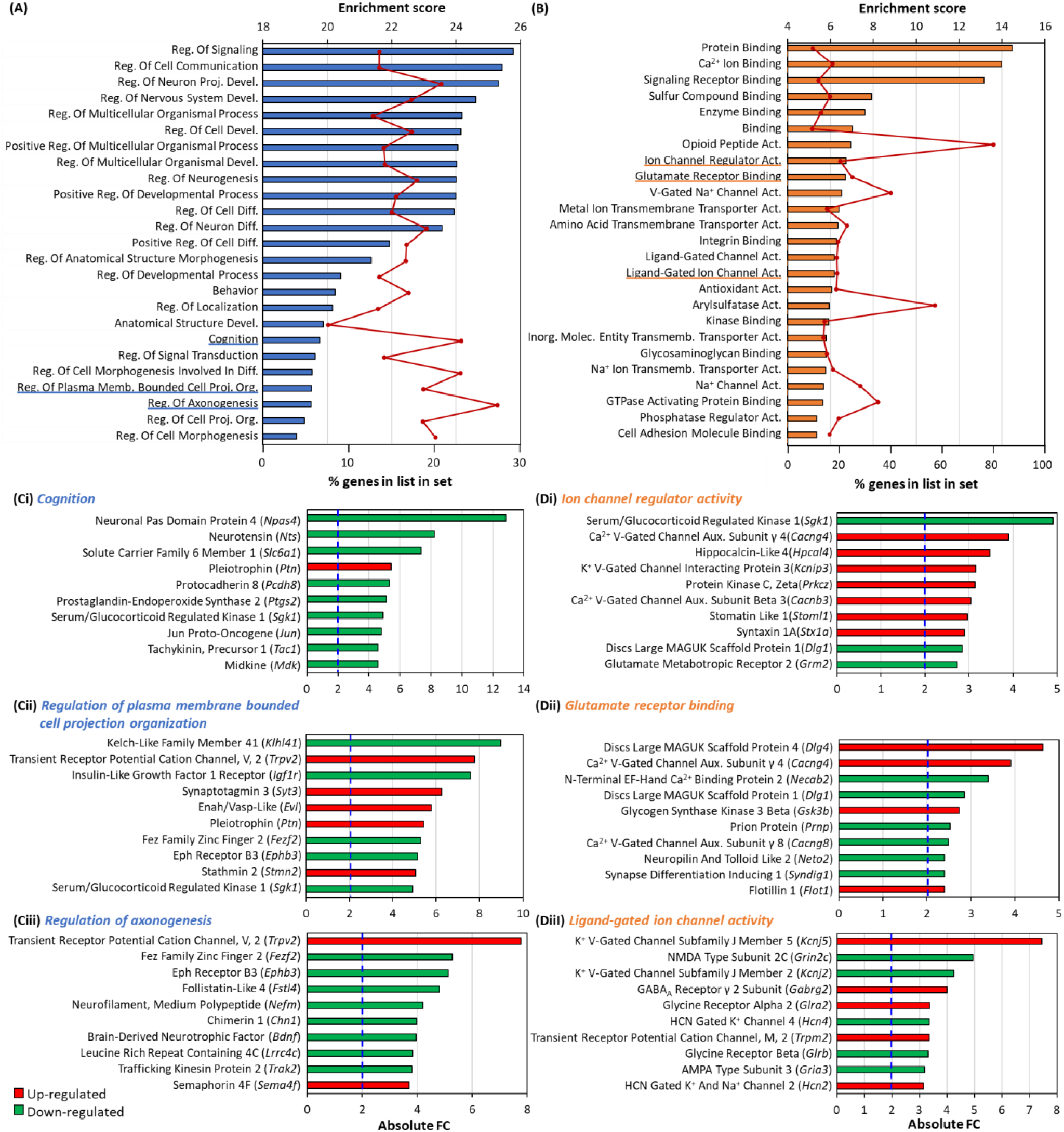
GO analysis of Arc knock-down. **(A)** and **(B)** Gene set enrichment analysis was performed to investigate the Biological Processes **(A)** and Molecular Functions **(B)** that the altered genes were involved in. The enrichment score is plotted against the category names. The enrichment score is the negative natural logarithm of the enrichment P-value derived from Fisher’s exact test and reflects the degree to which the gene sets are overrepresented at the top or bottom of the entire ranked list of genes. Bars indicate the enrichment score while the line graph indicates the percentage of genes that are altered under the respective GO term. The top 25 biological processes **(B)** and molecular functions **(C)** are shown. Many of the categories are related to synaptic plasticity (underlined blue and orange). **(C)** and **(D)** Bar-charts showing genes involved in the stated category from Biological Processes **(Ci)**-**(Ciii)** and Molecular Functions **(Di)**-**(Diii)** and their respective fold changes. The top 10 regulated genes are shown. Dotted blue line indicates an absolute Fold Change of 2.

### Arc regulates expression of synaptic and plasticity genes

From the GO results in **Figure 7**, we observed that the knockdown of Arc affected many genes involved in synaptic plasticity, as well as genes implicated in processes underlying learning and memory. We have therefore investigated how Arc knockdown affected genes encoding synaptic proteins by manually curating a list of differentially expressed genes whose protein products are located at the presynaptic or postsynaptic compartment. A total of 232 synaptic genes were differentially expressed. These genes are involved in the development and growth of axons and dendrites, including *Ephb3*, *Lrfn2*, *Lama5*, *Neurod2, Sema4f, Caprin2*, and *Unc5c* and the modulation of the synapses and dendritic spines, including *Npas4*, *Pcdh8, Ephb3, Lrfn2, Bdnf, Atxn1*, *Cbln2*, *Cadps2*, *Caprin2*, *C1ql1*, *C1ql3*, and *Unc5c* (**Table 2**). Many of these synaptic genes are also involved in neuroplasticity, cognition, learning and memory, including *Syt3, Pcdh8, Pdyn, Lrfn2, Dlg4, Kcna4, Bdnf and Mapki8ip2*. **Figure 8** lists neuroplasticity genes and genes that are involved in cognition, learning and memory whose activity-dependent expression is regulated by Arc. Most of these genes were down regulated when activity-dependent Arc expression was prevented.

**Figure 8.**
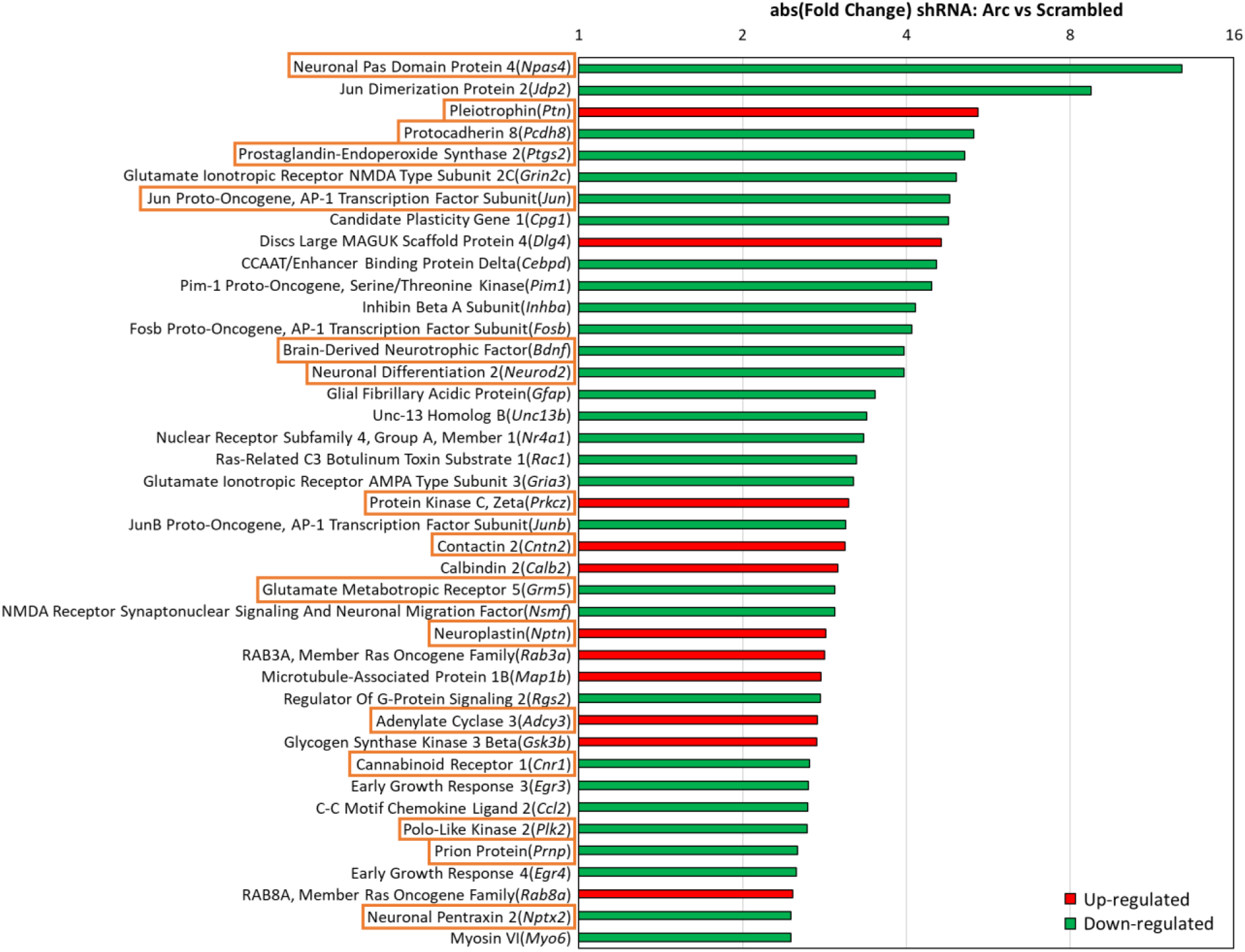
Neuronal plasticity genes regulated by Arc. Neuronal plasticity genes were manually curated in addition to reference to GO terminology from the Gene Ontology Consortium. Neuronal plasticity genes with absolute FC ? 2.5 are shown. Genes that are involved in cognition or learning and memory are marked by orange boxes.

**Table 2.**
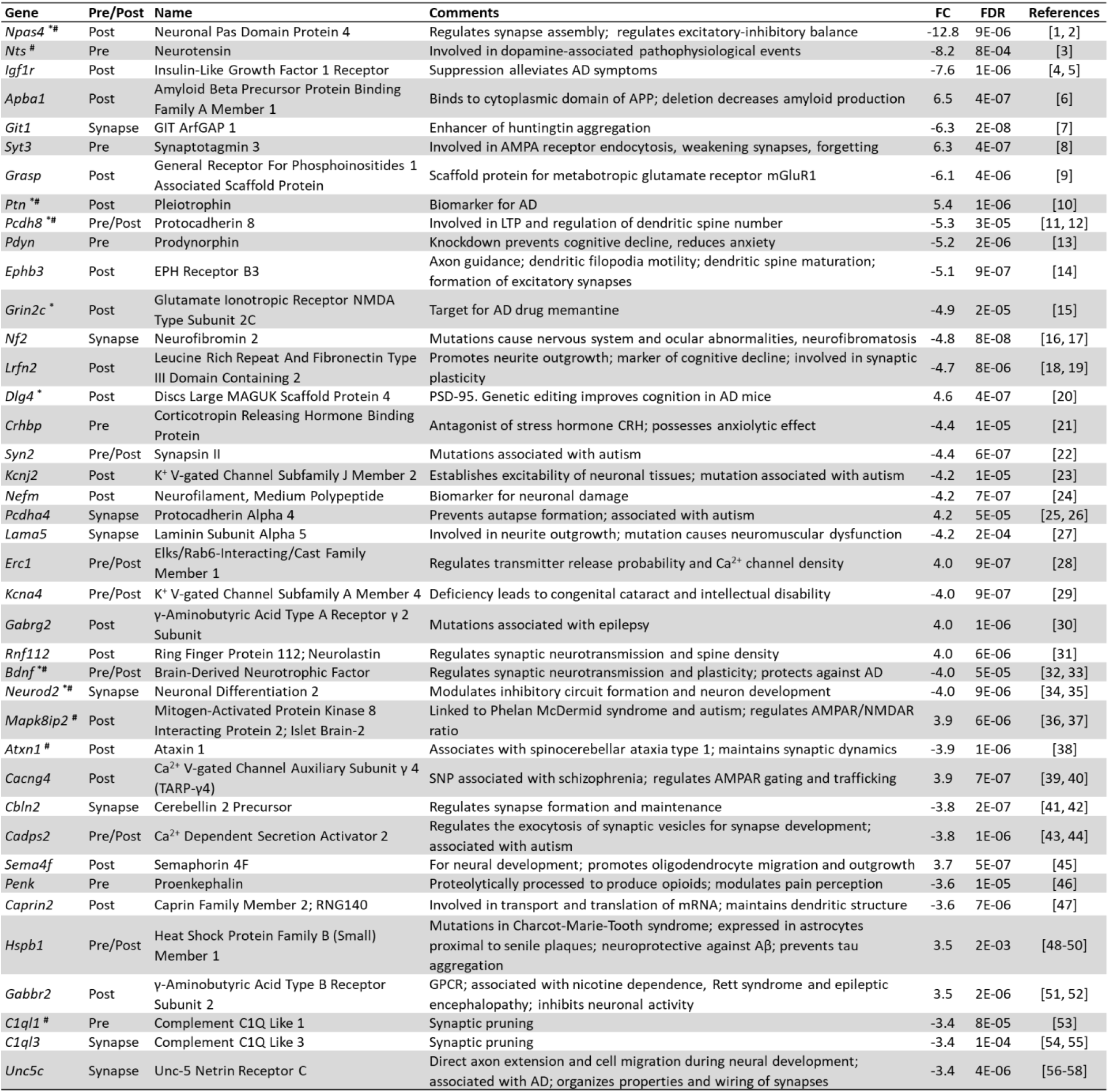
Synaptic genes whose expression is regulated by Arc. This table shows the top 40 synaptic genes (out of a total of 323) ranked by absolute fold change. Comments list relevant information about the function and disease association of the genes. An asterisk (*) indicates genes involved in neuroplasticity. A hashtag (#) indicates genes involved in cognition, learning and memory. APP: amyloid precursor protein; LTP: long-term potentiation; CRH: corticotropin-releasing hormone; V-gated: voltage-gated; NMDAR: *N*-methyl-D-aspartate receptor; TARP-γ4: transmembrane AMPR regulator protein γ4; Aβ: amyloid beta; GPCR: G-protein-coupled receptor. **References:** [1,2]^293,294^, [3]^295^; [4, 5]^264,265^, [6]^292^, [7]^296^, [8]^297^, [9]^298^, [10]^299^, [11, 12]^300,301^, [13]^302^, [14]^303^, [15]^304^, [16, 17]^305,306^, [18, 19]^307,308^, [20]^309^, [21]^310^, [22]^311^, [23]^312^, [24]^313^, [25, 26]^314,315^, [27]^316^, [28]^317^, [29]^318^, [30]^319^, [31]^320^, [32, 33]^321,322^, [34, 35]^323,324^, [36, 37]^325,326^, [38]^327^, [39, 40]^328,329^, [41, 42]^330,331^, [43, 44]^332,333^, [45]^334^, [46]^335^, [47]^336^, [48-50]^337^’^339^, [51, 52]^340,341^, [53]^342^, [54, 55]^343,344^ and [56-58]^345^’^347^.

### Arc knockdown altered synaptogenesis, synaptic plasticity and neuroinflammation pathways

From the GO results and the list of manually curated synaptic genes, we were interested in investigating the signaling pathways and the possible downstream effects resulting from Arc knockdown. We have analyzed the differentially expressed genes and their respective fold changes using IPA. **Figure 9A** shows the top 15 pathways that were altered due to Arc knockdown. IPA made inferences on the activation or inhibition of the pathways based on the differential expression observed and canonical information stored in the Ingenuity Knowledge Base. The degree of activation or inhibition of each identified pathway is indicated by the z-score. The ratio is calculated as the number of differential expressed genes for each pathway divided by the total number of genes involved in that pathway. Many identified pathways involved cellular signaling cascades, including those mediated by CDK5, PTEN, integrin and corticotropin releasing hormone (**Fig. 9A**). Pathways predicted to be responsible for the observed differential expression profile include opioid and endocannabinoid signaling, synaptogenesis, synaptic long-term depression (LTD) and neuroinflammation (**Fig. 9A** and **9B**). *Kcnj5*, *Ptgs2*, *Grin2c*, *Cacng4* and *Gnaq* are members of at least two of the pathways shown and are synaptic genes or associated with cognition (**Fig. 7, 8, 9A, Table 2**). Except for the neuroinflammation signaling pathway, all these pathways are associated with synaptic plasticity. Knockdown of Arc modulated neurotransmission, synaptic plasticity, spine formation/maintenance and neurite outgrowth, processes that are crucial for learning and memory (**Fig. 9B**)^68–70^. Interestingly, the two hallmarks of AD, the generation, clearance and accumulation of amyloid beta (Aβ) and the formation of neurofibrillary tangles (NFTs), are both affected by downregulation of the neuroinflammation signaling pathway resulting from Arc knockdown (**Fig. 9B**). These alterations in the generation and clearance of molecular markers and triggers of AD could indicate a possible role of Arc in the pathophysiology of AD^71^.

**Figure 9.**
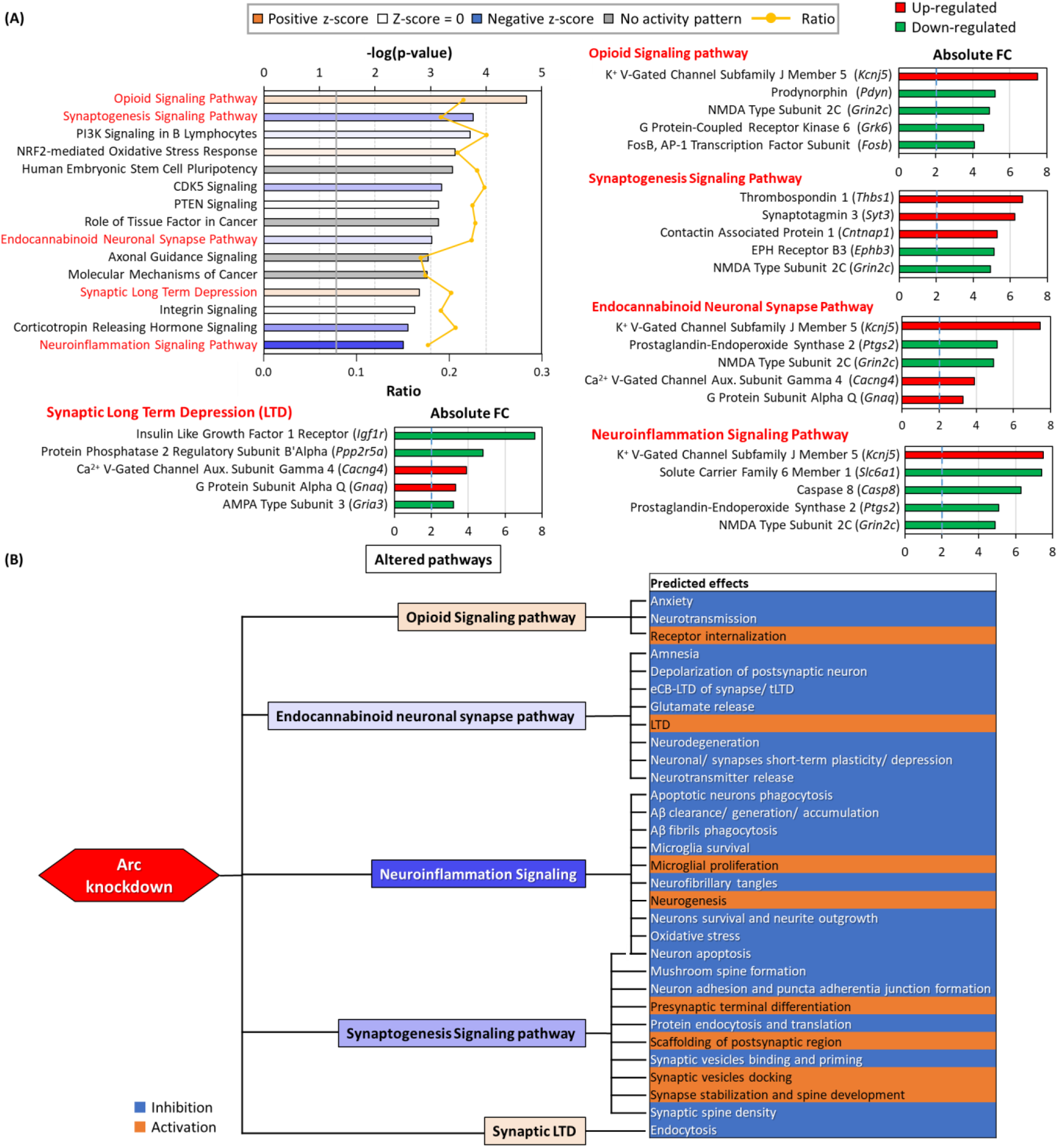
Pathways altered by Arc knockdown and predicted effects. **(A)** Bar-chart showing altered pathways identified after analysis with IPA. The orange line graph indicates the ratio of genes in our dataset that were involved in the specified pathway. The gray line indicates threshold at P-value 0.05. The orange and blue bars indicate predicted activation and inhibition of pathways respectively (determined by z-score). Top 15 significantly altered pathways are shown. Pathways with predicted activation or inhibition of downstream effects are in red, further elaborated in panel B. The top 5 genes altered in the respective pathways are shown on the right and bottom of the altered pathway bar-chart. **(B)** Diagram describing the predicted effects of the altered pathways. Five pathways are highlighted, and its downstream effects as predicted by IPA are listed.

### Arc knockdown changes the expression of genes involved in the aetiology and pathophysiology of AD

Considering that the generation, clearance and accumulation of amyloid beta and neurofibrillary tangles was predicted to be altered due to the knockdown of Arc, we investigated whether any neurological diseases or psychological disorders were correlated with the profile of differentially expressed genes mediated by Arc knockdown. **Table 3** summarized the disease annotation and predicted activation state for two disease/disorder classes whose associated genes were significantly altered by Arc knockdown. Absence of Arc was predicted to increase damage of the cerebral cortex and its neurons and cells. In addition, Arc knockdown was also associated with psychological disorders, including Huntington’s Disease, basal ganglia disorder, central nervous system (CNS) amyloidosis, tauopathy and Alzheimer disease. Of note, CNS amyloidosis and tauopathy are predictors of AD. The activation states of the five psychological disorders were not reported, possibly due to inconsistencies in the literature findings with respect to fold changes of the differentially expressed genes. However, the p-values for all five disorders were highly significant, suggesting that the progression of these disorders may be modulated by the Arc function.

**Table 3.** Prevention of activity-dependent expression of Arc resulted in gene expression profile changes that are associated with neurological diseases and psychological disorders, including Alzheimer’s disease.

| Disease and disorders | Diseases Annotation | p-Value | Predicted Activation State | Activation z-score |
| --- | --- | --- | --- | --- |
| Neurological Diseases | Damage of cerebral cortex cells | 4.4E-03 | Increased | 2.8 |
|  | Damage of cerebral cortex | 6.9E-04 | Increased | 2.6 |
|  | Damage of cortical neurons | 1.2E-02 | Increased | 2.4 |
|  | Neurodegeneration of inner hair cells | 3.3E-03 | Increased | 2.1 |
| Psychological Disorders | Huntington's Disease | 9.1E-09 |  |  |
|  | Disorder of basal ganglia | 8.2E-07 |  |  |
|  | Central nervous system amyloidosis | 2.5E-05 |  |  |
|  | Tauopathy | 3.2E-05 |  |  |
|  | Alzheimer disease | 4.2E-05 |  |  |

We next investigated how Arc knockdown could affect genes that were previously identified to increase susceptibility to AD. We have manually curated genes that were found to be genetic risk factors of AD and validated them by referencing the Genome Wide Associations Studies (GWAS) catalogue^72^. Notably, critical genetic risk factors of AD such as *Picalm*, *Apoe*, *Slc24a4*, and *Clu* were downregulated upon the knockdown of Arc^73–77^ (**Fig. 10**). Out of a total of 39 susceptibility genes identified, 26 were regulated by Arc (**Fig. 10)**.

**Figure 10.**
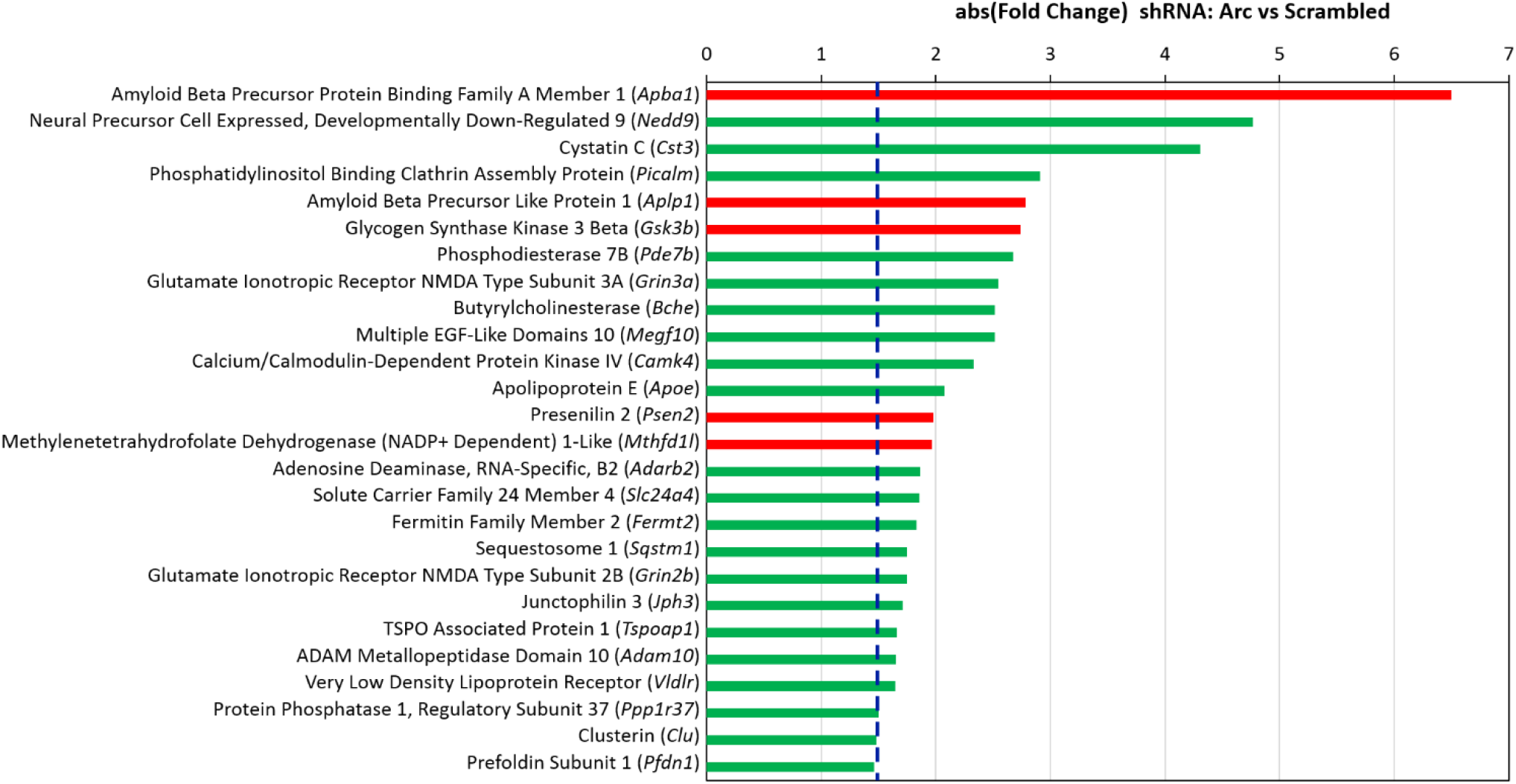
Alzheimer’s susceptibility genes affected by Arc knockdown. The expression levels of 26 AD susceptibility genes were affected when activity-dependent Arc expression was prevented by an shRNA. Green bars indicated that the mRNA level was downregulated, while red bars indicate upregulation. The blue line indicates an absolute fold change of 1.5.

Because Arc plays a role in the aetiology of AD by modulating its genetic risk factors, we investigated if Arc regulates genes that are more broadly involved in the pathophysiology of AD. **Table 4** lists the results. While some differentially expressed genes control amyloid beta formation/accumulation through the regulation of cleavage and stabilization of amyloid precursor protein (APP) (*Mmp13*, *Slc2a13*, *Apba1*, *Casp8, Ptgs2*, *Gpr3, Pawr, Timp3, Kcnip3, Plk2, Aplp2, Bace2, Apoe* and *Apba2*), others are involved in the hyperphosphorylation of tau and formation of neurofibrillary tangles (*Npas4*, *Cxcl1*, *Dryrk2*, *Tril*, *Pltp, Plk2* and *Selenop*). Arc knockdown also altered the expression of genes that are associated with the neurodegeneration and neurotoxicity observed in AD (*Casp8*, *Bcl2l11*, *Alg2*, *Tac1, Bdnf, Hmox1, Pawr, Ccl2, Selenop* and *Atf6*). Finally, Arc regulated genes associated with altered cognitive function, a characteristic of AD (*Mmp13, Pdyn, Tac1*, *Bdnf*, *Nr4a2*, *Penk*, *Pltp* and *Ccl2*). To date, presenilin 1 (*Psen1*) and glycogen synthase kinase 3 beta (*Gsk3b*) are the only AD mediators which have been reported to physically associate and interact with Arc^78–80^. Arc also interacts with endophilin 2/3 and dynamin and recruits them to early/recycling endosomes to traffic APP and beta secretase 1 (BACE1), crucial determinants of AD progression^80^. However, the observation that knocking down Arc resulted in more than 100 differentially expressed genes that are either AD susceptibility genes or genes implicated in the pathophysiology of AD (**Fig. 10** and **Table 4**), indicated that Arc could be mediating the expression of these genes via transcriptional regulation and not simply physical interactions. Arc has previously been reported to reside in the nucleus^20,24,30,31^ and we have shown how Arc physically associates with chromatin and with markers of active transcription and enhancers (**Fig. 3**, **4** and **5**). Therefore, we wanted to investigate how Arc downregulation affects transcription regulation.

**Table 4.**
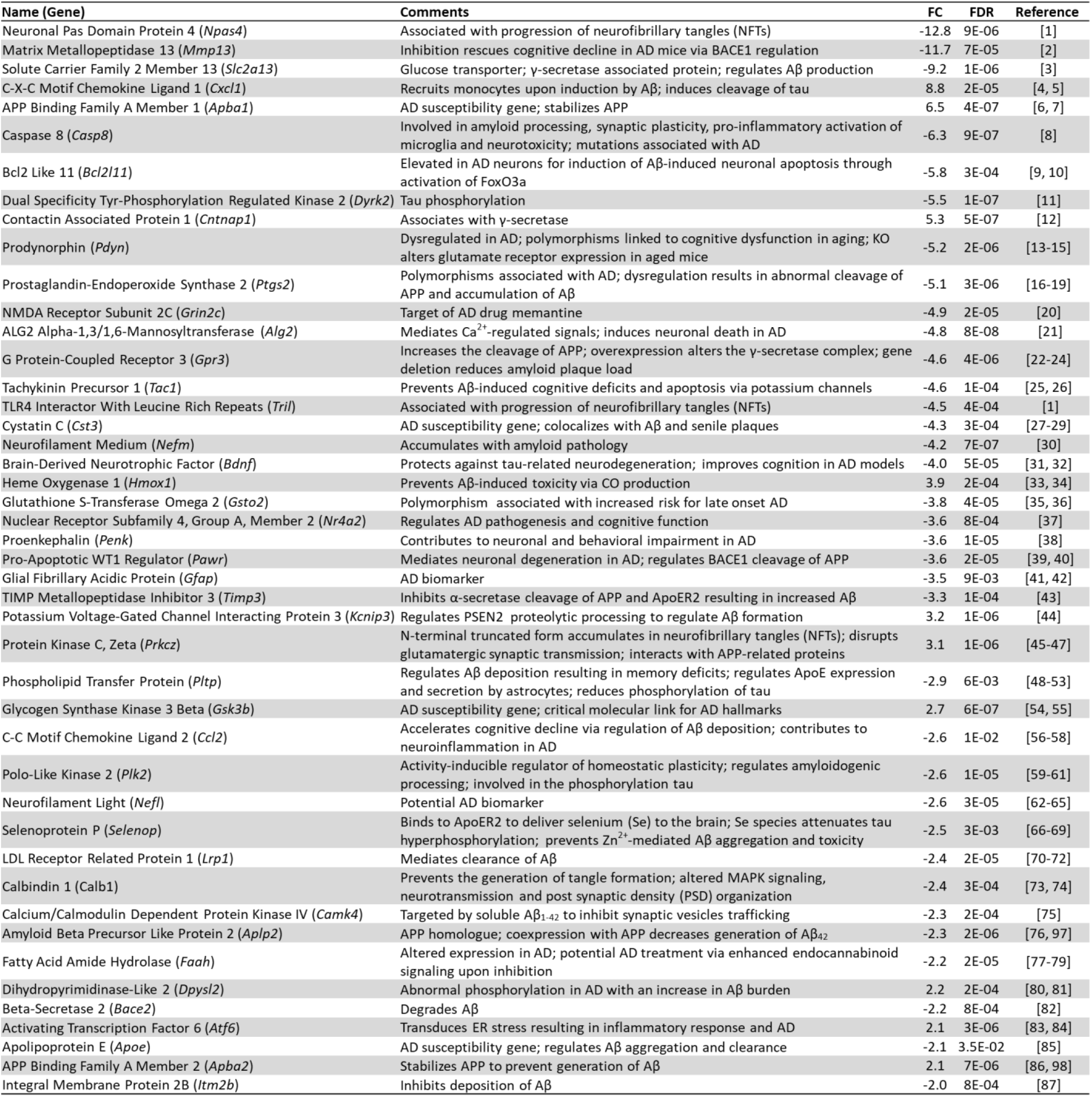
Alzheimer’s genes regulated by Arc. Only genes with more than one citation or citations that included mechanisms of regulating AD are presented. BACE1: β-secretase 1; Aβ: amyloid beta; APP: amyloid precursor protein; FoxO3a: forkhead box O3; KO: knock-out; CO: carbon monoxide; ApoER2: apolipoprotein E receptor 2; PSEN2: presenilin 2; Zn^2+^: zinc; MAPK: mitogen-activated protein kinase; ER: endoplasmic reticulum. **References:** [1]^238^, [2]^241^, [3]^248^, [4, 5]^250,251^, [6, 7]^292,348^, [8]^349^, [9, 10]^350,351^, [11]^352^, [12]^353^, [13-15]^354–356^, [1619]^85,357–359^, [20]^304^, [21]^360^, [22-24]^361–363^, [25, 26]^364,365^, [1]^238^, [27-29]^366–368^, [30]^264^, [31, 32]^321,322^, [33, 34]^369,370^, [35, 36]^371,372^, [37]^373^, [38]^374^, [39, 40]^375,376^, [41, 42]^377,378^, [43]^379^, [44]^380^, [45-47]^381–383^, [48-53]^384–389^, [54, 55]^390,391^, [56-58]^392–394^, [59-61]^395–397^, [62-65]^398–401^, [66-69]^402–405^, [70-72]^406–408^, [73, 74]^409,410^, [75]^411^, [76]^412^, [77-79]^413–415^, [80, 81]^416,417^, [82]^418^, [83, 84]^419,420^, [85]^421^, [86]^422^ and [87]^423^.

### Arc regulates the expression of transcription factors

From our GO analysis and a manual curation based on literature citations, we have identified 369 transcriptional regulators and transcription factors whose expression is controlled by Arc. **Table 5** shows the top 40 transcriptional regulators or factors whose mRNA levels were altered when activity-dependent Arc expression was prevented. Some of the transcriptional regulators are involved in neuronal development and differentiation (*Fgf1*, *Tgfb1i1, Fezf2*, *Jun*, *Magel2, Neurod2, Atxn1, Gdf15, Prdm1, Mycn, Nr4a2 and Pou2f2*), while others are involved in the development of neurological or neurodegenerative diseases (*Npas4, Igf1r*, *Txnip*, *Lgr4*, *Cebpd*, *Pim1, Magel2, Ireb2, Smad7, Sorbs1, Nfil3, Pknox2, Hdac9, Hmox1, Atxn1, Cbfb, Lrp2, Hipk3* and *Nr4a2*). Many of the transcriptional regulators/factors have been implicated in memory formation and plasticity, such as such as *Thbs1*, *Jun, Tet3, Fosb*, *Atxn1* and *Cbfb*. A GO analysis by DAVID^81^ was carried out to identify the biological processes that these transcription factors could be modulating. **Table 6** shows the top 20 biological processes that were regulated by altered transcription factor expression and that have neurological relevance. Corroborating the identified functions of the top 40 transcriptional regulators/factors (**Table 5**), differentially expressed transcriptional regulators/factors were observed to be highly enriched in biological processes such as differentiation of neurons, nervous system development, learning, long-term memory and aging (**Table 6**). Some of the transcriptional regulators were involved in multiple processes: *Npas4*, *Jun*, *Bdnf*, *Nr4a2* and *Elavl4* modulate learning, long-term memory, aging, neuron differentiation and nervous system development (**Table 6**).

**Table 5.**
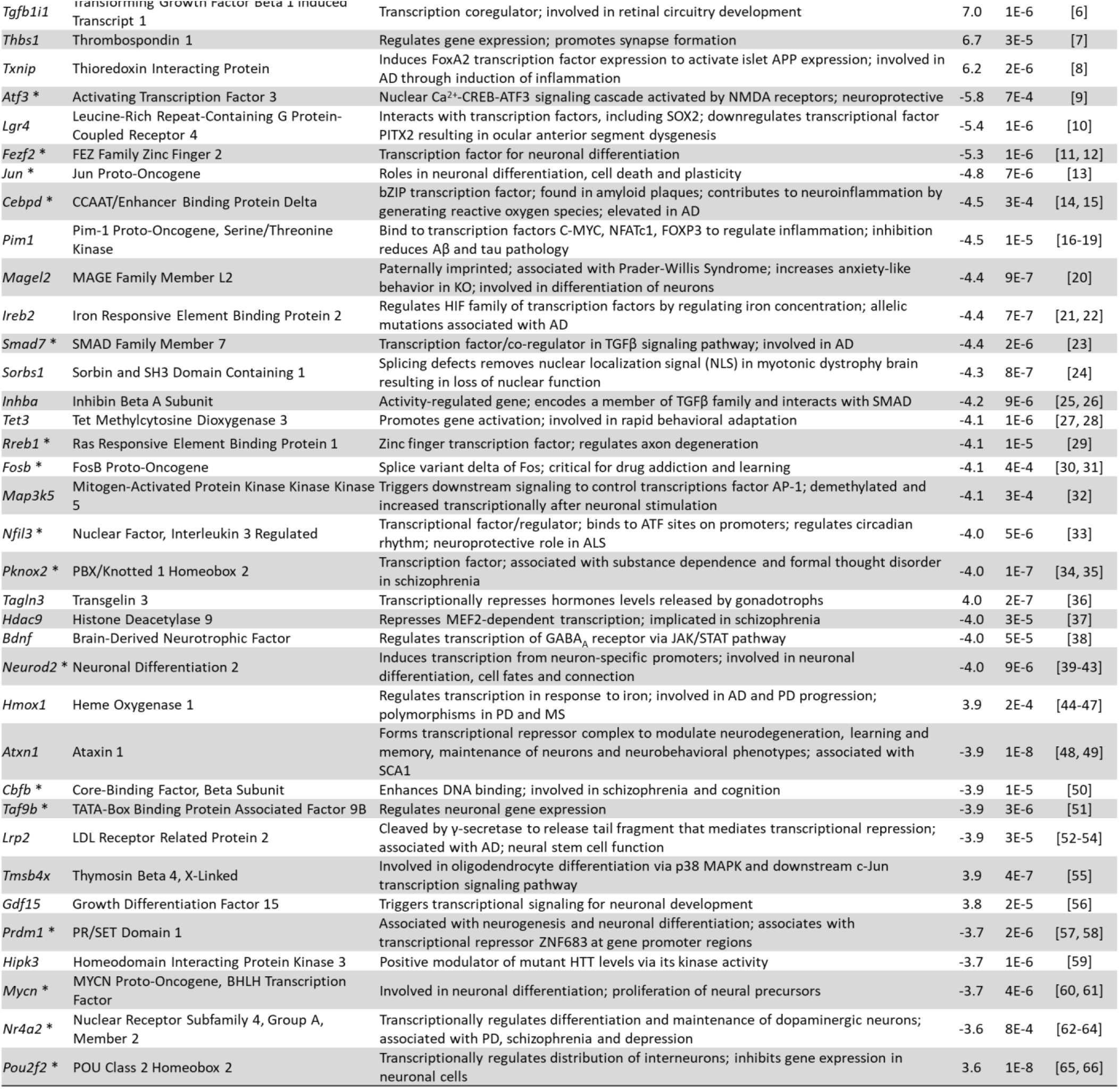
Genes involved in transcriptional regulation. This table shows the top 40 genes with neuronal relevance. * indicates transcription factors. ATF3: activating transcription factor 3; LEF1: lymphoid enhancer binding factor 1; FoxA2: forkhead box A2; APP: amyloid precursor protein; CREB: cAMP response element-binding protein; NMDAR: N-methyl-D-aspartate receptor; SOX2: SRY-box 2; PITX2: paired like homeodomain 2; bZIP: basic leucine zipper domain; C-MYC: MYC proto-oncogene, BHLH transcription factor; NFATc1: nuclear factor of activated T cells 1; FOXP3: forkhead box P3; HIF: hypoxia inducible factor; TGFβ: transforming growth factor beta; PD: Parkinson’s disease; NLS: nuclear localization signal; SMAD: transcription factors forming the core of the TGFβ signaling pathway; AP-1: activator protein 1; ALS: amyotrophic lateral sclerosis; MEF2: monocyte enhancer factor; GABAA: γ-aminobutyric acid type A; JAK/STAT: Janus kinases/ signal transducer and activator of transcription; SCA1: spinocerebellar ataxia type 1; MAPK: mitogen-activated protein kinase; ZNF683: zinc finger protein 683; HTT: huntingtin. **References::** [1]^424^, [2]^425^, [3]^426^, [4, 5]^264,427^, [6]^282^, [7]^428^, [8]^429^, [9]^430^, [10]^431^, [11, 12]^432,433^, [13]^434^, [14, 15]^435,436^, [16-19]^437–440^, [20]^441^, [21, 22]^442,443^, [23]^444^, [24]^445^, [25, 26]^446,447^, [27, 28]^448,449^, [29]^450^, [30, 31]^451,452^, [32]^453^, [33]^454^, [34, 35]^455,456^, [36]^457^, [37]^458^, [38] ^459^, [39-43]^323,324,460–462^, [44-47]^463–466^, [48, 49]^467,468^, [50]^469^, [51]^470^, [52-54]^471–473^, [55]^474^, [56]^475^, [57, 58]^476,477^, [59]^478^, [60, 61]^479,480^, [62-64]^481–483^ and [65, 66]^484,485^.

**Table 6.**
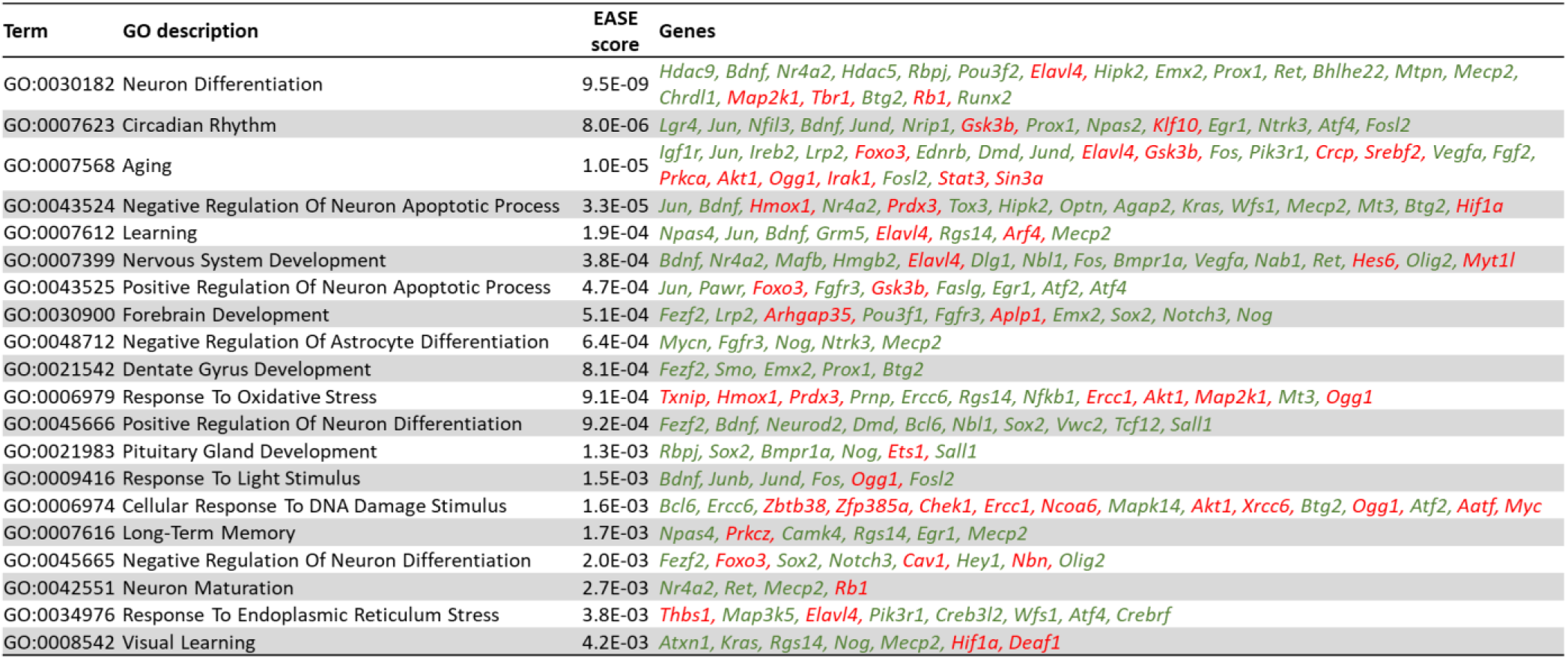
Biological processes that the transcriptional regulators/factors control. The top 20 biological processes of neurological relevance are listed. EASE score is a modified fisher exact p-value measuring the gene-enrichment in the annotated terms. Genes are arranged in order of highest to lowest absolute fold change. Genes highlighted in red are up-regulated while those highlighted in green are down-regulated.

### Upstream regulators associated with Arc-dependent genes

Because Arc knock down resulted in the differential expression of 1945 genes (**Fig. 6**), altering downstream pathways (**Fig. 9**) possibly leading to disease states (**Table 3**), we identified the upstream modulators that could explain the vast differential expression pattern observed. From the IPA analysis, 11 upstream regulators were predicted to critically contribute to the differential expression profile (**Table 7**). Except for *Sox2*, none of these upstream regulators were transcriptionally affected by Arc knockdown, suggesting Arc controls their function through a different mechanism. SOX2 and HDAC4 were both activated by the absence of Arc, while the function of the remaining 9 regulators was inhibited. Of note, the predicted inhibition of CREB1 (z-score = −3.5) and APP (z-score = −2.8) explains the differential expression of 100 and 94 genes, respectively (**Table 7**). The 11 upstream regulators predicted by IPA control the expression of *Nr4a2*, *Slc6a1* and *Igf1r*, genes that are also involved in AD progression, neuroinflammation pathways and synaptic LTD (**Fig. 9A** and **Table 4**). We have investigated the mechanisms by which Arc could alter the function of the identified upstream regulators resulting in the alteration of downstream pathways and AD progression. The downstream pathways investigated are (i) opioid signaling, (ii) synaptogenesis signaling, (iii) the endocannabinoid neuronal synapse pathway, (iv) synaptic LTD and (v) neuroinflammation signaling (**Fig 11)**. These are also the pathways whose downstream effects we focused on in **Figure 9**. APP, CREB1 and TNF are three upstream regulators identified by IPA that controlled the highest number of genes involved in the downstream pathways highlighted (**Fig. 11**). The top five genes regulated by APP were *Igf1r* (synaptic LTD)^82^, *Ptgs2* (endocannabinoid neuronal synapse pathway; neuroinflammation, AD progression)^83–86^, *Jun* (neuroinflammation, AD progression)^87–90^, *Dlg4* (PSD95, synaptogenesis)^91,92^ and *Syn2*(synaptogenesis)^93^ (**Fig. 11**). In addition to *Ptgs2* and *Syn2*, CREB1 regulated the differential expression of *Slc6a1* (neuroinflammation)^94^, *Pdyn* (opioid signaling)^95^ and *Fosb* (opioid signaling)^96^ (**Fig. 11**). Interestingly, TNF, whose transcription was not altered upon knockdown of Arc, regulates 15 genes (**Table 7**), the top five of which are *Casp8* (neuroinflammation)^97^, *Ptgs2* (also regulated by APP and CREB1), *Gabrg2* (neuroinflammation)^98,99^, *Bdnf* (synaptogenesis; neuroinflammation, AD progression)^100–103^ and *Penk* (opioid signaling, AD progression)^104^. While the top CREB1-regulated genes are mainly associated with the opioid signaling pathway, APP and TNF are implicated in neuroinflammation. Triggering of the neuroinflammation pathway leads to the altered expression of AD-associated genes such as *Ptgs2*, *Jun, Bdnf, Hmox1* and *Gabbr2*.

**Figure 11.**
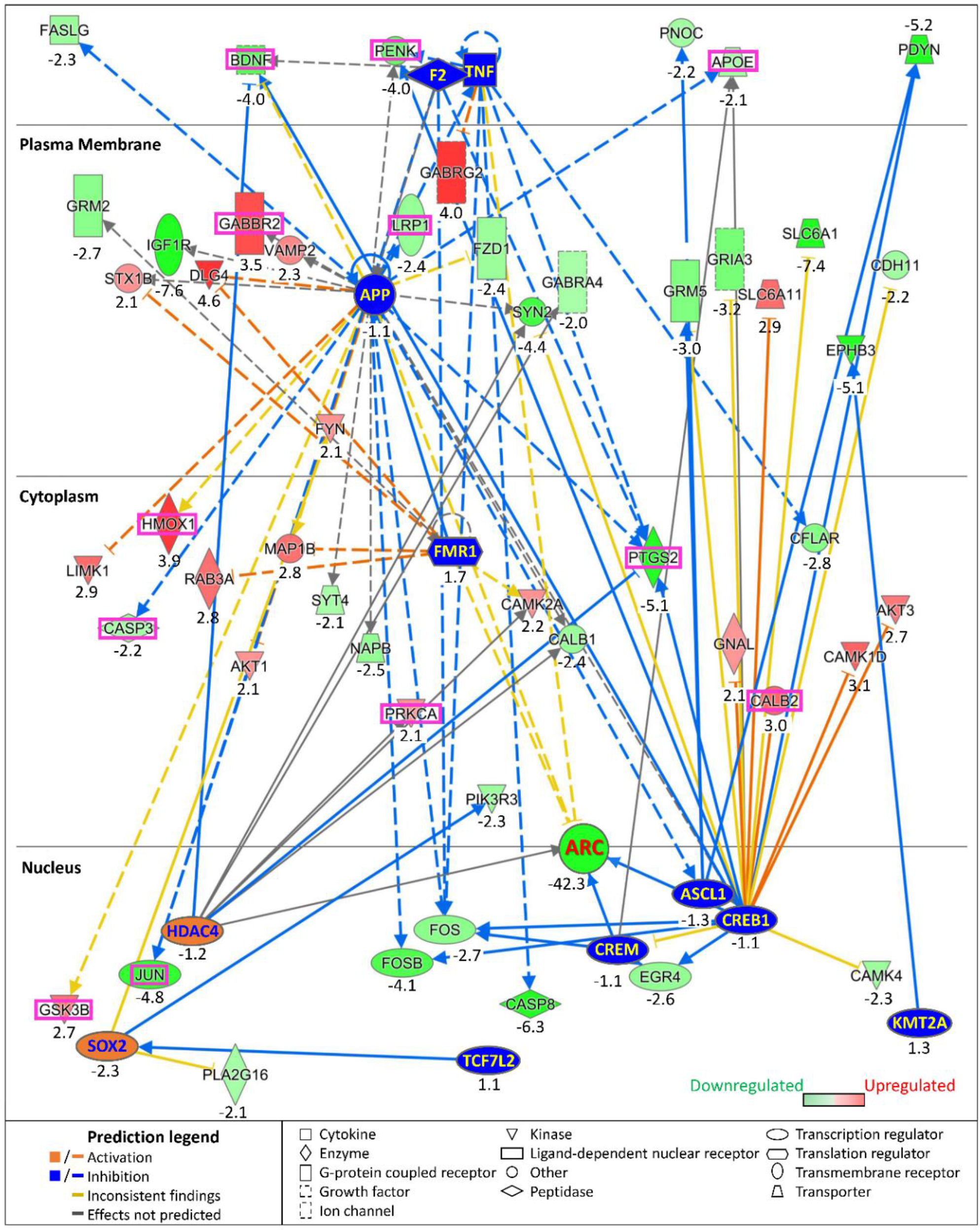
Upstream regulators of differential gene expression caused by Arc knockdown. The map shows 11 upstream regulators (blue and orange boxes) predicted by IPA to mediate altered gene expression upon Arc knockdown. Genes are positioned in the extracellular space, the plasma membrane, the cytosol or the nucleus, depending on where their associated proteins are located. Arc was positioned at the interface of nucleus and cytoplasm because it can be in either compartment. Only genes that were involved in the following pathways and disease annotations are shown: (i) opioid signaling, (ii) synaptogenesis, (iii) endocannabinoid neuronal synapse pathway, (iv) synaptic LTD, (v) neuroinflammation, (vi) CNS amyloidosis, (vii) tauopathy and (viii) AD. Genes associated with disease annotations are boxed in magenta. The respective fold changes are indicated below each gene.

### Arc over-expression alters gene expression in human embryonic kidney cells

The results presented thus far suggest that preventing Arc expression during neuronal network activation results in an altered gene expression profile affecting synaptic plasticity and cellular excitability, as well as neurodegenerative disease state. We therefore tested whether Arc can alter gene transcription, outside the context of neuronal network activation and without viral infection. We induced the expression of the endogenous Arc gene in human embryonic kidney (HEK293T) cells using a CRISPR-Cas9 approach^105^ (**Fig. 12A**). Whereas wildtype HEK293T cells expressed Arc at a very low level, targeting a transcription activator complex to its promoter increased Arc mRNA levels nearly 250-fold. This in turn altered the expression of 57 genes (absolute FC > 2, p < 0.05), with 54 genes up-regulated and 3 genes down-regulated. Many of the genes have neuronal functions (**Fig. 12B**). We have performed a GO analysis to understand the cellular components (**Fig. 12C**) and biological processes (**Fig. 12D**) these differentially expressed genes were involved in. We observed many genes that are typically expressed in neurons or are synaptic components, as indicated by the following GO terms: i) *synapse part* (p = 1.1E-04), ii) *presynapse* (p = 1.0E-03), iii) *neuron part* (p = 1.5E-03) and iv) *postsynaptic membrane* (p = 2.3E-03) (**Fig. 12C**). Differentially expressed genes upon the induction of Arc in HEK293T cells are involved in synaptic transmission processes or neuronal development, including i) *chemical synaptic transmission* (p = 2.5E-04), ii) *signal release from synapse* (p = 1.9E-03), iii) *interneuron precursor migration* (p = 3.2E-03) and iv) *axon guidance* (p = 3.2E-03) (**Fig. 12D**). Genes that are associated with these cellular components and processes were also highly altered, including i) *Chat* (p=4.7E-85, choline acetyltransferase) located at presynaptic terminals, synthesizes acetylcholine, ii) *Oprd1* (p=2.6E-62, δ-opioid receptor), activation reduces pain and improves negative emotional states, iii) *Arx* (p=1.1E-70, Aristaless Related Homeobox), a transcription factor involved in neuronal migration and development, iv) *Scn1b* (p=6.6E-22, Na channel β1 subunit), involved in axonal guidance, v) *Foxa3* (p=3.3E-24, Forkhead Box A3), a transcription factor involved in the determination of neuronal fate^106,107^, vi) *Pllp* (p=1.5e-25, Plasmolipin), involved in membrane organization and ion transport, vii) *Slc18a3* (p=1.6E-16), a vesicular acetylcholine transporter at the presynapse, viii) *Fndc11* (p=4.4E-14, Fibronectin Type III Domain Containing 11), a vesicular gene, and ix) *Adgrb1* (p=3.7E-12, Adhesion G Protein-Coupled Receptor B1), localized at the postsynapse, involved in synapse organization and cell projection morphogenesis (**Fig. 12B**).

**Figure 12.**
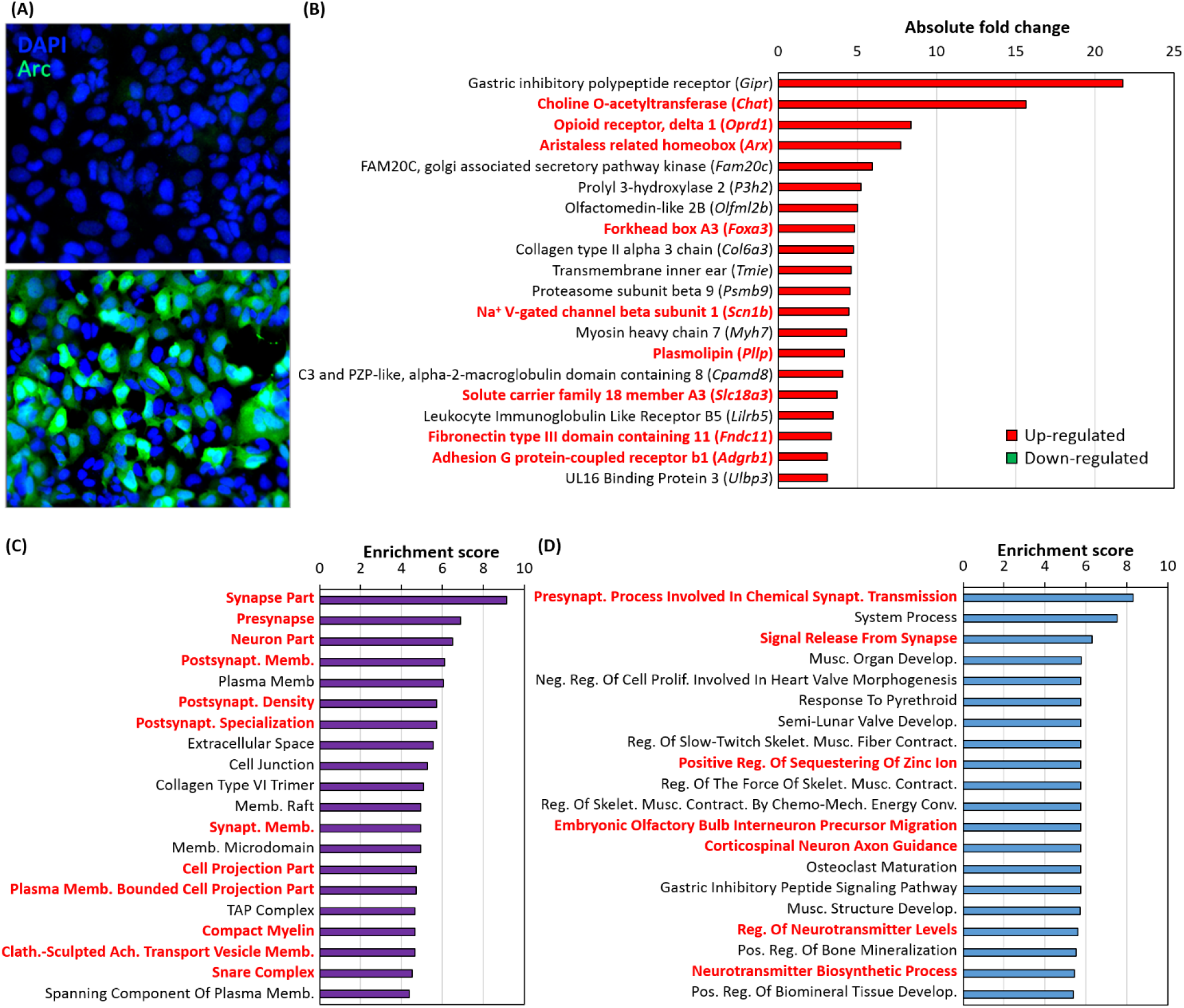
HEK293T cells exhibit neuronal properties upon induced expression of Arc. **(A)** Endogenous Arc expression was enhanced in HEK293T cells by targeting two single guide RNAs (sgRNAs) containing MS2 aptamers to the Arc promoter with the CRISPR/Cas9 Synergistic Activation Mediator system (see Methods for details). As a negative control, we used two sgRNAs targeting the promoter of the lac operon. Control cells (top) and Arc-induced cells (bottom) stained for Arc (green) and DNA was labeled with DAPI (blue). About 90% of the cells expressed Arc. **(B)** Graph showing the top 20 differentially expressed genes upon the induction of endogenous Arc in HEK293T cells. RNA-Seq was used to compare the mRNA levels between the Arc-induced and control HEK293T cells. Neuronal genes are bolded and highlighted in red. (**C**) and (**D**) GO analysis of the differential expressed genes upon overexpression of Arc. The top 20 cellular components (**C**) and biological processes (**D**) are presented. Neuronal features were bolded and highlighted in red. Na^+^: sodium; V-gated: voltage-gated; Postsynapt: postsynaptic; Memb: membrane; Synapt: synaptic; Clath: clathrin; Ach: acetylcholine; Presynapt: presynaptic; Musc: muscle; Develop: development; Neg: negative; Reg: regulation; Prolif: proliferation; Skelet: skeletal; Contract: contraction and Mech: mechanical; Conv: conversion.

Together with the results obtained with Arc knockdown in neurons, this finding strongly implicates Arc as a transcriptional regulator of neuronal development, synaptic function, plasticity and intrinsic excitability.

## DISCUSSION

Activity-regulated cytoskeleton-associated protein (Arc) was discovered in 1995 as a neuronal activity-dependent immediate-early gene^1,2^, which is rapidly transcribed in response to network activation associated with novel experiences^7–11^. Knockdown of Arc expression interferes with stabilization of short-term memory, indicating that Arc plays a critical role in memory consolidation^3,4^. Arc’s function has been most widely studied in excitatory synapses, where it regulates endocytosis of AMPA receptors^14,17^. Interestingly, AMPA receptor removal also underlies Aβ-induced synaptic depression and dendritic spine loss^108^, processes thought to be associated with cognitive dysfunction in Alzheimer’s disease (AD)^109^. In Arc knockout mice, long-term potentiation (LTP) is not stable, and dissipates within a few hours, consistent with the impaired memory consolidation observed in these mice^3–6^. However, the absence of the late form of LTP in Arc knockout mice cannot be explained by an AMPA receptor endocytosis deficit^4^, indicating that Arc must have additional functions. The data presented here identify a second function for Arc: regulation of neuronal activity-dependent transcription for genes associated with synaptic plasticity, intrinsic excitability and cellular signaling. Analysis of the differentially expressed genes points to Arc’s involvement of several neurological disorders, including Autism, Huntington’s and Alzheimer’s disease. This newly proposed role for Arc is supported by its interaction with chromatin and histone markers reported here (**Fig. 2-5, <u>Movie</u>**).

### Arc and chromatin

Pharmacological network stimulation induces Arc in a subset of cultured neurons (**Fig. 1**). Whereas chromatin in cultured hippocampal neurons is relatively uniform, Arc positive neurons are characterized by a larger number of densely packed heterochromatin puncta (chromocenters), likely harbouring silent genes, interspersed with highly open euchromatin domains, which are capable of active transcription (**Fig. 2**). This result is consistent with what has been observed *in vivo*, where Arc-deficient mice were found to have decreased heterochromatin domains^31^. These significant changes in chromatin structure observed in Arc-positive neurons are likely associated with equally substantial alterations in gene expression profiles. The correlation between Arc expression and chromatin remodeling that we observed does not establish a causative relationship. It is possible that Arc expression requires an alteration in chromatin structure, or alternatively, Arc expression may cause chromatin remodeling. Additional experiments are needed to decide on the underlying mechanism. It is also not clear at this time what determines which neurons will express Arc following network activation, although it likely has to do with the degree of participation of individual neurons in the enhanced network activity, which in turn depends on their synaptic connectivity.

Arc appears to physically interact with DNA: timelapse movies show dynamic chromatin loops that appear to invade Arc puncta (**Fig. 3**, **<u>Movie</u>**). The interaction is transient, lasting only a few seconds. Because these Arc puncta likely contain the histone acetylase Tip60^25^, it is conceivable that this interaction alters chromatin accessibility, thereby facilitating transcription. This idea is further strengthened by the association of Arc puncta with a histone marker for active enhancers (**Fig. 4**), as well as the close apposition between Arc puncta and puncta for a dual histone marker (H3K9Ac-S10P) that labels sites of active transcription (**Fig. 5**). A similar result has been obtained *in vivo*, where cocaine administration in rats results in an increase in nuclear Arc, which then associates with H3S10P^31^. Taken together, the data presented here on the interaction of Arc and chromatin may provide a mechanism for epigenetic regulation of gene transcription as the basis for memory consolidation.

### How does Arc regulate transcription?

Preventing Arc induction during neuronal network activation affects the transcription of a very large number of genes (**Fig. 6**). The domain structure of Arc protein appears to rule out that it can function as a transcription factor^78^. This raises the question: how does Arc regulate transcription? One possible mechanism, discussed above, is that Arc epigenetically controls gene transcription by regulating chromatin structure (through Tip60 or other chromatin remodelers) and modification of histones (e.g. H4K12Ac^25^). However, the differential gene expression associated with Arc knockdown is mediated through eleven upstream regulators identified by IPA (**Table 7**, **Fig. 11)**. This suggests that Arc has additional, less direct ways of regulating transcription. Interestingly, to date, none of the eleven upstream regulator proteins have been shown to either directly interact with or be modulated by Arc. They are also not transcriptionally controlled by Arc (except for *Sox2*) (**Table 7**). How then does Arc regulate transcription by activating or inhibiting these upstream regulators? Using IPA and its Ingenuity Knowledge Base, we were able to identify several known *interactors* of Arc that can modulate the action of the upstream regulators, which could then subsequentially alter gene transcription (**Fig. 13A**). Next, we will discuss the mechanisms by which four identified Arc interactors, NOTCH1, TIP60/*Kat5*, APP and GSK3B, could modulate the upstream regulators.

**Figure 13.**
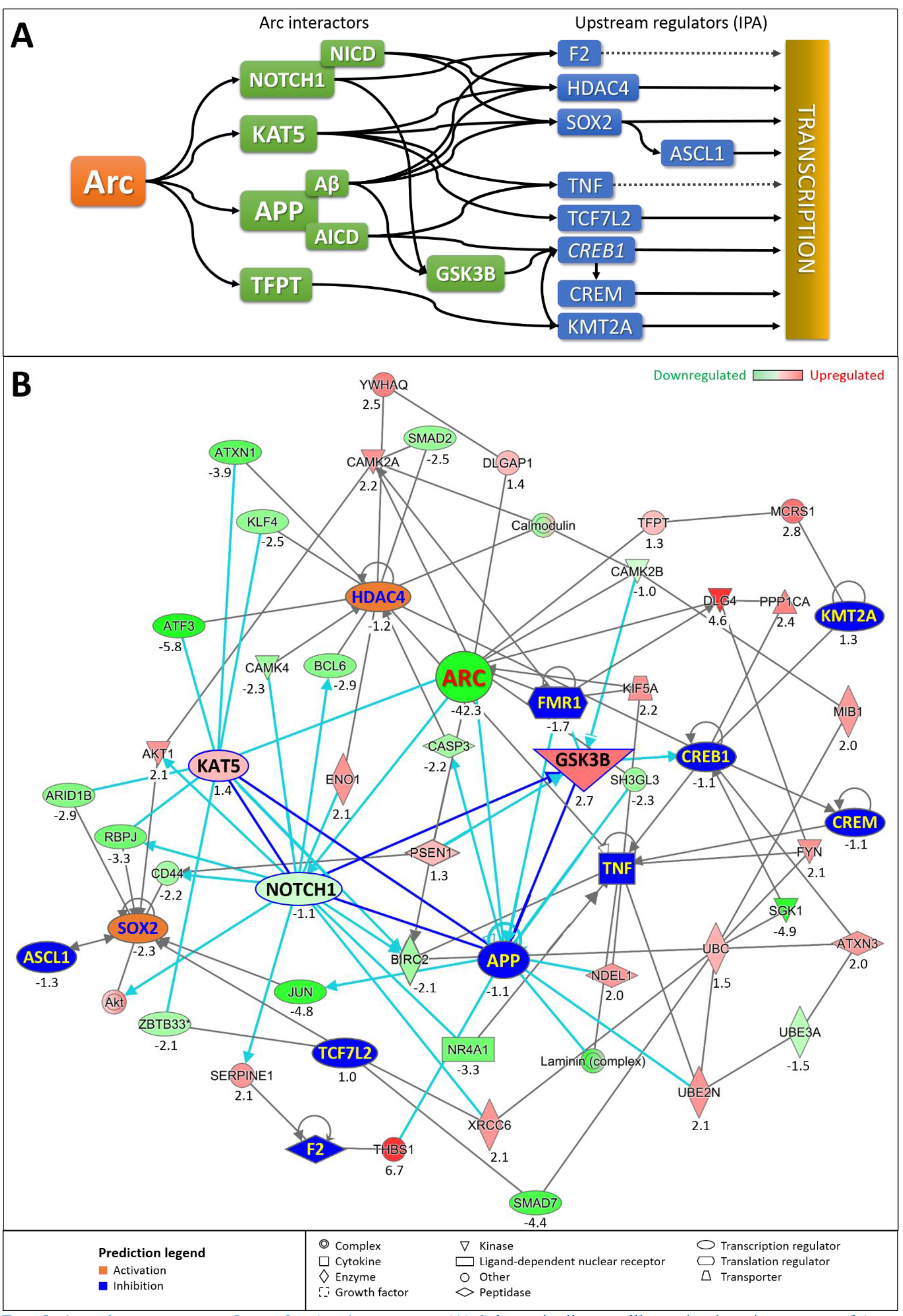
Regulation of upstream regulators by Arc interactors. **(A)** Schematic diagram illustrating how interactors of Arc could bring about changes in activity of upstream regulators identified by IPA, which in turn results in alteration of gene transcription. The dotted line indicates an indirect effect on transcription through the regulation of a transduction cascade. **(B)** Diagram showing the functional connectivity between Arc interactors (KAT5, NOTCH1, GSK3B and APP), upstream regulators highlighted in orange (activated) or blue (inhibited) and genes that mediate their interaction. Connections of Arc interactors with other genes are highlighted are highlighted in cyan. Connections between KAT5, NOTCH1, GSK3B and APP are highlighted in blue.

**Table 7.**
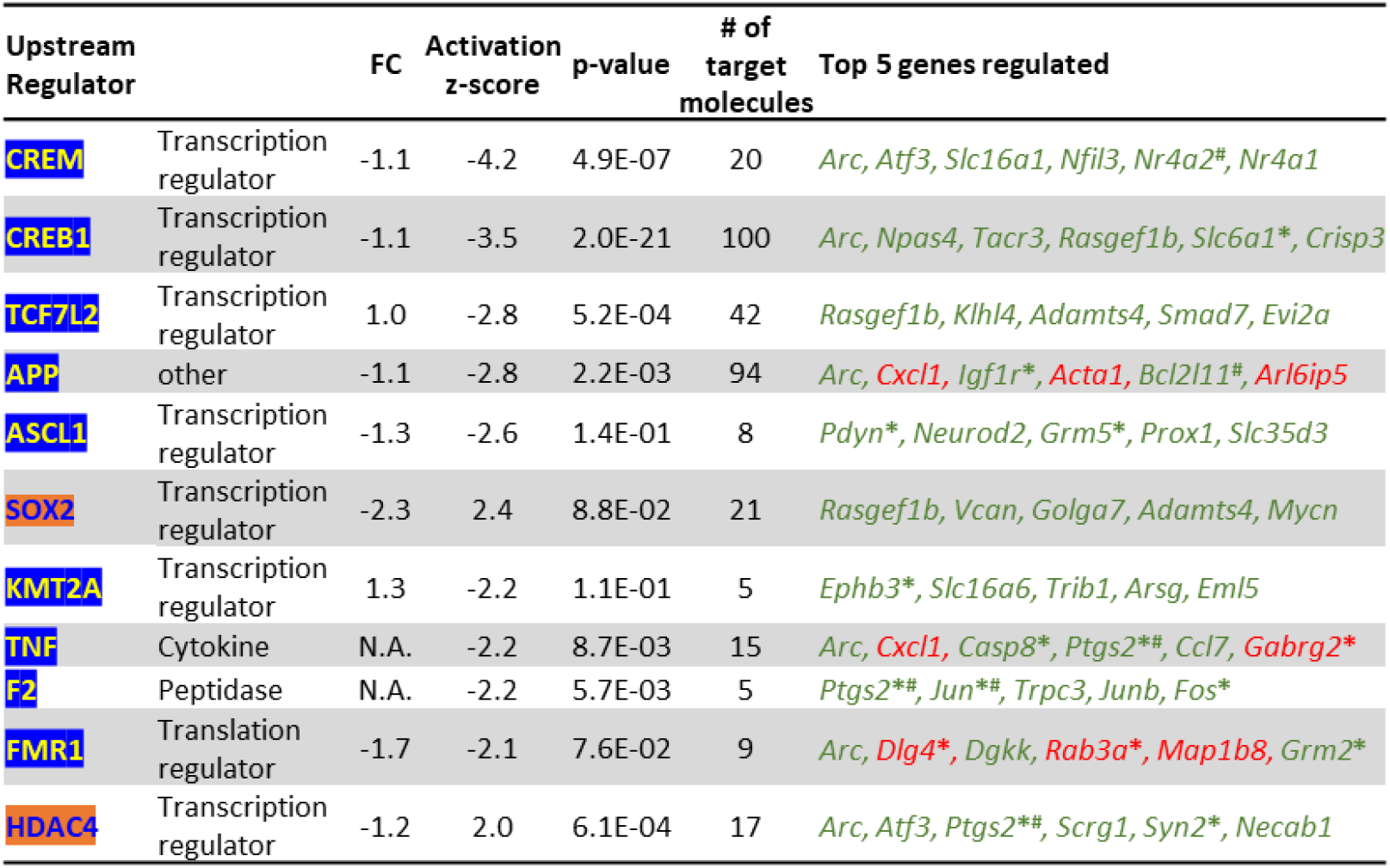
Upstream regulators associated with differential gene expression observed upon knockdown of Arc. Activation z-score indicates the predicted activity of upstream regulators by IPA analysis. Upstream regulators that were predicted to be inhibited are highlighted in blue while those activated are highlighted in orange. Regulated genes were highlighted in green (down-regulated) and red (up-regulated).

#### NOTCH1

NOTCH1 is a transmembrane receptor capable of signaling to the nucleus. Arc is required for the proteolytic cleavage of NOTCH1 to release its intracellular domain (NICD), which can translocate to the nucleus and alter transcription^110^. NICD regulates the expression of the transcriptional repressor BCL6^111^ and the activity of the calcium-dependent kinase CAMK4^112^, which in turn alter the localization and the nuclear-cytoplasmic shuttling of the histone deacetylase HDAC4, thereby affecting its downstream interactions/modulation^113,114^ (**Fig. 13B**). NOTCH1 could regulate the stability, nuclear localization and signaling of the transcription factor SOX2 through regulation of the protein kinase AKT1 and cell-surface glycoprotein CD44^115–118^ (**Fig. 13B**).

NOTCH1, through NICD, controls the expression of plasminogen activator inhibitor-1 (SERPINE1)^119^, an inhibitor of thrombin (F2)^120^ (**Fig. 13B**). NOTCH1 regulates the transcriptional activity of T-cell factor 4 (TCF7L2)^121^, through its interaction with the DNA-repair protein Ku70 (XRCC6)^122^ (**Fig. 13B**). NOTCH1 interacts with the nerve growth factor NR4A1/Nur77^123^, thereby modulating expression levels of the cytokine tumor necrosis factor alpha (TNF)^124^. Finally, NOTCH1 regulates the expression level of the inhibitor of apoptosis protein cIAP1/*Birc2*^125^, which also affects TNF expression^126^ (**Fig. 13B**).

#### TIP60/*Kat5*

The *Kat5* gene encodes TIP60, a member of the MYST family of histone acetyl transferases, which plays important roles in chromatin remodeling and transcription regulation^127^. In the fruit fly *Drosophila*, TIP60 has been implicated in epigenetic control of learning and memory^128^, while it mediates APP-induced apoptosis and lethality in a fly AD model^129^. Nuclear Arc interacts with TIP60 at perichromatin regions and recruits TIP60 to PML bodies, sites of epigenetic transcription regulation^25^. Arc levels correlate with acetylation status of H4K12, a substrate of TIP60 and a memory mark that declines with aging^29^, suggesting that Arc mediates activation of TIP60. TIP60/*Kat5* facilitates the repressive action of HDAC4 through the formation of complexes with the zinc-finger transcription factor KLF4^121,130^, the cAMP-dependent transcription factor ATF3^131–133^ and the neurodegenerative disease protein ataxin-1 (ATXN1)^134,135^ (**Fig. 13B**). Arc’s interaction with TIP60/*Kat5* may result in a complex being formed at the cIAP1/*Birc2* promoter region^136^ to mediate downstream signaling of TNF^126^ (**Fig. 13B**). TIP60/*Kat5* forms a complex with the Kaiso transcription factor ZBTB33^137^ resulting in the inhibition of the TCF7L2 transcriptional complex^138^ (**Fig. 13B**). Complexing of TIP60 with ARID1B could affect SOX2 signaling^139–141^. The regulation of SOX2 by TIP60/Kat5 could also have an implication on the transcriptional activity of Achaete-Scute homolog 1 (ASCL1), as SOX2 and ASCL1 regulate each other, possibly as a feedback loop^142,143^.

#### APP

The functional interaction between APP and Arc is crucial for Arc’s modulation of upstream regulators (**Fig. 13A**). Arc interacts with endophilin 2/3 (SH3GL3) and dynamin on early/recycling endosomes to alter the trafficking and localization of APP. The association of Arc with presenilin 1 (PSEN1) promotes the trafficking of γ-secretase to endosomes and enzymatic cleavage of APP^80^ (**Fig. 13B**). The generation of amyloid beta through APP cleavage leads to an altered downstream signaling, activity and production of HDAC4, SOX2 and F2 through changes of caspase-3 (CASP3)^144,145^, JUN^146,147^ and thrombospondin-1 (THBS1)^148,149^, respectively (**Fig. 13B**). Cleavage of APP generates a cytosolic fragment, AICD, which forms a transcriptionally active complex with TIP60 and the transcription factor FE65^150^. AICD also modulates the ubiquitin-proteasome system (UPS) via UBE2N^151^, to change downstream signaling induced by TNF^152^ (**Fig. 13B**). The modulation of the UPS via UBE2N, UBC and UBE3A^153^ could implicate the ubiquitination of serum- and glucocorticoid-regulated kinase-1 (SGK-1)^154–156^ and polyglutamine-expanded ataxin 3 (ATXN3)^157^ and their ability to regulate the transcription factor cAMP responsive element binding protein 1 (CREB1)^158,159^ (**Fig. 13B**). The modulation of CREB1 would further implicate changes in expression levels of cAMP responsive element modulator CREM^160–162^ (**Fig. 13B**). Finally, APP has a role in the regulation of TNF through indirect modulation of CREM^163^ and direct interactions with laminin could regulate the production of TNF^164,165^ (**Fig. 13B**).

#### GSK3B

Although glycogen synthase kinase 3 beta (GSK3B) is not regulated by Arc, promotion of cleavage of APP to amyloid beta enhances the induction and activation of GSK3B^166–168^. This could lead to modified downstream signaling of CREB1^169^ (**Fig. 13B**). GSK3B is also a downstream mediator of NOTCH1^170^, PSEN1^171^, and CAMK2B^172^, all of which are Arc interactors^80,110,173^. This creates an interesting situation as APP/amyloid beta is positively regulated by GSK3B^174,175^, creating a positive feedback loop for amyloid beta production and its downstream signaling^166–168^ (**Fig. 13B**).

#### Interactions among TIP60, NOTCH1 and APP

A delicate regulatory network exists among Arc’s interactors TIP60/*Kat5*, NOTCH1 and APP (**Fig. 13B**). Arc’s activation of the γ-secretase PSEN1 to promote cleavage of APP not only increases amyloid beta load, but also results in an increased level of the APP intracellular domain (AICD)^80,176^. AICD forms a complex with TIP60/*Kat5* to alter transcriptional activity crucial for AD progression^150,177–182^ (**Fig. 13B**). This AICD-TIP60 interaction is disrupted by NICD, formed when Arc activates NOTCH1^110^, thereby downregulating AICD signaling while promoting NICD signaling^183,184^ (**Fig. 13B**). The formation of NICD and AICD is competitive, as NOTCH1 and APP are both substrates of γ-secretase^185^, whose activity is regulated by Arc^80^. In addition, the induction of TIP60 histone acetylation activity by Arc^25^ could also increase the negative regulation of NOTCH1^183^ (**Fig. 13B**). This highlights Arc as an important modulator of the relationship and downstream signaling mediated by NOTCH1, TIP60/*Kat5* and APP. Of note, the mRNA levels of *Notch1, Kat5* and *App* were not significantly altered upon knockdown of Arc, indicating that the transcriptional changes brought about were due to protein interaction and activation (**Fig. 13B**), which is upstream of transcription (**Fig. 13A**). However, the modulation of upstream regulators by Arc is also dependent on the its subcellular localization.

#### Arc’s subcellular localization determines its function

When Arc is localized outside the nucleus it tends to accumulate in dendrites and spines, small membrane protrusions that harbour excitatory synapses. Here, Arc controls the removal of AMPA receptors by endocytosis, allowing it to regulate synaptic efficacy^14,17^. Synaptic Arc also associates with the synaptic scaffolding protein PSD-95/*Dlg4*, which complexes with the tyrosine kinase FYN^186–188^, allowing it to regulate brain-derived neurotrophic factor (BDNF) signalling through tyrosine receptor kinase B (TrkB), a major pathway for synapse maturation, plasticity and neurodevelopmental disorders^189^. Activation of FYN could also mediate the secretion of TNF^190^(**Fig. 13B**). A high-affinity interaction with calcium-calmodulin kinase 2 beta (CAMK2B) targets Arc to inactive synapses, where it removes GluA1 AMPA receptors from the postsynaptic membrane surface^173^.

Arc has been shown to possess both a nuclear localization signal (NLS) and a nuclear retention domain^24^, allowing it to translocate to the nucleus autonomously. Once in the nucleus, Arc has access to several other potential binding partners, including a nuclear spectrin isoform (βSpectrinIV∑5)^20^ and TIP60, a subunit of a chromosome remodeling complex^25^. Association with Amida, encoded by the *Tfpt* gene (**Fig. 13B**) facilitates Arc’s entry into the nucleus^191^. Amida is a subunit of the INO80 chromatin remodeling complex, which contains the transcriptional regulator MCRS1^192,193^. MCRS2, an isoform of MCRS1, is associated with the MLL chromatin remodeling complex, which also contains KMT2A (MLL1) (**Fig. 13B**). Arc’s association with Amida and possibly the INO80 and MLL complexes may provide Arc with yet another opportunity to control gene expression by altering chromatin structure.

The ability of Arc to translocate between the synapse and the nucleus, with unique functions in each subcellular compartment, further strengthens its role in memory consolidation, which requires both alterations of synaptic function and *de novo* gene transcription^194^.

### Arc controls synaptic plasticity and intrinsic excitability

Arc’s well-studied ability to alter synaptic efficacy by endocytosis of AMPA receptors established it as a critical regulator of synaptic plasticity^14,17,188,195^. Whereas this mechanism of activity-dependent removal of glutamate receptors supports Arc’s role in mediating long-term depression (LTD)^196–199^, it does not explain the absence of stable long-term potentiation (LTP) observed in Arc knock-out mice^4^. Because late-LTP is considered a critical cellular mechanism underlying memory consolidation, the molecular and cellular mechanism by which Arc supports memory stabilization has remained elusive. The data presented here showing that Arc transcriptionally regulates the expression of a large number of synaptic proteins, with functions in both the pre- and post-synaptic compartment (**Table 2**), provides a new mechanism by which Arc can control long-lasting changes in synaptic structure and function required for memory consolidation.

Formation of a memory trace not only requires longterm changes in the strength of the synapses connecting the neurons that constitute the engram, but also stable changes in their intrinsic excitability^200–202^. Because Arc controls the expression of a large number of ion channels and pumps/transporters, it appears that Arc is capable of supporting this functional aspect of memory consolidation as well.

### Arc and Alzheimer’s disease

Alzheimer’s Disease (AD) is a devastating neurodegenerative disorder^203,204^ characterized by the progressive loss of both synaptic function^205^ and long-term memory formation^206^. There is currently no therapy that prevents, stabilizes or reverses the progression of this disease, which is projected to take on epidemic proportions as the world population ages^207,208^. Several previous studies have revealed an association between *Arc* and AD. A landmark study published in 2011 showed that *Arc* protein is required for the formation of amyloid (Aβ) plaques^80^. Moreover, *Arc* protein levels are aberrantly regulated in the hippocampus of AD patients^209^, and are locally upregulated around amyloid plaques^210^, whereas a polymorphism in the *Arc* gene confers a decreased likelihood of developing AD^211^. It has been shown that spatial memory impairment is associated with dysfunctional Arc expression in the hippocampus of an AD mouse model^212^. These published results together with the data presented here, suggests that aberrant expression or dysfunction of Arc contributes to the pathophysiology of AD^205,213^.

### Arc and AD therapy

Arc’s ability to transcriptionally regulate AD susceptibility and AD pathophysiology related genes indicates a possibility for modifying expression and activity of Arc as a therapy for AD. Current treatments are symptomatic, not effective disease-modifying cures^214^. Many hypotheses have been proposed to underlie the development of AD, including i) amyloid beta aggregation, ii) tau hyperphosphorylation, iii) neuroinflammation, iv) neurotransmitter dysfunction, v) mitochondria dysfunction, vi) glucose metabolism, vii) vascular dysfunction and viii) viral infection^214–217^. These hypotheses have generated many new compounds, none of which showed efficacy in slowing cognitive decline or improving global functioning^214,216^. Arc appears a good therapeutic candidate for AD, because of its involvement in amyloid beta production, tau phosphorylation, neuroinflammation and neurotransmission. Moreover, we have shown that Arc can modulate the expression of many AD genetic risk factors and genes associated with the pathophysiology of AD (**Fig. 9**, **Fig. 10**, **Fig. 12** and **Table 4**). Currently, known drugs that could increase mRNA or protein expression of Arc include antidepressant drugs^218^, phencyclidine^219^ and corticosterone, a memory enhancing drug^220^. Arc expression could be altered by targeting TIP60 and PHF8, two histone modifiers that together control Arc transcription^22^. Drugs could also modulate Arc’s effect via its interactors such as TIP60 and NOTCH1. Natural and synthetic drug molecules targeting TIP60 exist, but they are currently used for cancer treatment^221^. Modulation of NOTCH1 function often involves inhibitors of γ-secretase, which would also affect APP cleavage^185,222^. These pharmaceutical modifications of Arc expression and activity could present a promising starting point for development of a more effective AD therapy.

## METHODS

### Animals and chemicals

All work involving the use of animals were performed according to the guidelines of the Institutional Animal Care and Use Committee (IACUC) and were approved by the IACUC at the SingHealth in Singapore. Time-mated E18 Sprague Dawley rats were purchased through the SingHealth Experimental Medicine Centre (SEMC). They were sacrificed immediately after delivery to the vivarium. All chemicals were purchased from Sigma-Aldrich unless otherwise stated.

### Culturing hippocampal and cortical neurons

Hippocampal and cortices were dissected from the E18 embryos of Sprague Dawley rats. Hippocampi or cortices underwent papain dissociation based on the protocol from the Papain Dissociation System (Worthington Biochemical Corporation). Gentle mechanical trituration was performed to ensure complete dissociation of tissues. Dissociated cells were plated on poly-D-lysine (Sigma) coated dishes at a plating density of 1.5 x 10^5^/cm^2^ in Neurobasal medium (Gibco) supplemented with 10% (v/v) foetal-bovine serum (FBS, Sigma), 1% (v/v), penicillin-streptomycin (P/S, Gibco) and 2% (v/v) B27 supplement (Gibco) for 2 hours. FBS-containing medium was then removed and replaced with FBS-free medium and cells were cultured FBS-free subsequently to prevent astrocytic over-growth. Medium was changed on Days In Vitro (DIV) 5. Subsequently, medium was changed every three to four days. Experiments were carried out on DIV 18-22.

### Inhibition of Arc expression by an shRNA

Four Arc shRNA plasmids (SureSilencing, Qiagen) were transfected into neuronal cultures using Lipofectamine 2000 (Qiagen). Pharmacological LTP was induced in neuronal cell cultures using a 4-hr treatment with **4BF**. Cells were fixed and stained for Arc protein. Immunofluorescence images were obtained using wide-field microscopy. Effectiveness of inhibition of Arc expression was based on co-occurrence of expression of the plasmids and the absence of Arc immunofluorescence. The most effective shRNA plasmid was chosen and adeno-associated virus AAV9 constructs harbouring an Arc shRNA and a scrambled version of this shRNA were synthesized using the annealed oligo cloning method. The oligos for the Arc shRNA were: i) 5′-GAT CCG GAG GAG ATC ATT CAG T-3′, ii) 5′-ATG TCT TCC TGT CAA CAT ACT GAA TGA TCT CCT CCT TTT TG-3′, iii) `5′-AAT TCA AAA AGG AGG AGA TCA TTC AGT-3′ and` iv) 5′-ATG TTG ACA GGA AGA CAT ACT GAA TGA TCT CCT ccG-3′. The oligos for Arc scrambled shRNA were i) 5′-GAT CCG GTA ATT TCG GAG GAT C-3′, ii) 5′-AAG TCT TCC TGT CAA CTT GAT CCT CCG AAA TTA CCT TTT TG-3′, iii) `5′-AAT TCA AAA AGG TAA TTT CGG AGG ATC-3′ and` iv) 5′-AAG TTG ACA GGA AGA CTT GAT CCT CCG AAA TTA CCG-3′. The ends of the annealed oligos harbor overhangs of the restriction sites for BamH1 and EcoR1. Oligos for the Arc shRNA were annealed in buffer A (mM) 100 NaCl and 50 HEPES, pH 7.4 while oligos for Arc scrambled shRNA were annealed in buffer B (mM) 10 Tris, pH 7.5-8.0, 50 NaCl and 1 EDTA at equimolar concentration by heating to a temperature of 95°C for 5 min then cooling it down to room temperature (rtp). The annealed oligos were ligated using T4 ligase (New England Biolabs) into the BamH1/EcoR1-cut vector pENN.AAV.U6.shRLuc.CMV.eGFP.SV40, generously provided by the University of Pennsylvania, Vector Core. Ligated products were transformed into Stbl3 competent cells (Thermo Fisher Scientific). Successful constructs were identified by restriction enzymes digestion and verified by sequencing. AAV9 virus harbouring the transgenes (concentrations at 1 x 10^13^– 1 x 10^14^ GC/ml range) were synthesized by University of Pennsylvania, Vector Core. Arc expression was prevented by treating neuronal cultures with 3 x 10^6^ multiplicity of infection (MOI) AAV9 Arc shRNA virus on DIV14. Induction of Arc expression by pharmacological LTP (see below) was performed between DIV19-22.

### Pharmacological LTP and immunofluorescence

Hippocampal or cortical neuronal cultures were treated with a combination of 100 μM 4-aminopyridine (4AP), 50 μM bicuculline (Bic) and 50 μM forskolin for the respective time stated to induce pharmacological LTP^21,32,33^. This drug combination will be referred to as **4BF** henceforth. At the end of the treatment, cells were fixed with 100 % ice-cold methanol at −20°C for 10 min. Cells were washed three times with 1x Phosphate Buffered Saline (PBS, in mM: 137 NaCl, 2.7 KCl and 12 phosphate buffer) containing 0.1% (v/v) Triton X-100 (PBS-Tx). Depending on the antibodies used, some cells were fixed again with 4% (w/v) paraformaldehyde (PFA) in 1x PBS containing 4% (w/v) sucrose. Cells were washed three times in 1x PBS-Tx and blocked in 2% (w/v) Bovine Serum Albumin (BSA) in 1x PBS for 1 h at rtp. Depending on the species the secondary antibodies were raised in, 10% (v/v) serum of the corresponding species was added to the blocking buffer. Cells were probed with primary antibodies as indicated for the experiments: i) anti-Arc (1:300, Santa Cruz, sc-17839), ii) anti-Arc (1:300, Synaptic Systems, 156 003), iii) anti-MAP2 (1:300, Millipore, AB5622), iv) anti-H3K27Ac (1:300, Wako, 306-34849) and v) anti-H3K9Ac-S10P (1:300, Abcam, ab12181) in antibody dilution buffer (1x PBS containing 1% (w/v) BSA, 5% (v/v) serum and 0.05% (v/v) Triton X-100) for 1 h at rtp. Cells were washed three times in 1x PBS-Tx. Cells were then probed with 1:1000 Alexa-Fluor 647, Alexa Fluor 568 or Alex-Fluor 488 (Molecular Probes) for 1 h at rtp. Cells were washed three times, followed by staining of DNA with 50 μM DAPI for 20 min at rtp. Cells were mounted in FluorSave (Calbiochem). For immunofluorescence staining for STORM imaging, cells were fixed with 3% paraformaldehyde and quenched with 0.1% NaBH4 as described in Oey *et al*. (2015)^22^. Blocking, primary and secondary antibody staining were carried out as above. A post-fixation, as described in Oey *et al*. (2015)^22^, was also carried out after secondary antibody binding.

### Transfection of neuronal cultures

The Arc-eYFP construct was generated as described in Bloomer *et al*. (2007)^20^. Neuronal cultures (DIV16) were transfected with Arc-eYFP and H2B-mCherry (Addgene, 20972) with Lipofectamine 2000 (Invitrogen) according to manufacturer’s protocol with some adjustment. Arc-eYFP:H2B-mCherry DNA was added to Lipofectamine at a ratio of 1:1. The Lipofectamine:DNA complex was incubated at rtp for 20 min before been added to the cells. The complex was added dropwise such that it was evenly distributed on the cell culture. Culture medium was added after 20 min and experiments were performed on DIV19.

### Widefield microscopy

Fluorescence images were obtained using widefield microscopy as detailed in Oey *et al*. (2015)^22^. Images obtained were analyzed using NIS Elements AR version 4.1 (Nikon) to perform background subtraction. Out-of-focus fluorescence was removed using 3D deconvolution (AutoQuant, Media Cybernetics). The Region-Of-Interest (ROI) analysis tool was used to mark nuclei based on DAPI intensity. Corresponding mean Arc intensity of each nucleus was also measured using the automated measurement module. The average of mean Arc intensity for all neurons from non-**4BF** stimulated controls were obtained for each set of experiments. This would be used as a cut-off threshold between Arc-positive and Arc-negative neurons for each set of experiments since Arc expression was only observed upon stimulation^223,224^. Nuclei images were cropped individually and analyzed using a custom MATLAB (Mathworks) program. Size and intensity thresholds were applied to identify and quantitate puncta in each nucleus. Batch processing using the same size and intensity threshold was performed. Mean size of the puncta and number of puncta were recorded. ROI ID for each nucleus was used to correlate the mean Arc intensity with the mean area or number of puncta. Statistical analysis was performed using GraphPad Prism Version 6.01. Statistical data shown are mean ± S.E.M. (standard error of the mean) across experiments.

### Spinning Disc confocal microscopy

Fluorescence images and time-lapse movies were obtained using a motorized Ti-E inverted microscope (Nikon) with a 60X oil Plan-Apo objective (1.49 NA) and a 100X Apo-TIRF objective (1.49 NA). Spinning disk confocal microscopy was achieved using the CSU-W1 Nipkow spinning disk confocal unit (Yokogawa Electric). A sCMOS camera (Zyla, Andor) was used to capture the confocal images. Laser lines used were 488 nm (100 mW) for GFP, 515 nm (100 mW) for eYFP and 561 nm (150 mW) for mCherry (Cube lasers, Coherent). Fast excitation/emission switching was obtained using a dichroic beam splitter (Di01-T405/488/568/647-13 x 15 x 0.5, Semrock) and filter wheels controlled by a MAC6000DC (Ludl). The Perfect Focus System (Nikon) was applied to ensure minimal focus drift during image acquisition. Z stacks were obtained using step sizes recommended for objectives used, which were processed using 3D blind deconvolution (AutoQuant) to remove out-of-focus fluorescence.

### Stochastic Optical Reconstruction Microscopy (STORM)

Dual color STORM image sequences were obtained using a Zeiss ELYRA PS.1 platform. Endogenous Arc and the dual histone marker H3K9Ac-S10P were labeled with primary antibodies and visualized using Alexa 488 and Alexa 647 secondary antibodies. Time-lapse movies of 10,000 frames were obtained of neuronal nuclei expressing Arc capturing the blinking of individual Alexa 488 and 647 molecules brought into the dark state by intense laser illumination. Fitting of a 2D Gaussian to each blinking dot allowed their XY localization to be determined with high precision (typically 30 nm). Super resolution images are generated from the localizations by superimposing a 2D Gaussian (green for 488 nm, red for 647 nm) for each localized position. Molecule localization and image rendering were performed by Zen software (Zeiss).

### Cell lysate preparation and Western blotting

Following **4BF** stimulation, neuronal cultures were washed gently with 1x PBS. Cells were gently scraped off and harvested in an Eppendorf tube. Cells were spun down at 10000 g for 5 min at 4°C to obtain the cell pellet. Total protein was isolated using an RNA-protein extraction kit (Macherey-Nagel), as specified by the manufacturer. A BCA kit (Pierce) was used to measure the concentration of proteins. 30 μg of each protein sample was denatured and reduced by boiling at 95°C for 5 min in 10% (v/v) 2-mercaptoethanol-containing Laemmli sample buffer (Bio-Rad). Samples were resolved by SDS-PAGE with a precast Tris-glycine gel (Bio-Rad) and transferred onto PVDF membranes using the Trans-Blot Turbo transfer System (BioRad) as indicated by manufacturer. Membranes were blocked for 1 h at rtp with 5% (w/v) non-fat milk block (Bio-Rad) in 1x TBS (in mM) (140 NaCl, 3 KCl, 25 Tris base) (First Base) containing 0.1% (v/v) Tween-20 (TBST), followed by primary antibodies incubation for 1 h (anti-Arc, 1:1000, Santa Cruz, sc-17839) in 1x TBST at rtp. Membranes were washed three times, each for 5 min in 1x TBST at rtp. Secondary antibodies binding was performed using the corresponding HRP-conjugated secondary (1:10000, Invitrogen) for 1 h in 1x TBST at rtp. Protein bands were detected with chemiluminescence substrate (Pierce) visualized with a Gel Doc XRS imaging system (Bio-RAD) or developed on scientific imaging film (Kodak).

### RNA sample preparation, library construction, RNA-Seq and analysis

**4BF** treated neuronal cells were washed, scraped and spun down as above. RNA samples were obtained from the cell pellet using the RNA-protein extraction kit as specified by manufacturer (Macherey-Nagel). Library construction and RNA sequencing were performed by the Duke-NUS Genome Biology Facility. 2.2 μg of RNA was used for library construction. Prior to library construction, quality of the RNA was analyzed with an Agilent Bioanalyzer. Following poly-A enrichment, recovered RNA was processed using the Illumina TruSeq stranded mRNA kit to generate the adaptor-ligated libraries. A total of 9 samples were analyzed. These samples came from 3 different sets of experiments (n = 3). Each set contained samples treated with i) 8 h **4BF,** ii) Arc shRNA + 8 h **4BF** and iii) Arc scrambled shRNA + 8 h **4BF**. Six samples were sequenced per lane on the HiSeq 3000 sequencer using 150 pair-end reads. For the HEK293T cells, RNA was obtained similarly. Three samples were analyzed, with two Arc induced samples and one control sample. The samples were processed as described above and sequenced on 1 lane on the HiSeq 3000 sequencer.

### Computational analyses of RNA-Seq data

FASTQ files obtained from the RNA-sequencing were mapped to the rat genome using Partek Flow (version 7.0.18.1210). Adapter sequences were trimmed. Contaminant reads contributed from rDNA, tRNA and mtDNA were filtered out using Bowtie2 (version 2.2.5). Filtered, trimmed reads of high quality (Phred score > 30) were then mapped onto the *Rattus norvegicus* genome (rn6) for the rat samples or Homo sapiens genome (hg38) for the HEK293T samples with Star (version 2.5.3a)^225^. Post alignment QA/QC were performed after the alignment step to determine if alignment had a good average coverage and were uniquely aligned. The unique, paired reads were used for gene expression quantification. Reads were assigned to genes using the Expectation/ Maximization (E/M) algorithm in Partek Flow^226^ based on the annotation model rn6 (Ensembl transcripts release 93) for the rat samples and the annotation model hg38 (Ensembl transcripts release 94) for the HEK293 samples. To ensure only informative genes were included in the downstream analysis, noise (maximum feature counts ≤ 30) was filtered out. Read counts between samples were normalized with the Upper quantile method^227^. As genes with very low expression might be inadequately represented and incorrectly identified as differentially expressed, a constant of 1 was added to normalized counts for rectification. Statistical analysis was performed using the Gene Specific Analysis (GSA) module in Partek Flow to identify differential gene expression. P-value and fold changes of differentially expressed genes were calculated based on the lognormal with shrinkage distribution. Average coverage of less than 3 were also filtered out prior to statistical analysis. Differential gene expression with a cut-off value of false discovery rate (FDR) step-up < 0.05^228^ and an absolute fold change ≥ 2 were considered for further gene ontology analysis in Partek Flow that is based on the GO Consortium (version 2018_08_01)^229,230^. GO analysis was also performed on the transcriptional regulators/factors that were observed to be altered upon Arc knockdown using DAVID (version 6.8)^81,231^. The EASE score obtained from the DAVID analysis is a modified Fisher Exact P-value to indicate gene enrichment in the annotation terms. Functional analysis on the statistically significant differential gene expression (FDR step-up < 0.05; absolute fold change ≥ 2) was performed by Ingenuity Pathway Analysis (IPA) (https://tinyurl.com/y6672c22, Qiagen). Pathways and their associated downstream effects, diseases, regulator networks and upstream regulators were identified by IPA. Predictions on the possible activation and inhibition of pathways, downstream effects and upstream regulators were inferred from the degree of consistency in the expression of the target genes compared to the fold changes in the differentially expressed gene list. This activation or inhibition status was expressed as a z-score, with z ≥ 2 indicating activation and z ≤ 2 indicating inhibition. Inferences made were based on at least one publication or from canonical information stored in the Ingenuity Knowledge Base. Fisher’s exact test was used to calculate the P-value for all analyses in IPA.

### Plasmid construction for Arc expression in HEK293T cells

The following plasmids were used: pSBbi-Hyg (Addgene #60524) and pSBbi-Pur (Addgene #60523) were a gift from Eric Kowarz. pCMV(CAT)T7-SB100 (Addgene #34879) was a gift from Zsuzsanna Izsvak. sgRNA(MS2) cloning backbone plasmid, (Addgene #61424), MS2-P65-HSF1_GFP, (Addgene #61423), and dCAS9-VP64_GFP (Addgene #61422) were a gift from Feng Zhang. The psBbi-Hyg-dCAS9-VP64 and the pSBbi-MS2-P65-HSF1-Pur plasmids were constructed by isolating the dCAS9-VP64 and MS2-P65-HSF1 sequences via PCR from the MS2-P65-HSF1_GFP and dCAS9-VP64_GFP plasmids and annealed into the SfiI-linearised pSBbi-Hyg and pSBbi-Pur plasmids. psBbi-Hyg-dCAS9-VP64 and pSBbi-MS2-P65-HSF1-Pur were then co-transfected with pCMV(CAT)T7-SB100 into HEK293T using JetPrime (Polyplus-Transfection) according to manufacturer’s instructions. HEK293T cells with successful transposition of both genes were selected with a combination of 0.75ug/ml puromycin (Gibco) and 200ug/ml hygromycin B (Nacalai) in DMEM + 10% FBS (Gibco) over several passages for a month.

### Transfection for endogenous Arc overexpression and purification of mRNA

sgRNA(MS2) backbone plasmids containing guide RNAs complementary to human Arc promoters were transfected into the mutated HEK293T cells. A separate control well was transfected with sgRNA(MS2) backbone containing LacZ promoter sgRNAs. After 48 hours, mRNA was purified using the NucleoSpin RNA kit (Macherey-Nagel) and submitted for RNA-sequencing by the Duke-NUS Genomics Core Facility. The table below lists the guide RNAs (sgRNAs) used:

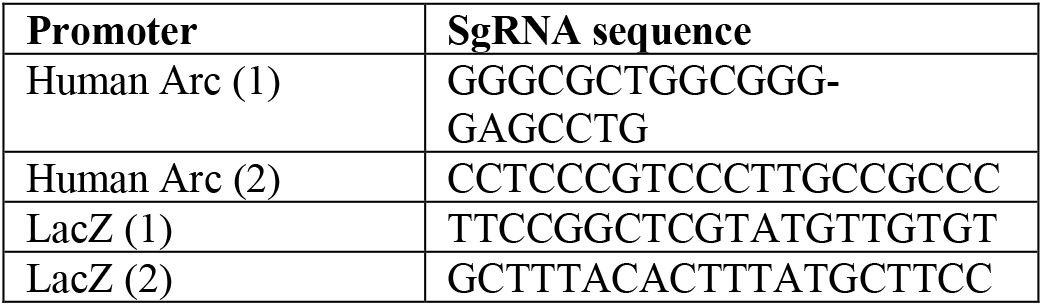

## ACKNOWLEDGEMENTS

We would like to thank the SingHealth Advanced BioImaging Core for excellent support with our microscopy experiments. The experiments were supported by grant OFIRG/0019/2016 from the National Medical Research Council (NMRC) to AMJ VanDongen.

## REFERENCES

1. Lyford, G.L., et al., Arc, a growth factor and activity-regulated gene, encodes a novel cytoskeleton-associated protein that is enriched in neuronal dendrites. Neuron, 1995. 14(2): 433–45.

2. Link, W., U. Konietzko, G. Kauselmann, M. Krug, B. Schwanke, U. Frey and D. Kuhl, Somatodendritic expression of an immediate early gene is regulated by synaptic activity. Proc Natl Acad Sci U S A, 1995. 92(12): 5734–8.

3. Guzowski, J.F., G.L. Lyford, G.D. Stevenson, F.P. Houston, J.L. McGaugh, P.F. Worley and C.A. Barnes, Inhibition of activity-dependent Arc protein expression in the rat hippocampus impairs the maintenance of long-term potentiation and the consolidation of long-term memory. J Neurosci, 2000. 20(11): 3993–4001.

4. Plath, N., et al., Arc/Arg3.1 is essential for the consolidation of synaptic plasticity and memories. Neuron, 2006. 52(3): 437–44.

5. Ploski, J.E., V.J. Pierre, J. Smucny, K. Park, M.S. Monsey, K.A. Overeem and G.E. Schafe, The activity-regulated cytoskeletal-associated protein (Arc/Arg3.1) is required for memory consolidation of pavlovian fear conditioning in the lateral amygdala. J Neurosci, 2008. 28(47): 12383–95.

6. Maddox, S.A. and G.E. Schafe, The activity-regulated cytoskeletal-associated protein (Arc/Arg3.1) is required for reconsolidation of a Pavlovian fear memory. J Neurosci, 2011. 31(19): 7073–82.

7. Guzowski, J.F., B.L. McNaughton, C.A. Barnes and P.F. Worley, Environment-specific expression of the immediate-early gene Arc in hippocampal neuronal ensembles. Nat Neurosci, 1999. 2(12): 1120–4.

8. Guzowski, J.F., B. Setlow, E.K. Wagner and J.L. McGaugh, Experience-dependent gene expression in the rat hippocampus after spatial learning: a comparison of the immediate-early genes Arc, c-fos, and zif268. J Neurosci, 2001. 21(14): 5089–98.

9. Chawla, M.K., J.F. Guzowski, V. Ramirez-Amaya, P. Lipa, K.L. Hoffman, L.K. Marriott, P.F. Worley, B.L. McNaughton and C.A. Barnes, Sparse, environmentally selective expression of Arc RNA in the upper blade of the rodent fascia dentata by brief spatial experience. Hippocampus, 2005. 15(5): 579–86.

10. Ramirez-Amaya, V., A. Vazdarjanova, D. Mikhael, S. Rosi, P.F. Worley and C.A. Barnes, Spatial exploration-induced Arc mRNA and protein expression: evidence for selective, network-specific reactivation. J Neurosci, 2005. 25(7): 1761–8.

11. Vazdarjanova, A., et al., Spatial exploration induces ARC, a plasticity-related immediate-early gene, only in calcium/calmodulin-dependent protein kinase II-positive principal excitatory and inhibitory neurons of the rat forebrain. J Comp Neurol, 2006. 498(3): 317–29.

12. Steward, O., C.S. Wallace, G.L. Lyford and P.F. Worley, Synaptic activation causes the mRNA for the IEG Arc to localize selectively near activated postsynaptic sites on dendrites. Neuron, 1998. 21(4): 741–51.

13. Steward, O. and P.F. Worley, Selective targeting of newly synthesized Arc mRNA to active synapses requires NMDA receptor activation. Neuron, 2001. 30(1): 227–40.

14. Chowdhury, S., J.D. Shepherd, H. Okuno, G. Lyford, R.S. Petralia, N. Plath, D. Kuhl, R.L. Huganir and P.F. Worley, Arc/Arg3.1 interacts with the endocytic machinery to regulate AMPA receptor trafficking. Neuron, 2006. 52(3): 445–59.

15. Shepherd, J.D. and M.F. Bear, New views of Arc, a master regulator of synaptic plasticity. Nat Neurosci, 2011. 14(3): 279–84.

16. Okuno, H., K. Minatohara and H. Bito, Inverse synaptic tagging: An inactive synapse-specific mechanism to capture activity-induced Arc/arg3.1 and to locally regulate spatial distribution of synaptic weights. Semin Cell Dev Biol, 2018. 77: 43–50.

17. Shepherd, J.D., G. Rumbaugh, J. Wu, S. Chowdhury, N. Plath, D. Kuhl, R.L. Huganir and P.F. Worley, Arc/Arg3.1 mediates homeostatic synaptic scaling of AMPA receptors. Neuron, 2006. 52(3): 475–84.

18. Gao, M., K. Sossa, L. Song, L. Errington, L. Cummings, H. Hwang, D. Kuhl, P. Worley and H.K. Lee, A specific requirement of Arc/Arg3.1 for visual experience-induced homeostatic synaptic plasticity in mouse primary visual cortex. J Neurosci, 2010. 30(21): 7168–78.

19. Beique, J.C., Y. Na, D. Kuhl, P.F. Worley and R.L. Huganir, Arc-dependent synapse-specific homeostatic plasticity. Proc Natl Acad Sci U S A, 2011. 108(2): 816–21.

20. Bloomer, W.A., H.M. VanDongen and A.M. VanDongen, Activity-regulated cytoskeleton-associated protein Arc/Arg3.1 binds to spectrin and associates with nuclear promyelocytic leukemia (PML) bodies. Brain Res, 2007. 1153: 20–33.

21. Bloomer, W.A., H.M. VanDongen and A.M. VanDongen, Arc/Arg3.1 translation is controlled by convergent N-methyl-D-aspartate and Gs-coupled receptor signaling pathways. J Biol Chem, 2008. 283(1): 582–92.

22. Oey, N.E., H.W. Leung, R. Ezhilarasan, L. Zhou, R.W. Beuerman, H.M.A. VanDongen and A.M.J. VanDongen, A Neuronal Activity-Dependent Dual Function Chromatin-Modifying Complex Regulates Arc Expression. eNeuro, 2015. 2(1): ENEURO.0020-14.2015.

23. Torok, D., R.W. Ching and D.P. Bazett-Jones, PML nuclear bodies as sites of epigenetic regulation. Front Biosci (Landmark Ed), 2009. 14: 1325–36.

24. Korb, E., C.L. Wilkinson, R.N. Delgado, K.L. Lovero and S. Finkbeiner, Arc in the nucleus regulates PML-dependent GluA1 transcription and homeostatic plasticity. Nat Neurosci, 2013. 16(7): 874–83.

25. Wee, C.L., S. Teo, N.E. Oey, G.D. Wright, H.M. VanDongen and A.M. VanDongen, Nuclear Arc Interacts with the Histone Acetyltransferase Tip60 to Modify H4K12 Acetylation. eNeuro, 2014. 1(1): ENEURO.0019-14.2014.

26. Qi, D., H. Jin, T. Lilja and M. Mannervik, Drosophila Reptin and other TIP60 complex components promote generation of silent chromatin. Genetics, 2006. 174(1): 241–51.

27. Tea, J.S. and L. Luo, The chromatin remodeling factor Bap55 functions through the TIP60 complex to regulate olfactory projection neuron dendrite targeting. Neural Dev, 2011. 6: 5.

28. Rust, K., M.D. Tiwari, V.K. Mishra, F. Grawe and A. Wodarz, Myc and the Tip60 chromatin remodeling complex control neuroblast maintenance and polarity in Drosophila. EMBO J, 2018. 37(16).

29. Peleg, S., et al., Altered histone acetylation is associated with age-dependent memory impairment in mice. Science, 2010. 328(5979): 753–6.

30. Park, A.Y., Y.S. Park, D. So, I.K. Song, J.E. Choi, H.J. Kim and K.J. Lee, Activity-Regulated Cytoskeleton-Associated Protein (Arc/Arg3.1) is Transiently Expressed after Heat Shock Stress and Suppresses Heat Shock Factor 1. Sci Rep, 2019. 9(1): 2592.

31. Salery, M., M. Dos Santos, E. Saint-Jour, L. Moumne, C. Pages, V. Kappes, S. Parnaudeau, J. Caboche and P. Vanhoutte, Activity-Regulated Cytoskeleton-Associated Protein Accumulates in the Nucleus in Response to Cocaine and Acts as a Brake on Chromatin Remodeling and Long-Term Behavioral Alterations. Biol Psychiatry, 2017. 81(7): 573–584.

32. Hardingham, G.E., F.J. Arnold and H. Bading, Nuclear calcium signaling controls CREB-mediated gene expression triggered by synaptic activity. Nat Neurosci, 2001. 4(3): 261–7.

33. Otmakhov, N., L. Khibnik, N. Otmakhova, S. Carpenter, S. Riahi, B. Asrican and J. Lisman, Forskolin-induced LTP in the CA1 hippocampal region is NMDA receptor dependent. J Neurophysiol, 2004. 91(5): 1955–62.

34. Duvarci, S., K. Nader and J.E. LeDoux, De novo mRNA synthesis is required for both consolidation and reconsolidation of fear memories in the amygdala. Learn Mem, 2008. 15(10): 747–55.

35. Pereira, L.M., C.M. de Castro, L.T.L. Guerra, T.M. Queiroz, J.T. Marques and G.S. Pereira, Hippocampus and Prefrontal Cortex Modulation of Contextual Fear Memory Is Dissociated by Inhibiting De Novo Transcription During Late Consolidation. Mol Neurobiol, 2019.

36. Deisseroth, K., H. Bito and R.W. Tsien, Signaling from synapse to nucleus: postsynaptic CREB phosphorylation during multiple forms of hippocampal synaptic plasticity. Neuron, 1996. 16(1): 89–101.

37. Thompson, K.R., K.O. Otis, D.Y. Chen, Y. Zhao, T.J. O’Dell and K.C. Martin, Synapse to nucleus signaling during long-term synaptic plasticity; a role for the classical active nuclear import pathway. Neuron, 2004. 44(6): 997–1009.

38. Dieterich, D.C., et al., Caldendrin-Jacob: a protein liaison that couples NMDA receptor signalling to the nucleus. PLoS Biol, 2008. 6(2): e34.

39. Kawashima, T., H. Okuno, M. Nonaka, A. Adachi-Morishima, N. Kyo, M. Okamura, S. Takemoto-Kimura, P.F. Worley and H. Bito, Synaptic activity-responsive element in the Arc/Arg3.1 promoter essential for synapse-to-nucleus signaling in activated neurons. Proc Natl Acad Sci U S A, 2009. 106(1): 316–21.

40. Bading, H., Nuclear calcium signalling in the regulation of brain function. Nat Rev Neurosci, 2013. 14(9): 593–608.

41. Tweedie-Cullen, R.Y., J.M. Reck and I.M. Mansuy, Comprehensive mapping of post-translational modifications on synaptic, nuclear, and histone proteins in the adult mouse brain. J Proteome Res, 2009. 8(11): 4966–82.

42. Kim, S. and B.K. Kaang, Epigenetic regulation and chromatin remodeling in learning and memory. Exp Mol Med, 2017. 49(1): e281.

43. Almassalha, L.M., et al., The Global Relationship between Chromatin Physical Topology, Fractal Structure, and Gene Expression. Sci Rep, 2017. 7: 41061.

44. Gottesfeld, J.M. and M.F. Carey, Introduction to the Thematic Minireview Series: Chromatin and transcription. J Biol Chem, 2018. 293(36): 13775–13777.

45. Levenson, J.M., K.J. O’Riordan, K.D. Brown, M.A. Trinh, D.L. Molfese and J.D. Sweatt, Regulation of histone acetylation during memory formation in the hippocampus. J Biol Chem, 2004. 279(39): 40545–59.

46. Korzus, E., M.G. Rosenfeld and M. Mayford, CBP histone acetyltransferase activity is a critical component of memory consolidation. Neuron, 2004. 42(6): 961–72.

47. Wood, M.A., J.D. Hawk and T. Abel, Combinatorial chromatin modifications and memory storage: a code for memory? Learn Mem, 2006. 13(3): 241–4.

48. Graff, J. and L.H. Tsai, Histone acetylation: molecular mnemonics on the chromatin. Nat Rev Neurosci, 2013. 14(2): 97–111.

49. Fischer, A., Epigenetic memory: the Lamarckian brain. EMBO J, 2014. 33(9): 945–67.

50. Zovkic, I.B., B.S. Paulukaitis, J.J. Day, D.M. Etikala and J.D. Sweatt, Histone H2A.Z subunit exchange controls consolidation of recent and remote memory. Nature, 2014. 515(7528): 582–6.

51. McNally, A.G., S.G. Poplawski, B.A. Mayweather, K.M. White and T. Abel, Characterization of a Novel Chromatin Sorting Tool Reveals Importance of Histone Variant H3.3 in Contextual Fear Memory and Motor Learning. Front Mol Neurosci, 2016. 9: 11.

52. Collins, B.E., C.B. Greer, B.C. Coleman and J.D. Sweatt, Histone H3 lysine K4 methylation and its role in learning and memory. Epigenetics Chromatin, 2019. 12(1): 7.

53. Deng, W. and G.A. Blobel, Do chromatin loops provide epigenetic gene expression states? Curr Opin Genet Dev, 2010. 20(5): 548–54.

54. Gondor, A., Dynamic chromatin loops bridge health and disease in the nuclear landscape. Semin Cancer Biol, 2013. 23(2): 90–8.

55. Hansen, A.S., C. Cattoglio, X. Darzacq and R. Tjian, Recent evidence that TADs and chromatin loops are dynamic structures. Nucleus, 2018. 9(1): 20–32.

56. Janssen, K.A., S. Sidoli and B.A. Garcia, Recent Achievements in Characterizing the Histone Code and Approaches to Integrating Epigenomics and Systems Biology. Methods Enzymol, 2017. 586: 359–378.

57. Creyghton, M.P., et al., Histone H3K27ac separates active from poised enhancers and predicts developmental state. Proc Natl Acad Sci U S A, 2010. 107(50): 21931–6.

58. Zhang, B., et al., A dynamic H3K27ac signature identifies VEGFA-stimulated endothelial enhancers and requires EP300 activity. Genome Res, 2013. 23(6): 917–27.

59. Deb, M., S. Kar, D. Sengupta, A. Shilpi, S. Parbin, S.K. Rath, V.A. Londhe and S.K. Patra, Chromatin dynamics: H3K4 methylation and H3 variant replacement during development and in cancer. Cell Mol Life Sci, 2014. 71(18): 3439–63.

60. Esnault, C., F. Gualdrini, S. Horswell, G. Kelly, A. Stewart, P. East, N. Matthews and R. Treisman, ERK-Induced Activation of TCF Family of SRF Cofactors Initiates a Chromatin Modification Cascade Associated with Transcription. Mol Cell, 2017. 65(6): 1081–1095 e5.

61. Rust, M.J., M. Bates and X. Zhuang, Sub-diffraction-limit imaging by stochastic optical reconstruction microscopy (STORM). Nat Methods, 2006. 3(10): 793–5.

62. Ahmed, S. and J.H. Brickner, Regulation and epigenetic control of transcription at the nuclear periphery. Trends Genet, 2007. 23(8): 396–402.

63. Kalverda, B., M.D. Röling and M. Fornerod, Chromatin organization in relation to the nuclear periphery. FEBS Letters, 2008. 582(14): 2017–2022.

64. Fu, M. and Y. Zuo, Experience-dependent structural plasticity in the cortex. Trends Neurosci, 2011. 34(4): 177–87.

65. Holtmaat, A. and P. Caroni, Functional and structural underpinnings of neuronal assembly formation in learning. Nat Neurosci, 2016. 19(12): 1553–1562.

66. Shah, M.M., R.S. Hammond and D.A. Hoffman, Dendritic ion channel trafficking and plasticity. Trends Neurosci, 2010. 33(7): 307–16.

67. Scheefhals, N. and H.D. MacGillavry, Functional organization of postsynaptic glutamate receptors. Mol Cell Neurosci, 2018. 91: 82–94.

68. van Kesteren, R.E. and G.E. Spencer, The role of neurotransmitters in neurite outgrowth and synapse formation. Rev Neurosci, 2003. 14(3): 217–31.

69. McCann, R.F. and D.A. Ross, A Fragile Balance: Dendritic Spines, Learning, and Memory. Biol Psychiatry, 2017. 82(2): e11–e13.

70. Abraham, W.C., O.D. Jones and D.L. Glanzman, Is plasticity of synapses the mechanism of long-term memory storage? npj Science of Learning, 2019. 4(1): 9.

71. Bloom, G.S., Amyloid-beta and tau: the trigger and bullet in Alzheimer disease pathogenesis. JAMA Neurol, 2014. 71(4): 505–8.

72. Buniello, A., et al., The NHGRI-EBI GWAS Catalog of published genome-wide association studies, targeted arrays and summary statistics 2019. Nucleic Acids Res, 2019. 47(D1): D1005–D1012.

73. Harold, D., et al., Genome-wide association study identifies variants at CLU and PICALM associated with Alzheimer’s disease. Nat Genet, 2009. 41(10): 1088–93.

74. Jansen, I.E., et al., Genome-wide meta-analysis identifies new loci and functional pathways influencing Alzheimer’s disease risk. Nat Genet, 2019. 51(3): 404–413.

75. Jun, G., et al., A novel Alzheimer disease locus located near the gene encoding tau protein. Mol Psychiatry, 2016. 21(1): 108–17.

76. Coon, K.D., et al., A high-density whole-genome association study reveals that APOE is the major susceptibility gene for sporadic late-onset Alzheimer’s disease. J Clin Psychiatry, 2007. 68(4): 613–8.

77. Lambert, J.C., et al., Meta-analysis of 74,046 individuals identifies 11 new susceptibility loci for Alzheimer’s disease. Nat Genet, 2013. 45(12): 1452–8.

78. Nikolaienko, O., S. Patil, M.S. Eriksen and C.R. Bramham, Arc protein: a flexible hub for synaptic plasticity and cognition. Semin Cell Dev Biol, 2018. 77: 33–42.

79. Gozdz, A., O. Nikolaienko, M. Urbanska, I.A. Cymerman, E. Sitkiewicz, M. Blazejczyk, M. Dadlez, C.R. Bramham and J. Jaworski, GSK3alpha and GSK3beta Phosphorylate Arc and Regulate its Degradation. Front Mol Neurosci, 2017. 10: 192.

80. Wu, J., et al., Arc/Arg3.1 regulates an endosomal pathway essential for activity-dependent beta-amyloid generation. Cell, 2011. 147(3): 615–28.

81. Huang da, W., B.T. Sherman and R.A. Lempicki, Systematic and integrative analysis of large gene lists using DAVID bioinformatics resources. Nat Protoc, 2009. 4(1): 44–57.

82. Wang, Y.T. and D.J. Linden, Expression of cerebellar long-term depression requires postsynaptic clathrin-mediated endocytosis. Neuron, 2000. 25(3): 635–47.

83. Fowler, C.J., The contribution of cyclooxygenase-2 to endocannabinoid metabolism and action. Br J Pharmacol, 2007. 152(5): 594–601.

84. Aid, S. and F. Bosetti, Targeting cyclooxygenases-1 and −2 in neuroinflammation: Therapeutic implications. Biochimie, 2011. 93(1): 46–51.

85. Ma, S.L., N.L. Tang, Y.P. Zhang, L.D. Ji, C.W. Tam, V.W. Lui, H.F. Chiu and L.C. Lam, Association of prostaglandin-endoperoxide synthase 2 (PTGS2) polymorphisms and Alzheimer’s disease in Chinese. Neurobiol Aging, 2008. 29(6): 856–60.

86. Woodling, N.S., et al., Cyclooxygenase inhibition targets neurons to prevent early behavioural decline in Alzheimer’s disease model mice. Brain, 2016. 139(Pt 7): 2063–81.

87. Raivich, G., c-Jun expression, activation and function in neural cell death, inflammation and repair. J Neurochem, 2008. 107(4): 898–906.

88. Savage, M.J., Y.G. Lin, J.R. Ciallella, D.G. Flood and R.W. Scott, Activation of c-Jun N-terminal kinase and p38 in an Alzheimer’s disease model is associated with amyloid deposition. J Neurosci, 2002. 22(9): 3376–85.

89. Yarza, R., S. Vela, M. Solas and M.J. Ramirez, c-Jun N-terminal Kinase (JNK) Signaling as a Therapeutic Target for Alzheimer’s Disease. Front Pharmacol, 2015. 6: 321.

90. Jiang, C., X. Zou, R. Zhu, Y. Shi, Z. Wu, F. Zhao and L. Chen, The correlation between accumulation of amyloid beta with enhanced neuroinflammation and cognitive impairment after intraventricular hemorrhage. Journal of Neurosurgery, 2018. 131(1): 54–63.

91. Broadhead, M.J., et al., PSD95 nanoclusters are postsynaptic building blocks in hippocampus circuits. Sci Rep, 2016. 6: 24626.

92. Craven, S.E., A.E. El-Husseini and D.S. Bredt, Synaptic targeting of the postsynaptic density protein PSD-95 mediated by lipid and protein motifs. Neuron, 1999. 22(3): 497–509.

93. Medrihan, L., F. Cesca, A. Raimondi, G. Lignani, P. Baldelli and F. Benfenati, Synapsin II desynchronizes neurotransmitter release at inhibitory synapses by interacting with presynaptic calcium channels. Nat Commun, 2013. 4: 1512.

94. Wang, Y., D. Feng, G. Liu, Q. Luo, Y. Xu, S. Lin, J. Fei and L. Xu, Gamma-aminobutyric acid transporter 1 negatively regulates T cell-mediated immune responses and ameliorates autoimmune inflammation in the CNS. J Immunol, 2008. 181(12): 8226–36.

95. Bodnar, R.J., Endogenous Opiates and Behavior: 2016. Peptides, 2018. 101: 167–212.

96. Kaplan, G.B., K.A. Leite-Morris, W. Fan, A.J. Young and M.D. Guy, Opiate sensitization induces FosB/DeltaFosB expression in prefrontal cortical, striatal and amygdala brain regions. PLoS One, 2011. 6(8): e23574.

97. Zhang, C.J., et al., TLR-stimulated IRAKM activates caspase-8 inflammasome in microglia and promotes neuroinflammation. J Clin Invest, 2018. 128(12): 5399–5412.

98. Crowley, T., J.F. Cryan, E.J. Downer and O.F. O’Leary, Inhibiting neuroinflammation: The role and therapeutic potential of GABA in neuro-immune interactions. Brain Behav Immun, 2016. 54: 260–277.

99. Haerian, B.S., et al., Contribution of GABRG2 Polymorphisms to Risk of Epilepsy and Febrile Seizure: a Multicenter Cohort Study and Meta-analysis. Mol Neurobiol, 2016. 53(8): 5457–67.

100. Ninan, I., Synaptic regulation of affective behaviors; role of BDNF. Neuropharmacology, 2014. 76 Pt C: 684–95.

101. Cunha, C., R. Brambilla and K.L. Thomas, A simple role for BDNF in learning and memory? Front Mol Neurosci, 2010. 3: 1.

102. Han, R., Z. Liu, N. Sun, S. Liu, L. Li, Y. Shen, J. Xiu and Q. Xu, BDNF Alleviates Neuroinflammation in the Hippocampus of Type 1 Diabetic Mice via Blocking the Aberrant HMGB1/RAGE/NF-kappaB Pathway. Aging Dis, 2019. 10(3): 611–625.

103. Giuffrida, M.L., A. Copani and E. Rizzarelli, A promising connection between BDNF and Alzheimer’s disease. Aging (Albany NY), 2018. 10(8): 1791–1792.

104. Charbogne, P., B.L. Kieffer and K. Befort, 15 years of genetic approaches in vivo for addiction research: Opioid receptor and peptide gene knockout in mouse models of drug abuse. Neuropharmacology, 2014. 76 Pt B: 204–17.

105. Zhang, Y., et al., CRISPR/gRNA-directed synergistic activation mediator (SAM) induces specific, persistent and robust reactivation of the HIV-1 latent reservoirs. Sci Rep, 2015. 5: 16277.

106. Seiliez, I., B. Thisse and C. Thisse, FoxA3 and goosecoid promote anterior neural fate through inhibition of Wnt8a activity before the onset of gastrulation. Dev Biol, 2006. 290(1): 152–63.

107. Dal-Pra, S., C. Thisse and B. Thisse, FoxA transcription factors are essential for the development of dorsal axial structures. Dev Biol, 2011. 350(2): 484–95.

108. Hsieh, H., J. Boehm, C. Sato, T. Iwatsubo, T. Tomita, S. Sisodia and R. Malinow, AMPAR removal underlies Abeta-induced synaptic depression and dendritic spine loss. Neuron, 2006. 52(5): 831–43.

109. Knobloch, M. and I.M. Mansuy, Dendritic spine loss and synaptic alterations in Alzheimer’s disease. Mol Neurobiol, 2008. 37(1): 73–82.

110. Alberi, L., et al., Activity-induced Notch signaling in neurons requires Arc/Arg3.1 and is essential for synaptic plasticity in hippocampal networks. Neuron, 2011. 69(3): 437–44.

111. Zhang, P., Y. Zhao and X.-H. Sun, Notch-Regulated Periphery B Cell Differentiation Involves Suppression of E Protein Function. The Journal of Immunology, 2013. 191(2): 726–736.

112. Choi, Y.-H., E.-J. Ann, J.-H. Yoon, J.-S. Mo, M.-Y. Kim and H.-S. Park, Calcium/calmodulin-dependent protein kinase IV (CaMKIV) enhances osteoclast differentiation via the up-regulation of Notch1 protein stability. Biochimica et Biophysica Acta (BBA) - Molecular Cell Research, 2013. 1833(1): 69–79.

113. Wang, D., J. Long, F. Dai, M. Liang, X.-H. Feng and X. Lin, BCL6 Represses Smad Signaling in Transforming Growth Factor-β Resistance. Cancer Research, 2008. 68(3): 783–789.

114. Zhao, X., A. Ito, C.D. Kane, T.-S. Liao, T.A. Bolger, S.M. Lemrow, A.R. Means and T.-P. Yao, The Modular Nature of Histone Deacetylase HDAC4 Confers Phosphorylation-dependent Intracellular Trafficking. Journal of Biological Chemistry, 2001. 276(37): 35042–35048.

115. Graziani, I., S. Eliasz, M.A. De Marco, Y. Chen, H.I. Pass, R.M. De May, P.R. Strack, L. Miele and M. Bocchetta, Opposite effects of Notch-1 and Notch-2 on mesothelioma cell survival under hypoxia are exerted through the Akt pathway. Cancer Res, 2008. 68(23): 9678–85.

116. Fang, L., L. Zhang, W. Wei, X. Jin, P. Wang, Y. Tong, J. Li, J.X. Du and J. Wong, A methylation-phosphorylation switch determines Sox2 stability and function in ESC maintenance or differentiation. Mol Cell, 2014. 55(4): 537–51.

117. Jeong, C.H., et al., Phosphorylation of Sox2 cooperates in reprogramming to pluripotent stem cells. Stem Cells, 2010. 28(12): 2141–50.

118. Zhou, L., N. Zhang, W. Song, N. You, Q. Li, W. Sun, Y. Zhang, D. Wang and K. Dou, The significance of Notch1 compared with Notch3 in high metastasis and poor overall survival in hepatocellular carcinoma. PLoS One, 2013. 8(2): e57382.

119. Yu, X.-M., et al., Notch1 Signaling Regulates the Aggressiveness of Differentiated Thyroid Cancer and Inhibits SERPINE1 Expression. Clinical Cancer Research, 2016. 22(14): 3582–3592.

120. Dekker, R.J., H. Pannekoek and A.J.G. Horrevoets, A steady-state competition model describes the modulating effects of thrombomodulin on thrombin inhibition by plasminogen activator inhibitor-1 in the absence and presence of vitronectin. European Journal of Biochemistry, 2003. 270(9): 1942–1951.

121. Idogawa, M., M. Masutani, M. Shitashige, K. Honda, T. Tokino, Y. Shinomura, K. Imai, S. Hirohashi and T. Yamada, Ku70 and Poly(ADP-Ribose) Polymerase-1 Competitively Regulate β-Catenin and T-Cell Factor-4–Mediated Gene Transactivation: Possible Linkage of DNA Damage Recognition and Wnt Signaling. Cancer Research, 2007. 67(3): 911–918.

122. Hicks, C., A. Pannuti and L. Miele, Associating GWAS Information with the Notch Signaling Pathway Using Transcription Profiling. Cancer Informatics, 2011. 10: CIN.S6072.

123. Jehn, B.M., W. Bielke, W.S. Pear and B.A. Osborne, Cutting edge: protective effects of notch-1 on TCR-induced apoptosis. J Immunol, 1999. 162(2): 635–8.

124. Kurakula, K., M. Vos, A. Logiantara, J.J. Roelofs, M.A. Nieuwenhuis, G.H. Koppelman, D.S. Postma, L.S. van Rijt and C.J. de Vries, Nuclear Receptor Nur77 Attenuates Airway Inflammation in Mice by Suppressing NF-kappaB Activity in Lung Epithelial Cells. J Immunol, 2015. 195(4): 1388–98.

125. Sade, H., S. Krishna and A. Sarin, The anti-apoptotic effect of Notch-1 requires p56lck-dependent, Akt/PKB-mediated signaling in T cells. J Biol Chem, 2004. 279(4): 2937–44.

126. Yamaguchi, N. and N. Yamaguchi, The seventh zinc finger motif of A20 is required for the suppression of TNF-alpha-induced apoptosis. FEBS Lett, 2015. 589(12): 1369–75.

127. Tapias, A. and Z.Q. Wang, Lysine Acetylation and Deacetylation in Brain Development and Neuropathies. Genomics Proteomics Bioinformatics, 2017. 15(1): 19–36.

128. Xu, S., R. Wilf, T. Menon, P. Panikker, J. Sarthi and F. Elefant, Epigenetic control of learning and memory in Drosophila by Tip60 HAT action. Genetics, 2014. 198(4): 1571–86.

129. Pirooznia, S.K., J. Sarthi, A.A. Johnson, M.S. Toth, K. Chiu, S. Koduri and F. Elefant, Tip60 HAT activity mediates APP induced lethality and apoptotic cell death in the CNS of a Drosophila Alzheimer’s disease model. PLoS One, 2012. 7(7): e41776.

130. Jang, S.H., H. Chen, P.K. Gregersen, B. Diamond and S.J. Kim, Kruppel-like factor4 regulates PRDM1 expression through binding to an autoimmune risk allele. JCI Insight, 2017. 2(1).

131. Cui, H., M. Guo, D. Xu, Z.C. Ding, G. Zhou, H.F. Ding, J. Zhang, Y. Tang and C. Yan, The stress-responsive gene ATF3 regulates the histone acetyltransferase Tip60. Nat Commun, 2015. 6: 6752.

132. Cui, H., X. Li, C. Han, Q.E. Wang, H. Wang, H.F. Ding, J. Zhang and C. Yan, The Stress-responsive Gene ATF3 Mediates Dichotomous UV Responses by Regulating the Tip60 and p53 Proteins. J Biol Chem, 2016. 291(20): 10847–57.

133. Darlyuk-Saadon, I., K. Weidenfeld-Baranboim, K.K. Yokoyama, T. Hai and A. Aronheim, The bZIP repressor proteins, c-Jun dimerization protein 2 and activating transcription factor 3, recruit multiple HDAC members to the ATF3 promoter. Biochim Biophys Acta, 2012. 1819(11-12): 1142–53.

134. Bolger, T.A., X. Zhao, T.J. Cohen, C.C. Tsai and T.P. Yao, The neurodegenerative disease protein ataxin-1 antagonizes the neuronal survival function of myocyte enhancer factor-2. J Biol Chem, 2007. 282(40): 29186–92.

135. Serra, H.G., et al., RORα-Mediated Purkinje Cell Development Determines Disease Severity in Adult SCA1 Mice. Cell, 2006. 127(4): 697–708.

136. Kim, J.W., S.M. Jang, C.H. Kim, J.H. An, E.J. Kang and K.H. Choi, New molecular bridge between RelA/p65 and NF-kappaB target genes via histone acetyltransferase TIP60 cofactor. J Biol Chem, 2012. 287(10): 7780–91.

137. Koh, D.I., et al., KAISO, a critical regulator of p53-mediated transcription of CDKN1A and apoptotic genes. Proc Natl Acad Sci U S A, 2014. 111(42): 15078–83.

138. Del Valle-Perez, B., D. Casagolda, E. Lugilde, G. Valls, M. Codina, N. Dave, A.G. de Herreros and M. Dunach, Wnt controls the transcriptional activity of Kaiso through CK1epsilon-dependent phosphorylation of p120-catenin. J Cell Sci, 2011. 124(Pt 13): 2298–309.

139. Nagl, N.G., Jr., X. Wang, A. Patsialou, M. Van Scoy and E. Moran, Distinct mammalian SWI/SNF chromatin remodeling complexes with opposing roles in cell-cycle control. EMBO J, 2007. 26(3): 752–63.

140. Kim, B.R., E. Coyaud, E.M.N. Laurent, J. St-Germain, E. Van de Laar, M.S. Tsao, B. Raught and N. Moghal, Identification of the SOX2 Interactome by BioID Reveals EP300 as a Mediator of SOX2-dependent Squamous Differentiation and Lung Squamous Cell Carcinoma Growth. Mol Cell Proteomics, 2017. 16(10): 1864–1888.

141. Singhal, N., J. Graumann, G. Wu, M.J. Arauzo-Bravo, D.W. Han, B. Greber, L. Gentile, M. Mann and H.R. Scholer, Chromatin-Remodeling Components of the BAF Complex Facilitate Reprogramming. Cell, 2010. 141(6): 943–55.

142. Krolewski, R.C., A. Packard, W. Jang, H. Wildner and J.E. Schwob, Ascl1 (Mash1) knockout perturbs differentiation of nonneuronal cells in olfactory epithelium. PLoS One, 2012. 7(12): e51737.

143. Borromeo, M.D., D.M. Meredith, D.S. Castro, J.C. Chang, K.C. Tung, F. Guillemot and J.E. Johnson, A transcription factor network specifying inhibitory versus excitatory neurons in the dorsal spinal cord. Development, 2014. 141(14): 2803–12.

144. Brouillette, J., R. Caillierez, N. Zommer, C. Alves-Pires, I. Benilova, D. Blum, B. De Strooper and L. Buee, Neurotoxicity and memory deficits induced by soluble low-molecular-weight amyloid-beta1-42 oligomers are revealed in vivo by using a novel animal model. J Neurosci, 2012. 32(23): 7852–61.

145. Paroni, G., M. Mizzau, C. Henderson, G. Del Sal, C. Schneider and C. Brancolini, Caspase-dependent regulation of histone deacetylase 4 nuclear-cytoplasmic shuttling promotes apoptosis. Mol Biol Cell, 2004. 15(6): 2804–18.

146. Chang, C.C., et al., Connective tissue growth factor activates pluripotency genes and mesenchymal-epithelial transition in head and neck cancer cells. Cancer Res, 2013. 73(13): 4147–57.

147. Medeiros, R., et al., Connecting TNF-alpha signaling pathways to iNOS expression in a mouse model of Alzheimer’s disease: relevance for the behavioral and synaptic deficits induced by amyloid beta protein. J Neurosci, 2007. 27(20): 5394–404.

148. Manns, J., M. Rico, L.L. Mason and D.L.C.A. Raul, Thrombospondin-1 (TSP1) Promotes Thrombin Generation on the Surface of Fibroblasts (HS-68) and Induces Up-Regulation of Connective Tissue Growth Factor (CTGF) Gene and Protein Expression. Blood, 2006. 108(11): 1755–1755.

149. Son, S.M., D.W. Nam, M.Y. Cha, K.H. Kim, J. Byun, H. Ryu and I. Mook-Jung, Thrombospondin-1 prevents amyloid beta-mediated synaptic pathology in Alzheimer’s disease. Neurobiol Aging, 2015. 36(12): 3214–3227.

150. Cao, X. and T.C. Sudhof, A transcriptionally active complex of APP with Fe65 and histone acetyltransferase Tip60. Science, 2001. 293(5527): 115–20.

151. Del Prete, D., R.C. Rice, A.M. Rajadhyaksha and L. D’Adamio, Amyloid Precursor Protein (APP) May Act as a Substrate and a Recognition Unit for CRL4CRBN and Stub1 E3 Ligases Facilitating Ubiquitination of Proteins Involved in Presynaptic Functions and Neurodegeneration. J Biol Chem, 2016. 291(33): 17209–27.

152. Fritsch, J., M. Stephan, V. Tchikov, S. Winoto-Morbach, S. Gubkina, D. Kabelitz and S. Schutze, Cell fate decisions regulated by K63 ubiquitination of tumor necrosis factor receptor 1. Mol Cell Biol, 2014. 34(17): 3214–28.

153. Marblestone, J.G., S. Butt, D.M. McKelvey, D.E. Sterner, M.R. Mattern, B. Nicholson and M.J. Eddins, Comprehensive ubiquitin E2 profiling of ten ubiquitin E3 ligases. Cell Biochem Biophys, 2013. 67(1): 161–7.

154. Belova, L., S. Sharma, D.R. Brickley, J.R. Nicolarsen, C. Patterson and S.D. Conzen, Ubiquitin-proteasome degradation of serum- and glucocorticoid-regulated kinase-1 (SGK-1) is mediated by the chaperone-dependent E3 ligase CHIP. Biochem J, 2006. 400(2): 235–44.

155. Bogusz, A.M., D.R. Brickley, T. Pew and S.D. Conzen, A novel N-terminal hydrophobic motif mediates constitutive degradation of serum- and glucocorticoid-induced kinase-1 by the ubiquitin-proteasome pathway. FEBS J, 2006. 273(13): 2913–28.

156. Gao, D., et al., Rictor forms a complex with Cullin-1 to promote SGK1 ubiquitination and destruction. Mol Cell, 2010. 39(5): 797–808.

157. Mishra, A., P. Dikshit, S. Purkayastha, J. Sharma, N. Nukina and N.R. Jana, E6-AP promotes misfolded polyglutamine proteins for proteasomal degradation and suppresses polyglutamine protein aggregation and toxicity. J Biol Chem, 2008. 283(12): 7648–56.

158. Li, F., T. Macfarlan, R.N. Pittman and D. Chakravarti, Ataxin-3 Is a Histone-binding Protein with Two Independent Transcriptional Corepressor Activities. Journal of Biological Chemistry, 2002. 277(47): 45004–45012.

159. David, S. and R.G. Kalb, Serum/glucocorticoid-inducible kinase can phosphorylate the cyclic AMP response element binding protein, CREB. FEBS Lett, 2005. 579(6): 1534–8.

160. Gundersen, B.B., L.A. Briand, J.L. Onksen, J. Lelay, K.H. Kaestner and J.A. Blendy, Increased hippocampal neurogenesis and accelerated response to antidepressants in mice with specific deletion of CREB in the hippocampus: role of cAMP response-element modulator tau. J Neurosci, 2013. 33(34): 13673–85.

161. Hummler, E., T.J. Cole, J.A. Blendy, R. Ganss, A. Aguzzi, W. Schmid, F. Beermann and G. Schutz, Targeted mutation of the CREB gene: compensation within the CREB/ATF family of transcription factors. Proc Natl Acad Sci U S A, 1994. 91(12): 5647–51.

162. Zhang, J.W., D.J. Klemm, C. Vinson and M.D. Lane, Role of CREB in transcriptional regulation of CCAAT/enhancer-binding protein beta gene during adipogenesis. J Biol Chem, 2004. 279(6): 4471–8.

163. Bohn, T., et al., Tumor immunoevasion via acidosis-dependent induction of regulatory tumor-associated macrophages. Nat Immunol, 2018. 19(12): 1319–1329.

164. Kibbey, M.C., M. Jucker, B.S. Weeks, R.L. Neve, W.E. Van Nostrand and H.K. Kleinman, beta-Amyloid precursor protein binds to the neurite-promoting IKVAV site of laminin. Proc Natl Acad Sci U S A, 1993. 90(21): 10150–3.

165. Adair-Kirk, T.L., J.J. Atkinson, D.G. Kelley, R.H. Arch, J.H. Miner and R.M. Senior, A chemotactic peptide from laminin alpha 5 functions as a regulator of inflammatory immune responses via TNF alpha-mediated signaling. J Immunol, 2005. 174(3): 1621–9.

166. Takashima, A., K. Noguchi, K. Sato, T. Hoshino and K. Imahori, Tau protein kinase I is essential for amyloid beta-protein-induced neurotoxicity. Proc Natl Acad Sci U S A, 1993. 90(16): 7789–93.

167. Takashima, A., K. Noguchi, G. Michel, M. Mercken, M. Hoshi, K. Ishiguro and K. Imahori, Exposure of rat hippocampal neurons to amyloid beta peptide (25-35) induces the inactivation of phosphatidyl inositol-3 kinase and the activation of tau protein kinase I/glycogen synthase kinase-3 beta. Neurosci Lett, 1996. 203(1): 33–6.

168. Reifert, J., D. Hartung-Cranston and S.C. Feinstein, Amyloid beta-mediated cell death of cultured hippocampal neurons reveals extensive Tau fragmentation without increased full-length tau phosphorylation. J Biol Chem, 2011. 286(23): 20797–811.

169. Tyson, D.R., J.T. Swarthout, S.C. Jefcoat and N.C. Partridge, PTH induction of transcriptional activity of the cAMP response element-binding protein requires the serine 129 site and glycogen synthase kinase-3 activity, but not casein kinase II sites. Endocrinology, 2002. 143(2): 674–82.

170. Verma, N.K., et al., LFA-1/ICAM-1 Ligation in Human T Cells Promotes Th1 Polarization through a GSK3beta Signaling-Dependent Notch Pathway. J Immunol, 2016. 197(1): 108–18.

171. Pigino, G., G. Morfini, A. Pelsman, M.P. Mattson, S.T. Brady and J. Busciglio, Alzheimer’s presenilin 1 mutations impair kinesin-based axonal transport. J Neurosci, 2003. 23(11): 4499–508.

172. Song, B., B. Lai, Z. Zheng, Y. Zhang, J. Luo, C. Wang, Y. Chen, J.R. Woodgett and M. Li, Inhibitory phosphorylation of GSK-3 by CaMKII couples depolarization to neuronal survival. J Biol Chem, 2010. 285(52): 41122–34.

173. Okuno, H., et al., Inverse synaptic tagging of inactive synapses via dynamic interaction of Arc/Arg3.1 with CaMKIIbeta. Cell, 2012. 149(4): 886–98.

174. Sun, X., S. Sato, O. Murayama, M. Murayama, J.M. Park, H. Yamaguchi and A. Takashima, Lithium inhibits amyloid secretion in COS7 cells transfected with amyloid precursor protein C100. Neurosci Lett, 2002. 321(1-2): 61–4.

175. Pierrot, N., S.F. Santos, C. Feyt, M. Morel, J.P. Brion and J.N. Octave, Calcium-mediated transient phosphorylation of tau and amyloid precursor protein followed by intraneuronal amyloid-beta accumulation. J Biol Chem, 2006. 281(52): 39907–14.

176. Pinnix, I., J.A. Ghiso, M.A. Pappolla and K. Sambamurti, Major carboxyl terminal fragments generated by gamma-secretase processing of the Alzheimer amyloid precursor are 50 and 51 amino acids long. Am J Geriatr Psychiatry, 2013. 21(5): 474–83.

177. von Rotz, R.C., B.M. Kohli, J. Bosset, M. Meier, T. Suzuki, R.M. Nitsch and U. Konietzko, The APP intracellular domain forms nuclear multiprotein complexes and regulates the transcription of its own precursor. Journal of Cell Science, 2004. 117(19): 4435–4448.

178. Muller, T., A. Schrotter, C. Loosse, K. Pfeiffer, C. Theiss, M. Kauth, H.E. Meyer and K. Marcus, A ternary complex consisting of AICD, FE65, and TIP60 down-regulates Stathmin1. Biochim Biophys Acta, 2013. 1834(1): 387–94.

179. Kim, H.S., et al., C-terminal fragments of amyloid precursor protein exert neurotoxicity by inducing glycogen synthase kinase-3beta expression. FASEB J, 2003. 17(13): 1951–3.

180. Hass, M.R. and B.A. Yankner, A {gamma}-secretase-independent mechanism of signal transduction by the amyloid precursor protein. J Biol Chem, 2005. 280(44): 36895–904.

181. Kinoshita, A., C.M. Whelan, O. Berezovska and B.T. Hyman, The gamma secretase-generated carboxyl-terminal domain of the amyloid precursor protein induces apoptosis via Tip60 in H4 cells. J Biol Chem, 2002. 277(32): 28530–6.

182. Sumioka, A., S. Nagaishi, T. Yoshida, A. Lin, M. Miura and T. Suzuki, Role of 14-3-3gamma in FE65-dependent gene transactivation mediated by the amyloid beta-protein precursor cytoplasmic fragment. J Biol Chem, 2005. 280(51): 42364–74.

183. Kim, S.Y., M.Y. Kim, J.S. Mo and H.S. Park, Notch1 intracellular domain suppresses APP intracellular domain-Tip60-Fe65 complex mediated signaling through physical interaction. Biochim Biophys Acta, 2007. 1773(6): 736–46.

184. Fischer, D.F., R. van Dijk, J.A. Sluijs, S.M. Nair, M. Racchi, C.N. Levelt, F.W. van Leeuwen and E.M. Hol, Activation of the Notch pathway in Down syndrome: cross-talk of Notch and APP. FASEB J, 2005. 19(11): 1451–8.

185. Berezovska, O., C. Jack, A. Deng, N. Gastineau, G.W. Rebeck and B. T. Hyman, Notch1 and amyloid precursor protein are competitive substrates for presenilin1-dependent gamma-secretase cleavage. J Biol Chem, 2001. 276(32): 30018–23.

186. Cao, C., M.S. Rioult-Pedotti, P. Migani, C.J. Yu, R. Tiwari, K. Parang, M.R. Spaller, D.J. Goebel and J. Marshall, Impairment of TrkB-PSD-95 signaling in Angelman syndrome. PLoS Biol, 2013. 11(2): e1001478.

187. Nair, R.R., S. Patil, A. Tiron, T. Kanhema, D. Panja, L. Schiro, K. Parobczak, G. Wilczynski and C.R. Bramham, Dynamic Arc SUMOylation and Selective Interaction with F-Actin-Binding Protein Drebrin A in LTP Consolidation In Vivo. Front Synaptic Neurosci, 2017. 9: 8.

188. Pastuzyn, E.D. and J.D. Shepherd, Activity-Dependent Arc Expression and Homeostatic Synaptic Plasticity Are Altered in Neurons from a Mouse Model of Angelman Syndrome. Frontiers in Molecular Neuroscience, 2017. 10(234).

189. Yoshii, A. and M. Constantine-Paton, Postsynaptic BDNF-TrkB signaling in synapse maturation, plasticity, and disease. Dev Neurobiol, 2010. 70(5): 304–22.

190. Martin-Avila, A., et al., Protein Tyrosine Kinase Fyn Regulates TLR4-Elicited Responses on Mast Cells Controlling the Function of a PP2A-PKCalpha/beta Signaling Node Leading to TNF Secretion. J Immunol, 2016. 196(12): 5075–88.

191. Irie, Y., K. Yamagata, Y. Gan, K. Miyamoto, E. Do, C.H. Kuo, E. Taira and N. Miki, Molecular cloning and characterization of Amida, a novel protein which interacts with a neuron-specific immediate early gene product arc, contains novel nuclear localization signals, and causes cell death in cultured cells. J Biol Chem, 2000. 275(4): 2647–53.

192. Cai, Y., et al., YY1 functions with INO80 to activate transcription. Nat Struct Mol Biol, 2007. 14(9): 872–4.

193. Chen, L., S.K. Ooi, R.C. Conaway and J.W. Conaway, Generation and purification of human INO80 chromatin remodeling complexes and subcomplexes. J Vis Exp, 2014(92): e51720.

194. Alberini, C.M. and E.R. Kandel, The regulation of transcription in memory consolidation. Cold Spring Harb Perspect Biol, 2014. 7(1): a021741.

195. Rial Verde, E.M., J. Lee-Osbourne, P.F. Worley, R. Malinow and H.T. Cline, Increased expression of the immediate-early gene arc/arg3.1 reduces AMPA receptor-mediated synaptic transmission. Neuron, 2006. 52(3): 461–74.

196. Waung, M.W., B.E. Pfeiffer, E.D. Nosyreva, J.A. Ronesi and K.M. Huber, Rapid translation of Arc/Arg3.1 selectively mediates mGluR-dependent LTD through persistent increases in AMPAR endocytosis rate. Neuron, 2008. 59(1): 84–97.

197. Park, S., et al., Elongation factor 2 and fragile X mental retardation protein control the dynamic translation of Arc/Arg3.1 essential for mGluR-LTD. Neuron, 2008. 59(1): 70–83.

198. Wang, H., et al., Metabotropic Glutamate Receptors Induce a Form of LTP Controlled by Translation and Arc Signaling in the Hippocampus. J Neurosci, 2016. 36(5): 1723–9.

199. Wilkerson, J.R., J.P. Albanesi and K.M. Huber, Roles for Arc in metabotropic glutamate receptor-dependent LTD and synapse elimination: Implications in health and disease. Semin Cell Dev Biol, 2018. 77: 51–62.

200. Dunn, A.R. and C.C. Kaczorowski, Regulation of intrinsic excitability: Roles for learning and memory, aging and Alzheimer’s disease, and genetic diversity. Neurobiol Learn Mem, 2019. 164: 107069.

201. Mozzachiodi, R., F.D. Lorenzetti, D.A. Baxter and J.H. Byrne, Changes in neuronal excitability serve as a mechanism of long-term memory for operant conditioning. Nat Neurosci, 2008. 11(10): 1146–8.

202. Otis, J.M., M.K. Fitzgerald, H. Yousuf, J.L. Burkard, M. Drake and D. Mueller, Prefrontal Neuronal Excitability Maintains Cocaine-Associated Memory During Retrieval. Front Behav Neurosci, 2018. 12: 119.

203. Scheltens, P., K. Blennow, M.M. Breteler, B. de Strooper, G.B. Frisoni, S. Salloway and W.M. Van der Flier, Alzheimer’s disease. Lancet, 2016.

204. Ballatore, C., V.M. Lee and J.Q. Trojanowski, Tau-mediated neurodegeneration in Alzheimer’s disease and related disorders. Nat Rev Neurosci, 2007. 8(9): 663–72.

205. Kerrigan, T.L. and A.D. Randall, A new player in the “synaptopathy” of Alzheimer’s disease - arc/arg 3.1. Front Neurol, 2013. 4: 9.

206. Palop, J.J. and L. Mucke, Amyloid-beta-induced neuronal dysfunction in Alzheimer’s disease: from synapses toward neural networks. Nat Neurosci, 2010. 13(7): 812–8.

207. Prince, M., R. Bryce, E. Albanese, A. Wimo, W. Ribeiro and C.P. Ferri, The global prevalence of dementia: a systematic review and metaanalysis. Alzheimers Dement, 2013. 9(1): 63–75 e2.

208. Collaborators, G.B.D.D., Global, regional, and national burden of Alzheimer’s disease and other dementias, 1990-2016: a systematic analysis for the Global Burden of Disease Study 2016. Lancet Neurol, 18(1): 88–106.

209. Lacor, P.N., et al., Synaptic targeting by Alzheimer’s-related amyloid beta oligomers. J Neurosci, 2004. 24(45): 10191–200.

210. Rudinskiy, N., J.M. Hawkes, R.A. Betensky, M. Eguchi, S. Yamaguchi, T.L. Spires-Jones and B.T. Hyman, Orchestrated experience-driven Arc responses are disrupted in a mouse model of Alzheimer’s disease. Nat Neurosci, 2012. 15(10): 1422–9.

211. Landgren, S., et al., A novel ARC gene polymorphism is associated with reduced risk of Alzheimer’s disease. Journal of neural transmission, 2012. 119(7): 833–42.

212. Morin, J.P., G. Ceron-Solano, G. Velazquez-Campos, G. Pacheco-Lopez, F. Bermudez-Rattoni and S. Diaz-Cintra, Spatial Memory Impairment is Associated with Intraneural Amyloid-beta Immunoreactivity and Dysfunctional Arc Expression in the Hippocampal-CA3 Region of a Transgenic Mouse Model of Alzheimer’s Disease. J Alzheimers Dis, 2016.

213. Rosi, S., Neuroinflammation and the plasticity-related immediate-early gene Arc. Brain Behav Immun, 2011. 25 Suppl 1: S39–49.

214. Long, J.M. and D.M. Holtzman, Alzheimer Disease: An Update on Pathobiology and Treatment Strategies. Cell, 2019. 179(2): 312–339.

215. Du, X., X. Wang and M. Geng, Alzheimer’s disease hypothesis and related therapies. Transl Neurodegener, 2018. 7: 2.

216. Liu, P.-P., Y. Xie, X.-Y. Meng and J.-S. Kang, History and progress of hypotheses and clinical trials for Alzheimer’s disease. Signal Transduct Targeted Ther, 2019. 4(1): 29.

217. Hara, Y., N. McKeehan and H.M. Fillit, Translating the biology of aging into novel therapeutics for Alzheimer disease. Neurology, 2019. 92(2): 84–93.

218. Pei, Q., T.S. Zetterstrom, M. Sprakes, R. Tordera and T. Sharp, Antidepressant drug treatment induces Arc gene expression in the rat brain. Neuroscience, 2003. 121(4): 975–82.

219. Thomsen, M.S., H.H. Hansen and J.D. Mikkelsen, Opposite effect of phencyclidine on activity-regulated cytoskeleton-associated protein (Arc) in juvenile and adult limbic rat brain regions. Neurochem Int, 2010. 56(2): 270–5.

220. McReynolds, J.R., K. Donowho, A. Abdi, J.L. McGaugh, B. Roozendaal and C.K. McIntyre, Memory-enhancing corticosterone treatment increases amygdala norepinephrine and Arc protein expression in hippocampal synaptic fractions. Neurobiol Learn Mem, 2010. 93(3): 312–21.

221. Judes, G., K. Rifai, M. Ngollo, M. Daures, Y.J. Bignon, F. Penault-Llorca and D. Bernard-Gallon, A bivalent role of TIP60 histone acetyl transferase in human cancer. Epigenomics, 2015. 7(8): 1351–63.

222. Venkatesh, V., R. Nataraj, G.S. Thangaraj, M. Karthikeyan, A. Gnanasekaran, S.B. Kaginelli, G. Kuppanna, C.G. Kallappa and K.M. Basalingappa, Targeting Notch signalling pathway of cancer stem cells. Stem Cell Investig, 2018. 5: 5.

223. Gouty-Colomer, L.A., et al., Arc expression identifies the lateral amygdala fear memory trace. Mol Psychiatry, 2016. 21(8): 1153.

224. Minatohara, K., M. Akiyoshi and H. Okuno, Role of Immediate-Early Genes in Synaptic Plasticity and Neuronal Ensembles Underlying the Memory Trace. Front Mol Neurosci, 2015. 8: 78.

225. Dobin, A., C.A. Davis, F. Schlesinger, J. Drenkow, C. Zaleski, S. Jha, P. Batut, M. Chaisson and T.R. Gingeras, STAR: ultrafast universal RNA-seq aligner. Bioinformatics, 2013. 29(1): 15–21.

226. Xing, Y., T. Yu, Y.N. Wu, M. Roy, J. Kim and C. Lee, An expectation-maximization algorithm for probabilistic reconstructions of full-length isoforms from splice graphs. Nucleic Acids Res, 2006. 34(10): 3150–60.

227. Ritchie, M.E., B. Phipson, D. Wu, Y. Hu, C.W. Law, W. Shi and G.K. Smyth, limma powers differential expression analyses for RNA-sequencing and microarray studies. Nucleic Acids Res, 2015. 43(7): e47.

228. Benjamini, Y., D. Drai, G. Elmer, N. Kafkafi and I. Golani, Controlling the false discovery rate in behavior genetics research. Behav Brain Res, 2001. 125(1-2): 279–84.

229. Ashburner, M., et al., Gene ontology: tool for the unification of biology. The Gene Ontology Consortium. Nat Genet, 2000. 25(1): 25–9.

230. The Gene Ontology, C., The Gene Ontology Resource: 20 years and still GOing strong. Nucleic Acids Res, 2019. 47(D1): D330–D338.

231. Huang da, W., B.T. Sherman and R.A. Lempicki, Bioinformatics enrichment tools: paths toward the comprehensive functional analysis of large gene lists. Nucleic Acids Res, 2009. 37(1): 1–13.

232. Chang, Y.T., H. Kazui, M. Ikeda, C.W. Huang, S.H. Huang, S.W. Hsu, W.N. Chang and C.C. Chang, Genetic Interaction of APOE and FGF1 is Associated with Memory Impairment and Hippocampal Atrophy in Alzheimer’s Disease. Aging Dis, 2019. 10(3): 510–519.

233. Tao, Q.Q., Y.M. Sun, Z.J. Liu, W. Ni, P. Yang, H.L. Li, S.J. Lu and Z.Y. Wu, A variant within FGF1 is associated with Alzheimer’s disease in the Han Chinese population. Am J Med Genet B Neuropsychiatr Genet, 2014. 165B(2): 131–6.

234. Yamagata, H., et al., Promoter polymorphism in fibroblast growth factor 1 gene increases risk of definite Alzheimer’s disease. Biochem Biophys Res Commun, 2004. 321(2): 320–3.

235. Alazami, A.M., et al., Accelerating novel candidate gene discovery in neurogenetic disorders via whole-exome sequencing of prescreened multiplex consanguineous families. Cell Rep, 2015. 10(2): 148–61.

236. Lyubartseva, G., J.L. Smith, W.R. Markesbery and M.A. Lovell, Alterations of zinc transporter proteins ZnT-1, ZnT-4 and ZnT-6 in preclinical Alzheimer’s disease brain. Brain Pathol, 2010. 20(2): 343–50.

237. Fan, W., Y. Long, Y. Lai, X. Wang, G. Chen and B. Zhu, NPAS4 Facilitates the Autophagic Clearance of Endogenous Tau in Rat Cortical Neurons. J Mol Neurosci, 2016. 58(4): 401–10.

238. Miyashita, A., et al., Genes associated with the progression of neurofibrillary tangles in Alzheimer’s disease. Transl Psychiatry, 2014. 4: e396.

239. Ramamoorthi, K., R. Fropf, G.M. Belfort, H.L. Fitzmaurice, R.M. McKinney, R.L. Neve, T. Otto and Y. Lin, Npas4 regulates a transcriptional program in CA3 required for contextual memory formation. Science, 2011. 334(6063): 1669–75.

240. Ito, S., K. Kimura, M. Haneda, Y. Ishida, M. Sawada and K. Isobe, Induction of matrix metalloproteinases (MMP3, MMP12 and MMP13) expression in the microglia by amyloid-beta stimulation via the PI3K/Akt pathway. Exp Gerontol, 2007. 42(6): 532–7.

241. Zhu, B.L., et al., MMP13 inhibition rescues cognitive decline in Alzheimer transgenic mice via BACE1 regulation. Brain, 2019. 142(1): 176–192.

242. de Souza Silva, M.A., et al., Neurokinin3 receptor as a target to predict and improve learning and memory in the aged organism. Proc Natl Acad Sci U S A, 2013. 110(37): 15097–102.

243. Foroud, T., L.F. Wetherill, J. Kramer, J.A. Tischfield, J.I. Nurnberger, Jr., M.A. Schuckit, X. Xuei and H.J. Edenberg, The tachykinin receptor 3 is associated with alcohol and cocaine dependence. Alcohol Clin Exp Res, 2008. 32(6): 1023–30.

244. Teipel, S.J., M.J. Grothe, K. Wittfeld, W. Hoffmann, K. Hegenscheid, H. Volzke, G. Homuth and H.J. Grabe, Association of a neurokinin 3 receptor polymorphism with the anterior basal forebrain. Neurobiol Aging, 2015. 36(6): 2060–7.

245. Semerdjieva, S., H.H. Abdul-Razak, S.S. Salim, R.J. Yanez-Munoz, P.E. Chen, V. Tarabykin and P. Alifragis, Activation of EphA receptors mediates the recruitment of the adaptor protein Slap, contributing to the downregulation of N-methyl-D-aspartate receptors. Mol Cell Biol, 2013. 33(7): 1442–55.

246. Yaman, E., R. Gasper, C. Koerner, A. Wittinghofer and U.H. Tazebay, RasGEF1A and RasGEF1B are guanine nucleotide exchange factors that discriminate between Rap GTP-binding proteins and mediate Rap2-specific nucleotide exchange. FEBS J, 2009. 276(16): 4607–16.

247. Liu, Y., M. Zhang, W. Hao, I. Mihaljevic, X. Liu, K. Xie, S. Walter and K. Fassbender, Matrix metalloproteinase-12 contributes to neuroinflammation in the aged brain. Neurobiol Aging, 2013. 34(4): 1231–9.

248. Teranishi, Y., et al., Proton myo-inositol cotransporter is a novel gamma-secretase associated protein that regulates Abeta production without affecting Notch cleavage. FEBS J, 2015. 282(17): 3438–51.

249. Wang, J.G., J.A. Strong, W. Xie, R.H. Yang, D.E. Coyle, D.M. Wick, E.D. Dorsey and J.M. Zhang, The chemokine CXCL1/growth related oncogene increases sodium currents and neuronal excitability in small diameter sensory neurons. Mol Pain, 2008. 4: 38.

250. Zhang, K., et al., CXCL1 contributes to beta-amyloid-induced transendothelial migration of monocytes in Alzheimer’s disease. PLoS One, 2013. 8(8): e72744.

251. Zhang, X.F., et al., CXCL1 Triggers Caspase-3 Dependent Tau Cleavage in Long-Term Neuronal Cultures and in the Hippocampus of Aged Mice: Implications in Alzheimer’s Disease. J Alzheimers Dis, 2015. 48(1): 89–104.

252. Davis, W., Jr., The ATP-Binding Cassette Transporter-2 (ABCA2) Overexpression Modulates Sphingosine Levels and Transcription of the Amyloid Precursor Protein (APP) Gene. Curr Alzheimer Res, 2015. 12(9): 847–59.

253. Guedea, A.L., C. Schrick, Y.F. Guzman, K. Leaderbrand, V. Jovasevic, K.A. Corcoran, N.C. Tronson and J. Radulovic, ERK-associated changes of AP-1 proteins during fear extinction. Mol Cell Neurosci, 2011. 47(2): 137–44.

254. Gorbacheva, L., O. Davidova, E. Sokolova, S. Ishiwata, V. Pinelis, S. Strukova and G. Reiser, Endothelial protein C receptor is expressed in rat cortical and hippocampal neurons and is necessary for protective effect of activated protein C at glutamate excitotoxicity. J Neurochem, 2009. 111(4): 967–75.

255. Guo, H., D. Liu, H. Gelbard, T. Cheng, R. Insalaco, J.A. Fernandez, J.H. Griffin and B.V. Zlokovic, Activated protein C prevents neuronal apoptosis via protease activated receptors 1 and 3. Neuron, 2004. 41(4): 563–72.

256. Gahete, M.D., A. Rubio, J. Cordoba-Chacon, F. Gracia-Navarro, R. D. Kineman, J. Avila, R.M. Luque and J.P. Castano, Expression of the ghrelin and neurotensin systems is altered in the temporal lobe of Alzheimer’s disease patients. J Alzheimers Dis, 2010. 22(3): 819–28.

257. Woodworth, H.L., H.M. Batchelor, B.G. Beekly, R. Bugescu, J.A. Brown, G. Kurt, P.M. Fuller and G.M. Leinninger, Neurotensin Receptor-1 Identifies a Subset of Ventral Tegmental Dopamine Neurons that Coordinates Energy Balance. Cell Rep, 2017. 20(8): 1881–1892.

258. Xiao, Z., et al., Activation of neurotensin receptor 1 facilitates neuronal excitability and spatial learning and memory in the entorhinal cortex: beneficial actions in an Alzheimer’s disease model. J Neurosci, 2014. 34(20): 7027–42.

259. D’Haene, E., E.Z. Jacobs, P.J. Volders, T. De Meyer, B. Menten and S. Vergult, Identification of long non-coding RNAs involved in neuronal development and intellectual disability. Sci Rep, 2016. 6: 28396.

260. Katanosaka, K., S. Takatsu, K. Mizumura, K. Naruse and Y. Katanosaka, TRPV2 is required for mechanical nociception and the stretch-evoked response of primary sensory neurons. Sci Rep, 2018. 8(1): 16782.

261. Shibasaki, K., N. Murayama, K. Ono, Y. Ishizaki and M. Tominaga, TRPV2 enhances axon outgrowth through its activation by membrane stretch in developing sensory and motor neurons. J Neurosci, 2010. 30(13): 4601–12.

262. Duits, F.H., et al., Matrix Metalloproteinases in Alzheimer’s Disease and Concurrent Cerebral Microbleeds. J Alzheimers Dis, 2015. 48(3): 711–20.

263. Lakhan, S.E., A. Kirchgessner, D. Tepper and A. Leonard, Matrix metalloproteinases and blood-brain barrier disruption in acute ischemic stroke. Front Neurol, 2013. 4: 32.

264. George, C., G. Gontier, P. Lacube, J.C. Francois, M. Holzenberger and S. Aid, The Alzheimer’s disease transcriptome mimics the neuroprotective signature of IGF-1 receptor-deficient neurons. Brain, 2016. 140(7): 2012–2027.

265. Gontier, G., C. George, Z. Chaker, M. Holzenberger and S. Aid, Blocking IGF Signaling in Adult Neurons Alleviates Alzheimer’s Disease Pathology through Amyloid-beta Clearance. J Neurosci, 2015. 35(33): 11500–13.

266. Nieto Guil, A.F., M. Oksdath, L.A. Weiss, D.J. Grassi, L.J. Sosa, M. Nieto and S. Quiroga, IGF-1 receptor regulates dynamic changes in neuronal polarity during cerebral cortical migration. Sci Rep, 2017. 7(1): 7703.

267. Pristera, A., C. Blomeley, E. Lopes, S. Threlfell, E. Merlini, D. Burdakov, S. Cragg, F. Guillemot and S.L. Ang, Dopamine neuron-derived IGF-1 controls dopamine neuron firing, skill learning, and exploration. Proc Natl Acad Sci U S A, 2019. 116(9): 3817–3826.

268. Moccia, F., E. Zuccolo, T. Soda, F. Tanzi, G. Guerra, L. Mapelli, F. Lodola and E. D’Angelo, Stim and Orai proteins in neuronal Ca(2+) signaling and excitability. Front Cell Neurosci, 2015. 9: 153.

269. Zhang, H., S. Sun, L. Wu, E. Pchitskaya, O. Zakharova, K. Fon Tacer and I. Bezprozvanny, Store-Operated Calcium Channel Complex in Postsynaptic Spines: A New Therapeutic Target for Alzheimer’s Disease Treatment. J Neurosci, 2016. 36(47): 11837–11850.

270. Kim, J., J.P. Moody, C.K. Edgerly, O.L. Bordiuk, K. Cormier, K. Smith, M.F. Beal and R.J. Ferrante, Mitochondrial loss, dysfunction and altered dynamics in Huntington’s disease. Hum Mol Genet, 2010. 19(20): 3919–35.

271. Zhang, L., X.Q. Guo, J.F. Chu, X. Zhang, Z.R. Yan and Y.Z. Li, Potential hippocampal genes and pathways involved in Alzheimer’s disease: a bioinformatic analysis. Genet Mol Res, 2015. 14(2): 7218–32.

272. Wickman, K., C. Karschin, A. Karschin, M.R. Picciotto and D.E. Clapham, Brain localization and behavioral impact of the G-protein-gated K+ channel subunit GIRK4. J Neurosci, 2000. 20(15): 5608–15.

273. Carvill, G.L., et al., Mutations in the GABA Transporter SLC6A1 Cause Epilepsy with Myoclonic-Atonic Seizures. Am J Hum Genet, 2015. 96(5): 808–15.

274. Thoeringer, C.K., et al., The GABA transporter 1 (SLC6A1): a novel candidate gene for anxiety disorders. J Neural Transm (Vienna), 2009. 116(6): 649–57.

275. Fromer, M., et al., De novo mutations in schizophrenia implicate synaptic networks. Nature, 2014. 506(7487): 179–84.

276. Grzmil, P., et al., Targeted disruption of the mouse Npal3 gene leads to deficits in behavior, increased IgE levels, and impaired lung function. Cytogenet Genome Res, 2009. 125(3): 186–200.

277. Huang, R., M. Chen, L. Yang, M. Wagle, S. Guo and B. Hu, MicroRNA-133b Negatively Regulates Zebrafish Single Mauthner-Cell Axon Regeneration through Targeting tppp3 in Vivo. Front Mol Neurosci, 2017. 10: 375.

278. Meyer, M.A., Identification of 17 Highly Expressed Genes within Mouse Lumbar Spinal Cord Anterior Horn Region from an In-Situ Hybridization Atlas of 3430 Genes: Implications for Motor Neuron Disease. Neurol Int, 2014. 6(2): 5367.

279. Chen, J., H. Xiao, Z. Huang, Z. Hu, T. Qi, B. Zhang, X. Tao and S.H. Liu, MicroRNA124 regulate cell growth of prostate cancer cells by targeting iASPP. Int J Clin Exp Pathol, 2014. 7(5): 2283–90.

280. Carneiro, A.M. and R.D. Blakely, Serotonin-, protein kinase C-, and Hic-5-associated redistribution of the platelet serotonin transporter. J Biol Chem, 2006. 281(34): 24769–80.

281. Carneiro, A.M., S.L. Ingram, J.M. Beaulieu, A. Sweeney, S.G. Amara, S.M. Thomas, M.G. Caron and G.E. Torres, The multiple LIM domain-containing adaptor protein Hic-5 synaptically colocalizes and interacts with the dopamine transporter. J Neurosci, 2002. 22(16): 7045–54.

282. Kim, Y., et al., The LIM protein complex establishes a retinal circuitry of visual adaptation by regulating Pax6 alpha-enhancer activity. Elife, 2017. 6.

283. Stern, S., E. Debre, C. Stritt, J. Berger, G. Posern and B. Knoll, A nuclear actin function regulates neuronal motility by serum response factor-dependent gene transcription. J Neurosci, 2009. 29(14): 4512–8.

284. Ueda, K., F. Serajee and A.M. Huq, A Mutation in the ACTA1 gene Manifesting Nemaline Myopathy with Central Nervous System Lesions. J Clin Neurol, 2017. 13(3): 300–302.

285. Chander, P., M.J. Kennedy, B. Winckler and J.P. Weick, Neuron-Specific Gene 2 (NSG2) Encodes an AMPA Receptor Interacting Protein That Modulates Excitatory Neurotransmission. eNeuro, 2019. 6(1).

286. Yap, C.C., L. Digilio, L. McMahon and B. Winckler, The endosomal neuronal proteins Nsg1/NEEP21 and Nsg2/P19 are itinerant, not resident proteins of dendritic endosomes. Sci Rep, 2017. 7(1): 10481.

287. Cheng, C., S.K. Lau and L.C. Doering, Astrocyte-secreted thrombospondin-1 modulates synapse and spine defects in the fragile X mouse model. Mol Brain, 2016. 9(1): 74.

288. Tyzack, G.E., et al., Astrocyte response to motor neuron injury promotes structural synaptic plasticity via STAT3-regulated TSP-1 expression. Nat Commun, 2014. 5: 4294.

289. Bray, E.R., et al., Thrombospondin-1 Mediates Axon Regeneration in Retinal Ganglion Cells. Neuron, 2019. 103(4): 642–657.e7.

290. Ho, A., W. Morishita, D. Atasoy, X. Liu, K. Tabuchi, R.E. Hammer, R.C. Malenka and T.C. Sudhof, Genetic analysis of Mint/X11 proteins: essential presynaptic functions of a neuronal adaptor protein family. J Neurosci, 2006. 26(50): 13089–101.

291. Simms, B.A. and G.W. Zamponi, Neuronal voltage-gated calcium channels: structure, function, and dysfunction. Neuron, 2014. 82(1): 24–45.

292. Sullivan, S.E., G.M. Dillon, J.M. Sullivan and A. Ho, Mint proteins are required for synaptic activity-dependent amyloid precursor protein (APP) trafficking and amyloid beta generation. J Biol Chem, 2014. 289(22): 15374–83.

293. Lin, Y., B.L. Bloodgood, J.L. Hauser, A.D. Lapan, A.C. Koon, T.K. Kim, L.S. Hu, A.N. Malik and M.E. Greenberg, Activity-dependent regulation of inhibitory synapse development by Npas4. Nature, 2008. 455(7217): 1198–204.

294. Spiegel, I., A.R. Mardinly, H.W. Gabel, J.E. Bazinet, C.H. Couch, C. P. Tzeng, D.A. Harmin and M.E. Greenberg, Npas4 regulates excitatory-inhibitory balance within neural circuits through cell-type-specific gene programs. Cell, 2014. 157(5): 1216–29.

295. Ferraro, L., et al., Neurotensin NTS1-dopamine D2 receptor-receptor interactions in putative receptor heteromers: relevance for Parkinson’s disease and schizophrenia. Curr Protein Pept Sci, 2014. 15(7): 681–90.

296. Goehler, H., et al., A protein interaction network links GIT1, an enhancer of huntingtin aggregation, to Huntington’s disease. Mol Cell, 2004. 15(6): 853–65.

297. Awasthi, A., et al., Synaptotagmin-3 drives AMPA receptor endocytosis, depression of synapse strength, and forgetting. Science, 2019. 363(6422).

298. Kitano, J., K. Kimura, Y. Yamazaki, T. Soda, R. Shigemoto, Y. Nakajima and S. Nakanishi, Tamalin, a PDZ domain-containing protein, links a protein complex formation of group 1 metabotropic glutamate receptors and the guanine nucleotide exchange factor cytohesins. J Neurosci, 2002. 22(4): 1280–9.

299. Skillback, T., et al., A novel quantification-driven proteomic strategy identifies an endogenous peptide of pleiotrophin as a new biomarker of Alzheimer’s disease. Sci Rep, 2017. 7(1): 13333.

300. Yamagata, K., et al., Arcadlin is a neural activity-regulated cadherin involved in long term potentiation. J Biol Chem, 1999. 274(27): 19473–1979.

301. Yasuda, S., et al., Activity-induced protocadherin arcadlin regulates dendritic spine number by triggering N-cadherin endocytosis via TAO2beta and p38 MAP kinases. Neuron, 2007. 56(3): 456–71.

302. Menard, C., Y.C. Tse, C. Cavanagh, J.G. Chabot, H. Herzog, C. Schwarzer, T.P. Wong and R. Quirion, Knockdown of prodynorphin gene prevents cognitive decline, reduces anxiety, and rescues loss of group 1 metabotropic glutamate receptor function in aging. J Neurosci, 2013. 33(31): 12792–804.

303. Kayser, M.S., M.J. Nolt and M.B. Dalva, EphB receptors couple dendritic filopodia motility to synapse formation. Neuron, 2008. 59(1): 56–69.

304. Olivares, D., V.K. Deshpande, Y. Shi, D.K. Lahiri, N.H. Greig, J.T. Rogers and X. Huang, N-methyl D-aspartate (NMDA) receptor antagonists and memantine treatment for Alzheimer’s disease, vascular dementia and Parkinson’s disease. Curr Alzheimer Res, 2012. 9(6): 746–58.

305. Baser, M.E., et al., The location of constitutional neurofibromatosis 2 (NF2) splice site mutations is associated with the severity of NF2. J Med Genet, 2005. 42(7): 540–6.

306. Chan, C.C., C.A. Koch, M.I. Kaiser-Kupfer, D.M. Parry, D.H. Gutmann, Z. Zhuang and A.O. Vortmeyer, Loss of heterozygosity for the VF2 gene in retinal and optic nerve lesions of patients with neurofibromatosis 2. J Pathol, 2002. 198(1): 14–20.

307. Bereczki, E., et al., Synaptic markers of cognitive decline in neurodegenerative diseases: a proteomic approach. Brain, 2018. 141(2): 582–595.

308. Li, Y., et al., Lrfn2-Mutant Mice Display Suppressed Synaptic Plasticity and Inhibitory Synapse Development and Abnormal Social Communication and Startle Response. J Neurosci, 2018. 38(26): 5872–5887.

309. Bustos, F.J., et al., Epigenetic editing of the Dlg4/PSD95 gene improves cognition in aged and Alzheimer’s disease mice. Brain, 2017. 140(12): 3252–3268.

310. Li, K., M. Nakajima, I. Ibanez-Tallon and N. Heintz, A Cortical Circuit for Sexually Dimorphic Oxytocin-Dependent Anxiety Behaviors. Cell, 2016. 167(1): 60–72 e11.

311. Corradi, A., et al., SYN2 is an autism predisposing gene: loss-of-function mutations alter synaptic vesicle cycling and axon outgrowth. Hum Mol Genet, 2014. 23(1): 90–103.

312. Binda, A., I. Rivolta, C. Villa, E. Chisci, M. Beghi, C.M. Cornaggia, R. Giovannoni and R. Combi, A Vovel KCNJ2 Mutation Identified in an Autistic Proband Affects the Single Channel Properties of Kir2.1. Front Cell Neurosci, 2018. 12: 76.

313. Martinez-Morillo, E., C. Childs, B.P. Garcia, F.V. Alvarez Menendez, A.D. Romaschin, G. Cervellin, G. Lippi and E.P. Diamandis, Neurofilament medium polypeptide (NFM) protein concentration is increased in CSF and serum samples from patients with brain injury. Clin Chem Lab Med, 2015. 53(10): 1575–84.

314. Anitha, A., et al., Protocadherin alpha (PCDHA) as a novel susceptibility gene for autism. J Psychiatry Neurosci, 2013. 38(3): 192–8.

315. Rubinstein, R., et al., Molecular logic of neuronal self-recognition through protocadherin domain interactions. Cell, 2015. 163(3): 629–42.

316. Maselli, R.A., et al., Presynaptic congenital myasthenic syndrome with a homozygous sequence variant in LAMA5 combines myopia, facial tics, and failure of neuromuscular transmission. Am J Med Genet A, 2017. 173(8): 2240–2245.

317. Dong, W., et al., CAST/ELKS Proteins Control Voltage-Gated Ca(2+) Channel Density and Synaptic Release Probability at a Mammalian Central Synapse. Cell Rep, 2018. 24(2): 284–293 e6.

318. Kaya, N., et al., KCNA4 deficiency leads to a syndrome of abnormal striatum, congenital cataract and intellectual disability. J Med Genet, 2016. 53(11): 786–792.

319. Kang, J.Q. and R.L. Macdonald, Molecular Pathogenic Basis for GABRG2 Mutations Associated With a Spectrum of Epilepsy Syndromes, From Generalized Absence Epilepsy to Dravet Syndrome. JAMA Neurol, 2016. 73(8): 1009–16.

320. Lomash, R.M., X. Gu, R.J. Youle, W. Lu and K.W. Roche, Neurolastin, a Dynamin Family GTPase, Regulates Excitatory Synapses and Spine Density. Cell Rep, 2015. 12(5): 743–51.

321. Choi, S.H., et al., Combined adult neurogenesis and BDNF mimic exercise effects on cognition in an Alzheimer’s mouse model. Science, 2018. 361(6406).

322. Jiao, S.S., et al., Brain-derived neurotrophic factor protects against tau-related neurodegeneration of Alzheimer’s disease. Transl Psychiatry, 2016. 6(10): e907.

323. Bormuth, I., et al., Neuronal basic helix-loop-helix proteins Neurod2/6 regulate cortical commissure formation before midline interactions. J Neurosci, 2013. 33(2): 641–51.

324. Pieper, A., et al., NeuroD2 controls inhibitory circuit formation in the molecular layer of the cerebellum. Sci Rep, 2019. 9(1): 1448.

325. Giza, J., M.J. Urbanski, F. Prestori, B. Bandyopadhyay, A. Yam, V. Friedrich, K. Kelley, E. D’Angelo and M. Goldfarb, Behavioral and cerebellar transmission deficits in mice lacking the autism-linked gene islet brain-2. J Neurosci, 2010. 30(44): 14805–16.

326. Mitz, A.R., T.J. Philyaw, L. Boccuto, A. Shcheglovitov, S.M. Sarasua, W.E. Kaufmann and A. Thurm, Identification of 22q13 genes most likely to contribute to Phelan McDermid syndrome. Eur J Hum Genet, 2018. 26(3): 293–302.

327. Hatanaka, Y., K. Watase, K. Wada and Y. Nagai, Abnormalities in synaptic dynamics during development in a mouse model of spinocerebellar ataxia type 1. Sci Rep, 2015. 5: 16102.

328. Guan, F., T. Zhang, X. Liu, W. Han, H. Lin, L. Li, G. Chen and T. Li, Evaluation of voltage-dependent calcium channel gamma gene families identified several novel potential susceptible genes to schizophrenia. Sci Rep, 2016. 6: 24914.

329. Korber, C., M. Werner, S. Kott, Z.L. Ma and M. Hollmann, The transmembrane AMPA receptor regulatory protein gamma 4 is a more effective modulator of AMPA receptor function than stargazin (gamma 2). J Neurosci, 2007. 27(31): 8442–7.

330. Seigneur, E. and T.C. Sudhof, Genetic Ablation of All Cerebellins Reveals Synapse Organizer Functions in Multiple Regions Throughout the Brain. J Neurosci, 2018. 38(20): 4774–4790.

331. Tao, W., J. Diaz-Alonso, N. Sheng and R.A. Nicoll, Postsynaptic delta1 glutamate receptor assembles and maintains hippocampal synapses via Cbln2 and neurexin. Proc Natl Acad Sci U S A, 2018. 115(23): E5373–E5381.

332. Sadakata, T., et al., Autistic-like phenotypes in Cadps2-knockout mice and aberrant CADPS2 splicing in autistic patients. J Clin Invest, 2007. 117(4): 931–43.

333. Shinoda, Y., T. Sadakata, T. Akagi, Y. Sakamaki, T. Hashikawa, Y. Sano and T. Furuichi, Calcium-dependent activator protein for secretion 2 (CADPS2) deficiency causes abnormal synapse development in hippocampal mossy fiber terminals. Neurosci Lett, 2018. 677: 65–71.

334. Armendariz, B.G., A. Bribian, E. Perez-Martinez, A. Martinez, F. de Castro, E. Soriano and F. Burgaya, Expression of Semaphorin 4F in neurons and brain oligodendrocytes and the regulation of oligodendrocyte precursor migration in the optic nerve. Mol Cell Neurosci, 2012. 49(1): 54–67.

335. Minett, M.S., et al., Endogenous opioids contribute to insensitivity to pain in humans and mice lacking sodium channel Nav1.7. Nat Commun, 2015. 6: 8967.

336. Shiina, N. and M. Tokunaga, RNA granule protein 140 (RNG140), a paralog of RNG105 localized to distinct RNA granules in neuronal dendrites in the adult vertebrate brain. J Biol Chem, 2010. 285(31): 24260–9.

337. Ackerley, S., P.A. James, A. Kalli, S. French, K.E. Davies and K. Talbot, A mutation in the small heat-shock protein HSPB1 leading to distal hereditary motor neuronopathy disrupts neurofilament assembly and the axonal transport of specific cellular cargoes. Hum Mol Genet, 2006. 15(2): 347–54.

338. Baughman, H.E.R., A.F. Clouser, R.E. Klevit and A. Nath, HspB1 and Hsc70 chaperones engage distinct tau species and have different inhibitory effects on amyloid formation. J Biol Chem, 2018. 293(8): 2687–2700.

339. King, M., F. Nafar, J. Clarke and K. Mearow, The small heat shock protein Hsp27 protects cortical neurons against the toxic effects of beta-amyloid peptide. J Neurosci Res, 2009. 87(14): 3161–75.

340. Li, M.D., J.E. Mangold, C. Seneviratne, G.B. Chen, J.Z. Ma, X.Y. Lou and T.J. Payne, Association and interaction analyses of GABBR1 and GABBR2 with nicotine dependence in European- and African-American populations. PLoS One, 2009. 4(9): e7055.

341. Yoo, Y., et al., GABBR2 mutations determine phenotype in rett syndrome and epileptic encephalopathy. Ann Neurol, 2017. 82(3): 466–478.

342. Kakegawa, W., et al., Anterograde C1ql1 signaling is required in order to determine and maintain a single-winner climbing fiber in the mouse cerebellum. Neuron, 2015. 85(2): 316–29.

343. Chew, K.S., D.C. Fernandez, S. Hattar, T.C. Sudhof and D.C. Martinelli, Anatomical and Behavioral Investigation of C1ql3 in the Mouse Suprachiasmatic Nucleus. J Biol Rhythms, 2017. 32(3): 222–236.

344. Martinelli, D.C., K.S. Chew, A. Rohlmann, M.Y. Lum, S. Ressl, S. Hattar, A.T. Brunger, M. Missler and T.C. Sudhof, Expression of C1ql3 in Discrete Neuronal Populations Controls Efferent Synapse Numbers and Diverse Behaviors. Neuron, 2016. 91(5): 1034–1051.

345. Li, Q., B.L. Wang, F.R. Sun, J.Q. Li, X.P. Cao and L. Tan, The role of UNC5C in Alzheimer’s disease. Ann Transl Med, 2018. 6(10): 178.

346. Schroeder, A. and J. de Wit, Leucine-rich repeat-containing synaptic adhesion molecules as organizers of synaptic specificity and diversity. Exp Mol Med, 2018. 50(4): 10.

347. Shao, Q., T. Yang, H. Huang, F. Alarmanazi and G. Liu, Uncoupling of UNC5C with Polymerized TUBB3 in Microtubules Mediates Netrin-1 Repulsion. J Neurosci, 2017. 37(23): 5620–5633.

348. Bertram, L., et al., Family-based association between Alzheimer’s disease and variants in UBQLN1. N Engl J Med, 2005. 352(9): 884–94.

349. Rehker, J., et al., Caspase-8, association with Alzheimer’s Disease and functional analysis of rare variants. PLoS One, 2017. 12(10): e0185777.

350. Saha, P. and S.C. Biswas, Amyloid-beta induced astrocytosis and astrocyte death: Implication of FoxO3a-Bim-caspase3 death signaling. Mol Cell Neurosci, 2015. 68: 203–11.

351. Sanphui, P. and S.C. Biswas, FoxO3a is activated and executes neuron death via Bim in response to beta-amyloid. Cell Death Dis, 2013. 4: e625.

352. Woods, Y.L., P. Cohen, W. Becker, R. Jakes, M. Goedert, X. Wang and C.G. Proud, The kinase DYRK phosphorylates protein-synthesis initiation factor eIF2Bepsilon at Ser539 and the microtubule-associated protein tau at Thr212: potential role for DYRK as a glycogen synthase kinase 3-priming kinase. Biochem J, 2001. 355(Pt 3): 609–15.

353. Inoue, M., et al., Human brain proteins showing neuron-specific interactions with gamma-secretase. FEBS J, 2015. 282(14): 2587–99.

354. Kolsch, H., et al., Gene polymorphisms in prodynorphin (PDYN) are associated with episodic memory in the elderly. J Neural Transm (Vienna), 2009. 116(7): 897–903.

355. Menard, C., H. Herzog, C. Schwarzer and R. Quirion, Possible role of dynorphins in Alzheimer’s disease and age-related cognitive deficits. Neurodegener Dis, 2014. 13(2-3): 82–5.

356. Yakovleva, T., Z. Marinova, A. Kuzmin, N.G. Seidah, V. Haroutunian, L. Terenius and G. Bakalkin, Dysregulation of dynorphins in Alzheimer disease. Neurobiol Aging, 2007. 28(11): 1700–8.

357. Abdullah, L., G. Ait-Ghezala, F. Crawford, T.A. Crowell, W.W. Barker, R. Duara and M. Mullan, The cyclooxygenase 2-765 C promoter allele is a protective factor for Alzheimer’s disease. Neurosci Lett, 2006. 395(3): 240–3.

358. Chen, Q., B. Liang, Z. Wang, X. Cheng, Y. Huang, Y. Liu and Z. Huang, Influence of four polymorphisms in ABCA1 and PTGS2 genes on risk of Alzheimer’s disease: a meta-analysis. Neurol Sci, 2016. 37(8): 1209–20.

359. Guan, P.P. and P. Wang, Integrated communications between cyclooxygenase-2 and Alzheimer’s disease. FASEB J, 2019. 33(1): 13–33.

360. Vito, P., E. Lacana and L. D’Adamio, Interfering with apoptosis: Ca(2+)-binding protein ALG-2 and Alzheimer’s disease gene ALG-3. Science, 1996. 271(5248): 521–5.

361. Nelson, C.D. and M. Sheng, Gpr3 stimulates Abeta production via interactions with APP and beta-arrestin2. PLoS One, 2013. 8(9): e74680.

362. Thathiah, A., et al., The orphan G protein-coupled receptor 3 modulates amyloid-beta peptide generation in neurons. Science, 2009. 323(5916): 946–51.

363. Huang, Y., et al., Loss of GPR3 reduces the amyloid plaque burden and improves memory in Alzheimer’s disease mouse models. Science Translational Medicine, 2015. 7(309): 309ra164-309ra164.

364. Campolongo, P., et al., Systemic administration of substance P recovers beta amyloid-induced cognitive deficits in rat: involvement of Kv potassium channels. PLoS One, 2013. 8(11): e78036.

365. Pieri, M., et al., SP protects cerebellar granule cells against beta-amyloid-induced apoptosis by down-regulation and reduced activity of Kv4 potassium channels. Neuropharmacology, 2010. 58(1): 268–76.

366. Beyer, K., J.I. Lao, M. Gomez, N. Riutort, P. Latorre, J.L. Mate and A. Ariza, Alzheimer’s disease and the cystatin C gene polymorphism: an association study. Neurosci Lett, 2001. 315(1-2): 17–20.

367. Finckh, U., et al., Genetic association of a cystatin C gene polymorphism with late-onset Alzheimer disease. Arch Neurol, 2000. 57(11): 1579–83.

368. Kaur, G. and E. Levy, Cystatin C in Alzheimer’s disease. Front Mol Neurosci, 2012. 5: 79.

369. Hettiarachchi, N., M. Dallas, M. Al-Owais, H. Griffiths, N. Hooper, J. Scragg, J. Boyle and C. Peers, Heme oxygenase-1 protects against Alzheimer’s amyloid-beta(1-42)-induced toxicity via carbon monoxide production. Cell Death Dis, 2014. 5: e1569.

370. Hettiarachchi, N.T., J.P. Boyle, M.L. Dallas, M.M. Al-Owais, J.L. Scragg and C. Peers, Heme oxygenase-1 derived carbon monoxide suppresses Abeta1-42 toxicity in astrocytes. Cell Death Dis, 2017. 8(6): e2884.

371. Allen, M., et al., Glutathione S-transferase omega genes in Alzheimer and Parkinson disease risk, age-at-diagnosis and brain gene expression: an association study with mechanistic implications. Mol Neurodegener, 2012. 7: 13.

372. Li, Y.J., et al., Glutathione S-transferase omega-1 modifies age-at-onset of Alzheimer disease and Parkinson disease. Hum Mol Genet, 2003. 12(24): 3259–67.

373. Moon, M., E.S. Jung, S.G. Jeon, M.Y. Cha, Y. Jang, W. Kim, C. Lopes, I. Mook-Jung and K.S. Kim, Nurr1 (NR4A2) regulates Alzheimer’s disease-related pathogenesis and cognitive function in the 5XFAD mouse model. Aging Cell, 2019. 18(1): e12866.

374. Meilandt, W.J., G.Q. Yu, J. Chin, E.D. Roberson, J.J. Palop, T. Wu, K. Scearce-Levie and L. Mucke, Enkephalin elevations contribute to neuronal and behavioral impairments in a transgenic mouse model of Alzheimer’s disease. J Neurosci, 2008. 28(19): 5007–17.

375. Guo, Q., W. Fu, J. Xie, H. Luo, S.F. Sells, J.W. Geddes, V. Bondada, V.M. Rangnekar and M.P. Mattson, Par-4 is a mediator of neuronal degeneration associated with the pathogenesis of Alzheimer disease. Nat Med, 1998. 4(8): 957–62.

376. Xie, J. and Q. Guo, PAR-4 is involved in regulation of beta-secretase cleavage of the Alzheimer amyloid precursor protein. J Biol Chem, 2005. 280(14): 13824–32.

377. Ishiki, A., M. Kamada, Y. Kawamura, C. Terao, F. Shimoda, N. Tomita, H. Arai and K. Furukawa, Glial fibrillar acidic protein in the cerebrospinal fluid of Alzheimer’s disease, dementia with Lewy bodies, and frontotemporal lobar degeneration. J Neurochem, 2016. 136(2): 258–61.

378. Oeckl, P., et al., Glial Fibrillary Acidic Protein in Serum is Increased in Alzheimer’s Disease and Correlates with Cognitive Impairment. J Alzheimers Dis, 2019. 67(2): 481–488.

379. Hoe, H.S., et al., The metalloprotease inhibitor TIMP-3 regulates amyloid precursor protein and apolipoprotein E receptor proteolysis. J Neurosci, 2007. 27(40): 10895–905.

380. Buxbaum, J.D., E.K. Choi, Y. Luo, C. Lilliehook, A.C. Crowley, D. E. Merriam and W. Wasco, Calsenilin: a calcium-binding protein that interacts with the presenilins and regulates the levels of a presenilin fragment. Nat Med, 1998. 4(10): 1177–81.

381. Crary, J.F., C.Y. Shao, S.S. Mirra, A.I. Hernandez and T.C. Sacktor, Atypical protein kinase C in neurodegenerative disease I: PKMzeta aggregates with limbic neurofibrillary tangles and AMPA receptors in Alzheimer disease. J Neuropathol Exp Neurol, 2006. 65(4): 319–26.

382. Liang, D., G. Han, X. Feng, J. Sun, Y. Duan and H. Lei, Concerted perturbation observed in a hub network in Alzheimer’s disease. PLoS One, 2012. 7(7): e40498.

383. Park, B., W. Lee and K. Han, Modeling the interactions of Alzheimer-related genes from the whole brain microarray data and diffusion tensor images of human brain. BMC Bioinformatics, 2012. 13 Suppl 7: S10.

384. Desrumaux, C., et al., Increased amyloid-beta peptide-induced memory deficits in phospholipid transfer protein (PLTP) gene knockout mice. Neuropsychopharmacology, 2013. 38(5): 817–25.

385. Dong, W., J.J. Albers and S. Vuletic, Phospholipid transfer protein reduces phosphorylation of tau in human neuronal cells. J Neurosci Res, 2009. 87(14): 3176–85.

386. Mansuy, M., et al., Deletion of plasma Phospholipid Transfer Protein (PLTP) increases microglial phagocytosis and reduces cerebral amyloid-beta deposition in the J20 mouse model of Alzheimer’s disease. Oncotarget, 2018. 9(28): 19688–19703.

387. Tong, Y., et al., Phospholipid transfer protein (PLTP) deficiency accelerates memory dysfunction through altering amyloid precursor protein (APP) processing in a mouse model of Alzheimer’s disease. Hum Mol Genet, 2015. 24(19): 5388–403.

388. Vuletic, S., E.R. Peskind, S.M. Marcovina, J.F. Quinn, M.C. Cheung, H. Kennedy, J.A. Kaye, L.W. Jin and J.J. Albers, Reduced CSF PLTP activity in Alzheimer’s disease and other neurologic diseases; PLTP induces ApoE secretion in primary human astrocytes in vitro. J Neurosci Res, 2005. 80(3): 406–13.

389. Wang, H., Y. Yu, W. Chen, Y. Cui, T. Luo, J. Ma, X.C. Jiang and S. Qin, PLTP deficiency impairs learning and memory capabilities partially due to alteration of amyloid-beta metabolism in old mice. J Alzheimers Dis, 2014. 39(1): 79–88.

390. Hernandez, F., J.J. Lucas and J. Avila, GSK3 and tau: two convergence points in Alzheimer’s disease. J Alzheimers Dis, 2013. 33 Suppl 1: S141–4.

391. Llorens-Martin, M., J. Jurado, F. Hernandez and J. Avila, GSK-3beta, a pivotal kinase in Alzheimer disease. Front Mol Neurosci, 2014. 7: 46.

392. Guedes, J.R., T. Lao, A.L. Cardoso and J. El Khoury, Roles of Microglial and Monocyte Chemokines and Their Receptors in Regulating Alzheimer’s Disease-Associated Amyloid-beta and Tau Pathologies. Front Neurol, 2018. 9: 549.

393. Naert, G. and S. Rivest, A deficiency in CCR2+ monocytes: the hidden side of Alzheimer’s disease. J Mol Cell Biol, 2013. 5(5): 284–93.

394. Westin, K., P. Buchhave, H. Nielsen, L. Minthon, S. Janciauskiene and O. Hansson, CCL2 is associated with a faster rate of cognitive decline during early stages of Alzheimer’s disease. PLoS One, 2012. 7(1): e30525.

395. Khandelwal, P.J., S.B. Dumanis, A.M. Herman, G.W. Rebeck and C.E. Moussa, Wild type and P301L mutant Tau promote neuro-inflammation and alpha-Synuclein accumulation in lentiviral gene delivery models. Mol Cell Neurosci, 2012. 49(1): 44–53.

396. Lee, Y., J.S. Lee, K.J. Lee, R.S. Turner, H.S. Hoe and D.T.S. Pak, Polo-like kinase 2 phosphorylation of amyloid precursor protein regulates activity-dependent amyloidogenic processing. Neuropharmacology, 2017. 117: 387–400.

397. Seeburg, D.P., M. Feliu-Mojer, J. Gaiottino, D.T. Pak and M. Sheng, Critical role of CDK5 and Polo-like kinase 2 in homeostatic synaptic plasticity during elevated activity. Neuron, 2008. 58(4): 571–83.

398. Kern, S., et al., Association of Cerebrospinal Fluid Neurofilament Light Protein With Risk of Mild Cognitive Impairment Among Individuals Without Cognitive Impairment. JAMA Neurol, 2018.

399. Lewczuk, P., et al., Plasma neurofilament light as a potential biomarker of neurodegeneration in Alzheimer’s disease. Alzheimers Res Ther, 2018. 10(1): 71.

400. Preische, O., et al., Serum neurofilament dynamics predicts neurodegeneration and clinical progression in presymptomatic Alzheimer’s disease. Nat Med, 2019. 25(2): 277–283.

401. Mattsson, N., N.C. Cullen, U. Andreasson, H. Zetterberg and K. Blennow, Association Between Longitudinal Plasma Neurofilament Light and Neurodegeneration in Patients With Alzheimer Disease. JAMA Neurology, 2019. 76(7): 791–799.

402. Corcoran, N.M., D. Martin, B. Hutter-Paier, M. Windisch, T. Nguyen, L. Nheu, L.E. Sundstrom, A.J. Costello and C.M. Hovens, Sodium selenate specifically activates PP2A phosphatase, dephosphorylates tau and reverses memory deficits in an Alzheimer’s disease model. J Clin Neurosci, 2010. 17(8): 1025–33.

403. Du, X., H. Li, Z. Wang, S. Qiu, Q. Liu and J. Ni, Selenoprotein P and selenoprotein M block Zn2+-mediated Abeta42 aggregation and toxicity. Metallomics, 2013. 5(7): 861–70.

404. Song, G., Z. Zhang, L. Wen, C. Chen, Q. Shi, Y. Zhang, J. Ni and Q. Liu, Selenomethionine ameliorates cognitive decline, reduces tau hyperphosphorylation, and reverses synaptic deficit in the triple transgenic mouse model of Alzheimer’s disease. J Alzheimers Dis, 2014. 41(1): 85–99.

405. van Eersel, J., Y.D. Ke, X. Liu, F. Delerue, J.J. Kril, J. Gotz and L.M. Ittner, Sodium selenate mitigates tau pathology, neurodegeneration, and functional deficits in Alzheimer’s disease models. Proc Natl Acad Sci U S A, 2010. 107(31): 13888–93.

406. Liu, C.C., J. Hu, N. Zhao, J. Wang, N. Wang, J.R. Cirrito, T. Kanekiyo, D.M. Holtzman and G. Bu, Astrocytic LRP1 Mediates Brain Abeta Clearance and Impacts Amyloid Deposition. J Neurosci, 2017. 37(15): 4023–4031.

407. Storck, S.E., A.M.S. Hartz, J. Bernard, A. Wolf, A. Kachlmeier, A. Mahringer, S. Weggen, J. Pahnke and C.U. Pietrzik, The concerted amyloid-beta clearance of LRP1 and ABCB1/P-gp across the blood-brain barrier is linked by PICALM. Brain Behav Immun, 2018. 73: 21–33.

408. Storck, S.E., et al., Endothelial LRP1 transports amyloid-beta(1-42) across the blood-brain barrier. J Clin Invest, 2016. 126(1): 123–36.

409. Kook, S.Y., et al., Crucial role of calbindin-D28k in the pathogenesis of Alzheimer’s disease mouse model. Cell Death Differ, 2014. 21(10): 1575–87.

410. Riascos, D., D. de Leon, A. Baker-Nigh, A. Nicholas, R. Yukhananov, J. Bu, C.K. Wu and C. Geula, Age-related loss of calcium buffering and selective neuronal vulnerability in Alzheimer’s disease. Acta Neuropathol, 2011. 122(5): 565–76.

411. Park, D., M. Na, J.A. Kim, U. Lee, E. Cho, M. Jang and S. Chang, Activation of CaMKIV by soluble amyloid-beta1-42 impedes trafficking of axonal vesicles and impairs activity-dependent synaptogenesis. Sci Signal, 2017. 10(487).

412. Kaden, D., P. Voigt, L.M. Munter, K.D. Bobowski, M. Schaefer and G. Multhaup, Subcellular localization and dimerization of APLP1 are strikingly different from APP and APLP2. J Cell Sci, 2009. 122(Pt 3): 368–77.

413. D’Addario, C., et al., Epigenetic regulation of fatty acid amide hydrolase in Alzheimer disease. PLoS One, 2012. 7(6): e39186.

414. Montanari, S., et al., Fatty Acid Amide Hydrolase (FAAH), Acetylcholinesterase (AChE), and Butyrylcholinesterase (BuChE): Networked Targets for the Development of Carbamates as Potential Anti-Alzheimer’s Disease Agents. J Med Chem, 2016. 59(13): 6387–406.

415. Mulder, J., et al., Molecular reorganization of endocannabinoid signalling in Alzheimer’s disease. Brain, 2011. 134(Pt 4): 1041–60.

416. Williamson, R., et al., CRMP2 hyperphosphorylation is characteristic of Alzheimer’s disease and not a feature common to other neurodegenerative diseases. J Alzheimers Dis, 2011. 27(3): 615–25.

417. Xing, H., Y.A. Lim, J.R. Chong, J.H. Lee, D. Aarsland, C.G. Ballard, P.T. Francis, C.P. Chen and M.K. Lai, Increased phosphorylation of collapsin response mediator protein-2 at Thr514 correlates with beta-amyloid burden and synaptic deficits in Lewy body dementias. Mol Brain, 2016. 9(1): 84.

418. Abdul-Hay, S.O., T. Sahara, M. McBride, D. Kang and M.A. Leissring, Identification of BACE2 as an avid ss-amyloid-degrading protease. Mol Neurodegener, 2012. 7: 46.

419. Montibeller, L. and J. de Belleroche, Amyotrophic lateral sclerosis (ALS) and Alzheimer’s disease (AD) are characterised by differential activation of ER stress pathways: focus on UPR target genes. Cell Stress Chaperones, 2018. 23(5): 897–912.

420. Salminen, A., A. Kauppinen, T. Suuronen, K. Kaarniranta and J. Ojala, ER stress in Alzheimer’s disease: a novel neuronal trigger for inflammation and Alzheimer’s pathology. J Neuroinflammation, 2009. 6: 41.

421. Liu, C.C., C.C. Liu, T. Kanekiyo, H. Xu and G. Bu, Apolipoprotein E and Alzheimer disease: risk, mechanisms and therapy. Nat Rev Neurol, 2013. 9(2): 106–18.

422. Miller, C.C., D.M. McLoughlin, K.F. Lau, M.E. Tennant and B. Rogelj, The X11 proteins, Abeta production and Alzheimer’s disease. Trends Neurosci, 2006. 29(5): 280–5.

423. Kim, J., et al., BRI2 (ITM2b) inhibits Abeta deposition in vivo. J Neurosci, 2008. 28(23): 6030–6.

424. Delmas, E., N. Jah, C. Pirou, S. Bouleau, N. Le Floch, J.L. Vayssiere, B. Mignotte and F. Renaud, FGF1 C-terminal domain and phosphorylation regulate intracrine FGF1 signaling for its neurotrophic and anti-apoptotic activities. Cell Death Dis, 2016. 7: e2079.

425. Coutellier, L., S. Beraki, P.M. Ardestani, N.L. Saw and M. Shamloo, Npas4: a neuronal transcription factor with a key role in social and cognitive functions relevant to developmental disorders. PLoS One, 2012. 7(9): e46604.

426. Hunt, D., G. Raivich and P.N. Anderson, Activating transcription factor 3 and the nervous system. Front Mol Neurosci, 2012. 5: 7.

427. Warsito, D., S. Sjostrom, S. Andersson, O. Larsson and B. Sehat, Nuclear IGF1R is a transcriptional co-activator of LEF1/TCF. EMBO Rep, 2012. 13(3): 244–50.

428. Christopherson, K.S., et al., Thrombospondins are astrocyte-secreted proteins that promote CNS synaptogenesis. Cell, 2005. 120(3): 421–33.

429. Jing, G., C. Westwell-Roper, J. Chen, G. Xu, C.B. Verchere and A. Shalev, Thioredoxin-interacting protein promotes islet amyloid polypeptide expression through miR-124a and FoxA2. J Biol Chem, 2014. 289(17): 11807–15.

430. Zhang, S.J., et al., A signaling cascade of nuclear calcium-CREB-ATF3 activated by synaptic NMDA receptors defines a gene repression module that protects against extrasynaptic NMDA receptor-induced neuronal cell death and ischemic brain damage. J Neurosci, 2011. 31(13): 4978–90.

431. Weng, J., et al., Deletion of G protein-coupled receptor 48 leads to ocular anterior segment dysgenesis (ASD) through down-regulation of Pitx2. Proc Natl Acad Sci U S A, 2008. 105(16): 6081–6.

432. Zhang, S., J. Li, R. Lea, K. Vleminckx and E. Amaya, Fezf2 promotes neuronal differentiation through localised activation of Wnt/beta-catenin signalling during forebrain development. Development, 2014. 141(24): 4794–805.

433. Zuccotti, A., C. Le Magueresse, M. Chen, A. Neitz and H. Monyer, The transcription factor Fezf2 directs the differentiation of neural stem cells in the subventricular zone toward a cortical phenotype. Proc Natl Acad Sci U S A, 2014. 111(29): 10726–31.

434. Schlingensiepen, K.H., F. Wollnik, M. Kunst, R. Schlingensiepen, T. Herdegen and W. Brysch, The role of Jun transcription factor expression and phosphorylation in neuronal differentiation, neuronal cell death, and plastic adaptations in vivo. Cell Mol Neurobiol, 1994. 14(5): 487–505.

435. Li, R., R. Strohmeyer, Z. Liang, L.F. Lue and J. Rogers, CCAAT/enhancer binding protein delta (C/EBPdelta) expression and elevation in Alzheimer’s disease. Neurobiol Aging, 2004. 25(8): 991–9.

436. Wang, S.M., S.W. Lim, Y.H. Wang, H.Y. Lin, M.D. Lai, C.Y. Ko and J.M. Wang, Astrocytic CCAAT/Enhancer-binding protein delta contributes to reactive oxygen species formation in neuroinflammation. Redox Biol, 2018. 16: 104–112.

437. Li, Z., et al., PIM1 kinase phosphorylates the human transcription factor FOXP3 at serine 422 to negatively regulate its activity under inflammation. J Biol Chem, 2014. 289(39): 26872–81.

438. Rainio, E.M., J. Sandholm and P.J. Koskinen, Cutting edge: Transcriptional activity of NFATc1 is enhanced by the Pim-1 kinase. J Immunol, 2002. 168(4): 1524–7.

439. Velazquez, R., D.M. Shaw, A. Caccamo and S. Oddo, Pim1 inhibition as a novel therapeutic strategy for Alzheimer’s disease. Mol Neurodegener, 2016. 11(1): 52.

440. Zippo, A., A. De Robertis, R. Serafini and S. Oliviero, PIM1-dependent phosphorylation of histone H3 at serine 10 is required for MYC-dependent transcriptional activation and oncogenic transformation. Nat Cell Biol, 2007. 9(8): 932–44.

441. Lee, A.K. and P.R. Potts, A Comprehensive Guide to the MAGE Family of Ubiquitin Ligases. J Mol Biol, 2017. 429(8): 1114–1142.

442. Coon, K.D., et al., Preliminary demonstration of an allelic association of the IREB2 gene with Alzheimer’s disease. J Alzheimers Dis, 2006. 9(3): 225–33.

443. Takahashi-Makise, N., D.M. Ward and J. Kaplan, On the mechanism of iron sensing by IRP2: new players, new paradigms. Nat Chem Biol, 2009.5(12): 874–875.

444. von Bernhardi, R., F. Cornejo, G.E. Parada and J. Eugenin, Role of TGFbeta signaling in the pathogenesis of Alzheimer’s disease. Front Cell Neurosci, 2015. 9: 426.

445. Suenaga, K., et al., Muscleblind-like 1 knockout mice reveal novel splicing defects in the myotonic dystrophy brain. PLoS One, 2012. 7(3): e33218.

446. Bluthgen, N., M. van Bentum, B. Merz, D. Kuhl and G. Hermey, Profiling the MAPK/ERK dependent and independent activity regulated transcriptional programs in the murine hippocampus in vivo. Sci Rep, 2017. 7: 45101.

447. Namwanje, M. and C.W. Brown, Activins and Inhibins: Roles in Development, Physiology, and Disease. Cold Spring Harbor Perspectives in Biology, 2016. 8(7): a021881.

448. Li, X., W. Wei, Q.Y. Zhao, J. Widagdo, D. Baker-Andresen, C.R. Flavell, A. D’Alessio, Y. Zhang and T.W. Bredy, Neocortical Tet3-mediated accumulation of 5-hydroxymethylcytosine promotes rapid behavioral adaptation. Proc Natl Acad Sci U S A, 2014. 111(19): 7120–5.

449. Perera, A., et al., TET3 is recruited by REST for context-specific hydroxymethylation and induction of gene expression. Cell Rep, 2015. 11(2): 283–94.

450. Farley, J.E., T.C. Burdett, R. Barria, L.J. Neukomm, K.P. Kenna, J.E. Landers and M.R. Freeman, Transcription factor Pebbled/RREB1 regulates injury-induced axon degeneration. Proc Natl Acad Sci U S A, 2018. 115(6): 1358–1363.

451. Cates, H.M., et al., Threonine 149 phosphorylation enhances DeltaFosB transcriptional activity to control psychomotor responses to cocaine. J Neurosci, 2014. 34(34): 11461–9.

452. Eagle, A.L., P.A. Gajewski, M. Yang, M.E. Kechner, B.S. Al Masraf, P.J. Kennedy, H. Wang, M.S. Mazei-Robison and A.J. Robison, Experience-Dependent Induction of Hippocampal DeltaFosB Controls Learning. J Neurosci, 2015. 35(40): 13773–83.

453. Grassi, D., et al., Neuronal Activity, TGFbeta-Signaling and Unpredictable Chronic Stress Modulate Transcription of Gadd45 Family Members and DNA Methylation in the Hippocampus. Cereb Cortex, 2017. 27(8): 4166–4181.

454. Tamai, S., K. Imaizumi, N. Kurabayashi, M.D. Nguyen, T. Abe, M. Inoue, Y. Fukada and K. Sanada, Neuroprotective role of the basic leucine zipper transcription factor NFIL3 in models of amyotrophic lateral sclerosis. J Biol Chem, 2014. 289(3): 1629–38.

455. Chen, X., K. Cho, B.H. Singer and H. Zhang, The nuclear transcription factor PKNOX2 is a candidate gene for substance dependence in European-origin women. PLoS One, 2011. 6(1): e16002.

456. Wang, K.S., Q. Zhang, X. Liu, L. Wu and M. Zeng, PKNOX2 is associated with formal thought disorder in schizophrenia: a meta-analysis of two genome-wide association studies. J Mol Neurosci, 2012. 48(1): 265–72.

457. Feng, J., M.A. Lawson and P. Melamed, A proteomic comparison of immature and mature mouse gonadotrophs reveals novel differentially expressed nuclear proteins that regulate gonadotropin gene transcription and RNA splicing. Biol Reprod, 2008. 79(3): 546–61.

458. Lang, B., T.M. Alrahbeni, D.S. Clair, D.H. Blackwood, C. International Schizophrenia, C.D. McCaig and S. Shen, HDAC9 is implicated in schizophrenia and expressed specifically in post-mitotic neurons but not in adult neural stem cells. Am J Stem Cells, 2012. 1(1): 31–41.

459. Lund, I.V., Y. Hu, Y.H. Raol, R.S. Benham, R. Faris, S.J. Russek and A.R. Brooks-Kayal, BDNF selectively regulates GABAA receptor transcription by activation of the JAK/STAT pathway. Sci Signal, 2008. 1(41): ra9.

460. Bayam, E., G.S. Sahin, G. Guzelsoy, G. Guner, A. Kabakcioglu and G. Ince-Dunn, Genome-wide target analysis of NEUROD2 provides new insights into regulation of cortical projection neuron migration and differentiation. BMC Genomics, 2015. 16: 681.

461. Chen, F., J.T. Moran, Y. Zhang, K.M. Ates, D. Yu, L.A. Schrader, P.M. Das, F.E. Jones and B.J. Hall, The transcription factor NeuroD2 coordinates synaptic innervation and cell intrinsic properties to control excitability of cortical pyramidal neurons. J Physiol, 2016. 594(13): 3729–44.

462. Lin, C.H., S. Hansen, Z. Wang, D.R. Storm, S.J. Tapscott and J.M. Olson, The dosage of the neuroD2 transcription factor regulates amygdala development and emotional learning. Proc Natl Acad Sci U S A, 2005. 102(41): 14877–82.

463. Agundez, J.A., et al., Heme Oxygenase-1 and 2 Common Genetic Variants and Risk for Multiple Sclerosis. Sci Rep, 2016. 6: 20830.

464. Ayuso, P., C. Martinez, P. Pastor, O. Lorenzo-Betancor, A. Luengo, F.J. Jimenez-Jimenez, H. Alonso-Navarro, J.A. Agundez and E. Garcia-Martin, An association study between Heme oxygenase-1 genetic variants and Parkinson’s disease. Front Cell Neurosci, 2014. 8: 298.

465. Lin, Q., et al., Heme oxygenase-1 protein localizes to the nucleus and activates transcription factors important in oxidative stress. J Biol Chem, 2007. 282(28): 20621–33.

466. Schipper, H.M. and W. Song, A heme oxygenase-1 transducer model of degenerative and developmental brain disorders. Int J Mol Sci, 2015. 16(3): 5400–19.

467. Crespo-Barreto, J., J.D. Fryer, C.A. Shaw, H.T. Orr and H.Y. Zoghbi, Partial loss of ataxin-1 function contributes to transcriptional dysregulation in spinocerebellar ataxia type 1 pathogenesis. PLoS Genet, 2010. 6(7): e1001021.

468. Lu, H.C., et al., Disruption of the ATXN1-CIC complex causes a spectrum of neurobehavioral phenotypes in mice and humans. Nat Genet, 2017. 49(4): 527–536.

469. Mitchell, A.C., et al., MEF2C transcription factor is associated with the genetic and epigenetic risk architecture of schizophrenia and improves cognition in mice. Mol Psychiatry, 2018. 23(1): 123–132.

470. Herrera, F.J., T. Yamaguchi, H. Roelink and R. Tjian, Core promoter factor TAF9B regulates neuronal gene expression. Elife, 2014. 3: e02559.

471. Auderset, L., L.M. Landowski, L. Foa and K.M. Young, Low Density Lipoprotein Receptor Related Proteins as Regulators of Neural Stem and Progenitor Cell Function. Stem Cells Int, 2016. 2016: 2108495.

472. Shah, M., O.Y. Baterina, Jr., V. Taupin and M.G. Farquhar, ARH directs megalin to the endocytic recycling compartment to regulate its proteolysis and gene expression. J Cell Biol, 2013. 202(1): 113–27.

473. Spuch, C., S. Ortolano and C. Navarro, LRP-1 and LRP-2 receptors function in the membrane neuron. Trafficking mechanisms and proteolytic processing in Alzheimer’s disease. Front Physiol, 2012. 3: 269.

474. Santra, M., M. Chopp, Z.G. Zhang, M. Lu, S. Santra, A. Nalani, S. Santra and D.C. Morris, Thymosin beta 4 mediates oligodendrocyte differentiation by upregulating p38 MAPK. Glia, 2012. 60(12): 1826–38.

475. Carrillo-Garcia, C., S. Prochnow, I.K. Simeonova, J. Strelau, G. Holzl-Wenig, C. Mandl, K. Unsicker, O. von Bohlen Und Halbach and F. Ciccolini, Growth/differentiation factor 15 promotes EGFR signalling,and regulates proliferation and migration in the hippocampus of neonatal and young adult mice. Development, 2014. 141(4): 773–83.

476. Katoh, K., Y. Omori, A. Onishi, S. Sato, M. Kondo and T. Furukawa, Blimp1 suppresses Chx10 expression in differentiating retinal photoreceptor precursors to ensure proper photoreceptor development. J Neurosci, 2010. 30(19): 6515–26.

477. Kinameri, E., T. Inoue, J. Aruga, I. Imayoshi, R. Kageyama, T. Shimogori and A.W. Moore, Prdm proto-oncogene transcription factor family expression and interaction with the Notch-Hes pathway in mouse neurogenesis. PLoS One, 2008. 3(12): e3859.

478. Yu, M., et al., Suppression of MAPK11 or HIPK3 reduces mutant Huntingtin levels in Huntington’s disease models. Cell Res, 2017. 27(12): 1441–1465.

479. Knoepfler, P.S., P.F. Cheng and R.N. Eisenman, N-myc is essential during neurogenesis for the rapid expansion of progenitor cell populations and the inhibition of neuronal differentiation. Genes Dev, 2002. 16(20): 2699–712.

480. Zinin, N., I. Adameyko, M. Wilhelm, N. Fritz, P. Uhlen, P. Ernfors and M.A. Henriksson, MYC proteins promote neuronal differentiation by controlling the mode of progenitor cell division. EMBO Rep, 2014. 15(4): 383–91.

481. Ancin, I., J.A. Cabranes, B. Vazquez-Alvarez, J.L. Santos, E. Sanchez-Morla, M. Alaerts, J. Del-Favero and A. Barabash, NR4A2: effects of an “orphan” receptor on sustained attention in a schizophrenic population. Schizophr Bull, 2013. 39(3): 555–63.

482. Buervenich, S., et al., NURR1 mutations in cases of schizophrenia and manic-depressive disorder. Am J Med Genet, 2000. 96(6): 808–13.

483. Jacobs, F.M., A.J. van der Linden, Y. Wang, L. von Oerthel, H.S. Sul, J.P. Burbach and M.P. Smidt, Identification of Dlk1, Ptpru and Klhl1 as novel Nurr1 target genes in meso-diencephalic dopamine neurons. Development, 2009. 136(14): 2363–73.

484. Harris, A., G. Masgutova, A. Collin, M. Toch, M. Hidalgo-Figueroa, B. Jacob, L.M. Corcoran, C. Francius and F. Clotman, Onecut Factors and Pou2f2 Regulate the Distribution of V2 Interneurons in the Mouse Developing Spinal Cord. Front Cell Neurosci, 2019. 13: 184.

485. Lillycrop, K.A. and D.S. Latchman, Alternative splicing of the Oct-2 transcription factor RNA is differentially regulated in neuronal cells and B cells and results in protein isoforms with opposite effects on the activity of octamer/TAATGARAT-containing promoters. J Biol Chem, 1992. 267(35): 24960–5.

